# pH-dependent structural dynamics of the neuropeptide Y in aqueous solution

**DOI:** 10.1101/2024.11.04.621936

**Authors:** Hoa Thi Nguyen, Marc Spehr, Ana-Nicoleta Bondar, Paolo Carloni

**Author notes:** Author Contributions The manuscript was written through contributions of all authors. All authors have given approval to the final version of the manuscript.

## Abstract

**ABSTRACT.** The neuropeptide Y regulates key molecular processes in the brain. Its interaction with the cell membrane, where it binds to specialized receptors with key physiological roles, likely depends on the pH. The structural ensembles of the porcine and human peptides, solved by Nuclear Magnetic Resonance (NMR) acid pH in aqueous solution, indicate an *α*-helical core with unstructured termini. However, the protonation states of the carboxylic and histidine residues of the peptide, and the interplay between protonation states and peptide conformational dynamics, have not been explored. Here we perform constant pH simulations and graph-based analyses to investigate dynamics and H-bond patterns of the neuropeptide Y in the pH range from 3.0 to 7.0. We find that an *α*-helical core, as observed in the NMR experiments, is presented at all pH values, though its length can vary by 2-3 residues depending on the pH. The pK_a_ of Asp16, part of the *α*-helix, and of Asp11 may shift by more than on pH unit. Based on these findings, we suggest that performing constant pH simulations may be required to describe accurately the interactions of the peptide with its cellular partners at the pH values of interest.

## INTRODUCTION

The neuropeptide Y (NPY) is a 36-residue peptide (Chart 1) abundant in mammalian brains[1], where it is involved in regulating key processes such as memory formation and feeding[2]. In the hypothalamus, human (h)NPY participates in cell signaling paths that delay aging[3]. In the peripheral nervous system, hNPY is involved in vasoconstriction[4]. The peptide can still be detected in postmortem human brain tissue[5], which allows researchers to study hNPY distribution and concentration in human diseases such as neurodegenerative diseases. hNPY may bind to the cell membrane[6] and/or to its membrane-embedded target receptors [7, 8]. These are the Y1, Y2, Y4, and Y5 receptors, which belong to the G protein Coupled Receptor (GPCR) superfamily[9–11]. hNPY-based peptide conjugates (i) are being designed for cancer diagnosis because the receptors targeted by hNPY are overexpressed in cancer cells[12], which often have a dysregulated pH [13]. They are also used to deliver therapeutic cargo to these cells[14–16]. pH likely affects NPY’s membrane interactions substantially, as suggested by two observations: **(i)** Electrostatic interactions – which drive peptide binding to the membrane[17] – are weaker in a buffer at pH 7.4 than in aqueous solution at pH 7.0[18]. **(ii)** The estimated free energy of peptide binding to a standard membrane is approximately 40% the energy calculated for a corresponding membrane with a high concentration of acidic lipids[19]. These observations raise the key question of as to whether and how the conformational dynamics of NPY depend on pH, which in turn could influence its membrane and receptor binding. To study the conformational dynamics of the peptide as a function of the pH, here we carry out extensive atomic-level simulations using constant pH and standard molecular dynamics simulations. We rely on graph-based approaches to characterize the protonation-dependent dynamics of the peptide.

The three-dimensional folds of NPY and of the two other peptides from the pancreatic polypeptide family, the pancreatic peptide (PP) and peptide YY (PYY), have been studied for decades [20]. The first structure of the PP peptide from turkey (avian, aPP), solved in the early 80’s by X-ray crystallography, revealed the PP fold in this family[21]. It consists of the following regions: (i) an N-terminal region with the polyproline type-II helix (residues 2 to 8) containing the three conserved (see P2, P5, and P8 in Chart I); (ii) an amphipathic α-helical core (residues 14 to 31); (iii) a flexible C-terminal segment[21–23]. The N-terminal segment is back-folded anti-parallel to the *α*-helical segment, such that the peptide assumes a hairpin structure. The three conserved N-terminal Pro residues are on the same side of the polyproline helix[24] and interdigitate with hydrophobic residues of the α-helical core helix– a structural feature suggested to explain why a monomeric PP peptide would be stable in solution[24]. Intra-molecular H-bonds involving charged residues were suggested to help shield the hydrophobic core from the aqueous solution[24].

Fig 1A illustrates the intra-molecular hydrogen (H) - bonds revealed by the X-ray crystal structure of aPP, which was solved at very high (0.99Å) resolution[22]. Tyr7 and Asp10, and Asp16 and Arg19, are within H-bonding distance of 2.7-2.8Å. In the more recently solved X-ray crystal structure of antibody bound human (h)PYY peptide, the amidated C-terminal region of the peptide H-bonds to the Fab antibody. It is thought that such H-bonds might be more generally relevant for the binding of PP peptides to their membrane receptors[20]. Tyr20 and Arg25 of the *α*-helical core are important for the conformational dynamics of NPY: when either or both are mutated to Phe in hNPY, the peptide had higher thermal stability and higher propensity to oligomerize in aqueous solution[25].

**Fig 1.**
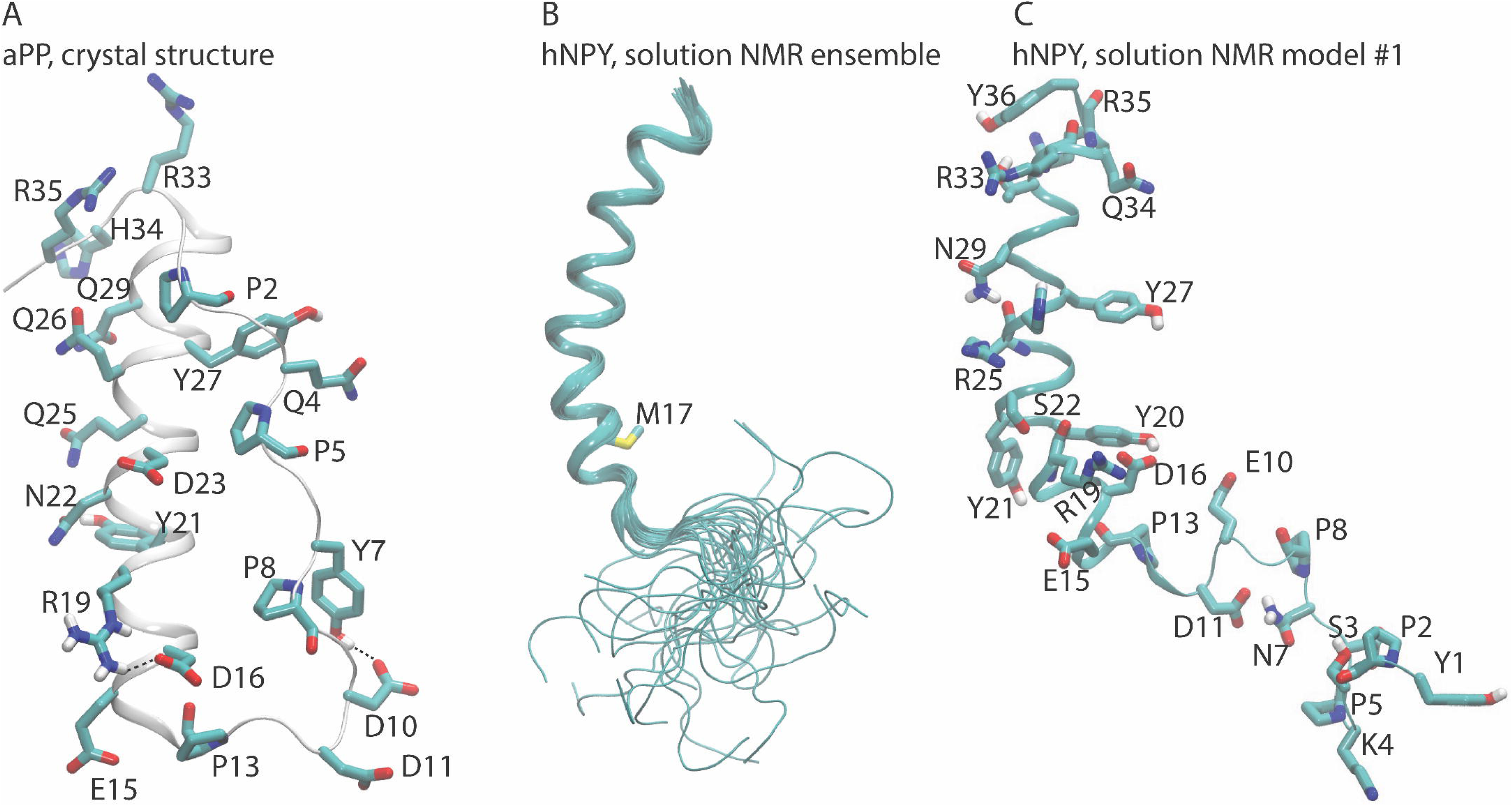
Architecture and Intramolecular H-Bonding of aPP and NPY peptides. (A) The aPP X-ray structure, solved at high resolution, reveals intramolecular H-bonds between Y7 and D10 and between D16 and R19 (PDBID 2BF9). This panel was created using Visual Molecular Dynamics (VMD)[32]. For clarity, only selected H atoms and the backbone of the proline residues are shown. (B) The NMR structural ensemble (26 conformations) of hNPY in aqueous solution (PDB ID 1RON). Note that the N-terminal region is flexible and lacks the tight packing against the α-helical core observed in the aPP crystal structure from panel A. M17 distinguishes hNPY from pNPY (Chart 1), which was investigated in this work. (C) Model #1 from the NMR ensemble with selected residues. In this model, the distances are as follows: S22-Oγ and the R19 backbone carbonyl group, 2.8Å; N29-Nδ2 and the R25 carbonyl, 3.2Å; S3-Oγ and the P5 carbonyl, 4.6Å; the side chains of N7 (Nδ2) and D11(Oδ1), 2.8Å; D16 (Oδ2) and Y20 (O), 2.8Å; and T32 (Oγ) and Y36 (OH), 3.4Å, D6 (Oδ1) and P5(O), 4.8Å.

The NMR structure of hNPY in solution at pH 3.2 and 310K indicates an α-helical core Pro13 to Thr32[26] (Fig 1B), a highly flexible N-terminal region. There is no close packing between the latter and the α-helical core, unlike the crystal structure of pYY (Figs 1A and 1B). As was initially observed for pYY, the NMR structure of hNPY reveals intramolecular H-bond distances between side chains — specifically, between Asp16 and Tyr20 and between Arg35 and Tyr36 (Fig 1C). Additionally, the ensemble of NMR structures of hNPY also indicates that intramolecular H-bonding between carboxylic sidechains and backbone carbonyl groups can be sampled at acidic pH - in one of the structural models of the ensemble (#26), the carboxyl group of Asp6 is within 3.0-3.1Å of the backbone carbonyl groups of Lys4 and Pro5, and the carboxyl group of Asp11 is within 2.8Å distance of the backbone carbonyl group of Pro8. Such short distances between a carboxylic group and a backbone carbonyl group could suggest that the former is protonated.

Given the important role of pH in NPY’s interactions with cellular partners and membranes, we study here the conformational dynamics of the peptide as a function of the pH by constant pH molecular dynamics simulations [27, 28]. These allow the protonation states of ionizable groups to respond to changes in the chemical environment and the external pH. This contrasts with standard MD, which uses fixed protonated states for ionizable side chains and it may provide limited information on peptide conformational dynamics when pK_a_ values of residues are near the chosen pH (as is the case for histidine at physiological pH). For these residues, both protonated and deprotonated states can coexist, and their relative amounts can change due to conformational rearrangement. Our study is complemented by graph-based approaches to characterize the protonation-dependent dynamics of the peptide.

In light of our long-term goal of studying peptide/membrane binding, we chose porcine (p)NPY as our model system. This decision was made for the following reasons: (i) a solution NMR structure of pNPY at pH 3.1 revealed very similar features[29]; (ii) hNPY and pNPY differ only for the M17L substitution[26] (see Chart 1 and Fig 1B) and (iii) the structure of pNPY bound to micelles has been determined[30].

**Chart 1**. The sequence of hNPY and pNPY sequences. We consider residues 29 to 64 (without the signal peptide and C-terminal residues 65 to 97 of the full length NPY, UniProt entry A4D158), labeled 1 starting from the N-terminal Tyr, such that Y1 corresponds to Y29 in full length sequence of NPY[31]. The C-terminus is amidated[26]. Note that the sequence of hNPY and pNPY are identical except for position 17 (Met in hNPY, Leu in pNPY). Asp and Glu residues, Arg and Lys, the histidine are colored red, blue, and green, respectively. The underlined residues belong to the helical segment of the peptide.

*h(p)NPY: Y_1_PS**K**_4_P**D**_6_NPG**E**_10_**D**_11_APA**E**_15_**D**_16_ **M**_17_**(L)**A**R**_19_YYSAL**R**_25_**H**_26_YINLIT**R**_33_Q**R**_35_Y_36_*

## METHODS

Our calculations are based on the hNPY NMR structural ensemble (model #1) in solution (PDB ID 1RON) [26]. The pNPY structure was obtained by replacing Met17 with a Leu residue. The Phyre2 web portal was used to model the mutant peptide[33]. For the equilibration phase by standard MD simulations, His26 was protonated in Nδ tautomer and the Glu and Asp residues (Chart 1) were considered negatively charged. For the constant pH simulations, the protonation of these residues is dictated by the algorithm. The peptide was placed in the center of coordinates of a box of 14,125 water molecules of 77 x 77 x 77 Å^3^ with 0.15 M neutralizing NaCl using the CHARMM-GUI protocol[34].

### Force-fields

For the standard MD simulations, we used the CHARMM36m force field parameters for the peptide and ions[35–40], and the TIP3P water model[41]. This combination of force fields is recommended for this approach[35, 42, 43]. For the CpHMD simulations we used the recommended protocol[44] whereby the protein is described with the CHARMM22/CMAP force field [38, 45], water molecules, with the CHARMM-modified TIP3P model[46], and ions, with CHARMM36[40]. Thus, a different water model was used in the two types of simulations in order to comply with the recommended protocols.

### Standard MD simulation

We used the SHAKE algorithm[47] to constrain all covalent bonds involving H atoms, and an integration timestep of 2fs. Following geometry optimization of the peptide using conjugate gradient[48], we performed a short, 10ns equilibration in the *NVT* ensemble (constant number of atoms *N*, constant volume *V*, and constant temperature *T*) with harmonic constraints of 1.0 kcal/mol/Å^2^ on the peptide hetero-atoms. We used for this phase the Langevin thermostat[49] to keep constant temperature conditions. The damping coefficient was 5.0 ps^-1^, and the target temperature was set at 303.15K. Hydrogen atoms were kept uncoupled from the thermostat to avoid noise. This reduces artifacts because of their fast movements[50]. We then switched off all constrains and performed simulations at constant temperature (T=303.15K), with same setup as above, and constant pressure (P=1atm) using the Langevin piston (Nose-Hoover Langevin barostat)[51]. The target pressure was set at 1.01325 bar (1atm), with an oscillation period of 50 fs. Long-range electrostatics interactions were calculated using the Particle Mesh Ewald (PME)[52], using 1Å of grid spacing and spline order of 6. The cutoff radius of van der Waals interaction and the real part of the electrostatics was set to 12 Å. We used the same starting coordinates to perform three *NPT* production runs, 500 ns-long each, with NAMD[53] package. The simulations differ for their Maxwell-Boltzmann distribution of the velocities.

### CpHMD simulations

The initial structure was a snapshot after 465ns of the standard MD simulation above. During the dynamics, the charge of a titratable residue might change. Because the Ewald summation requires the simulation box to remain neutral[54], we used the protocol by Wallace et al[55], whereby the protonation titration is coupled with the simultaneous ionization or neutralization of a dummy co-ion in solution.

The CpHMD simulations utilised the Nose-Hoover thermostat[56] and barostat[57] to maintain constant temperature and pressure, respectively. In the thermostat, the target temperature was set at 303.15K, and the mass of the thermostat was set at 1000 kcal.ps^2^. In the barostat, the reference pressure was set at 1 atm, the mass of piston of 500 amus. The friction coefficient for the piston was set to 20 ps^-1^. The temperature of the bath was maintained at 303.15K. The SHAKE algorithm[47] was used to constrain all covalent bonds involving hydrogen (H) atoms. We then switched off all constrains and performed simulations at constant temperature (T=303.15K) and pressure (P=1atm), with same setup. Following the recommend minimization procedure for CpHMD simulations in CHARMM[44], we performed a geometry optimization for the peptide using 10,000 steps of Steepest Descents (SD) and 5,000 steps of an adopted Newton-Raphson method (ABNR). Likewise, following the recommended protocol, we equilibrated pNPY in water using 3 stages as follows: **(i)** heating by 0.2 ns restrained MD from 5 K to 303.15 K, in 100,000 steps with harmonic force constant of 1.0 kcal/mol/Å^2^. **(ii)** Four CpHMD equilibration steps, 0.2 ns each, the first two in the *NVT* ensemble, then two in the *NPT* ensemble, with the peptide hetero-atoms restrained with harmonic force constant of 1.0; 0.5; 0.1, and 0 kcal/mol/Å^2^. **(iii)** CpHMD production run without any constraint.

For each pH value, we used the equilibrated structure to perform three independent CpHMD simulations. We name these simulations as replica R#1 (50ns), R#2 (50ns), and R#3(40ns).

### Predictions of pKa’s values of titratable residues

The titration coordinate *λ* takes values *λ* = 1 for the deprotonated state, and *λ* = 0 for the protonated state[27]. Since histidine and carboxyl sidechains each contain two titration sites (denoted as tautomeric states), an additional coordinate, *χ*, is used to describe the interconversion between the tautomeric states. The numbers of deprotonated (*N_deprot,i_*) and protonated states (*N_prot,i_*) for a residue *i* are then defined as:

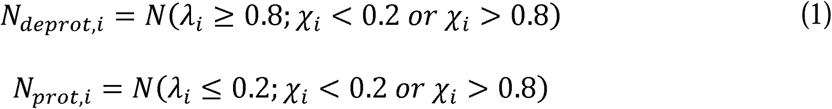

The mixed tautomeric states with 0.2 < χ*_i_* < 0.8 are discarded. We use cut-off values *λ*_i_*ε*[0.8,1.0] for deprotonated states, and *λ*_*i*_*ε* [0.0,0.2]for protonated states. The charge of atom a on a titrating residue *i* reads:

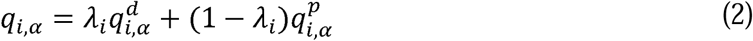

where 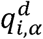 and 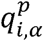 are the charges in the deprotonated and protonated states, respectively.

For a trajectory at a specific pH, and for a residue *i* (in this study, *i* = 1 for Asp6, *i* = 2 for Glu10, *i* = 3 for Asp11, *i* = 4 for Glu15, *i* = 5 for Asp16, and *i* = 6 for His26), the number *N_prot,i_* counts the times in which a titratable residue *i* is in the protonated (*λ_i_* ≤0.2) states, while *N_deprot,i_* counts the times of deprotonated (*λ_i_* ≥ 0.8) states. The deprotonated fraction of residue *S*_i_ can be calculated from the simulations at different pH as:

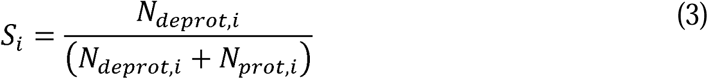

where *N_deprot,i_* and *N_prot,i_* are number of deprotonated and protonated states, respectively. The *titration curve* plots *S_i_*(*pH*) as a function of the pH, is obtained by fitting the *S_i_* values across all simulated pH values to the generalized Henderson-Hasselbalch (HH) equation,

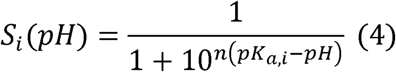

where *n* is the Hill coefficient.

The *pK_a,i_* value of residue *i* is given by the pH value at which *S_i_* = 0.5.

*H-bond network analyses.* To study how the internal H-bond network of the peptide as a function of the pH, we used the graph-based algorithm and graphical user interface Bridge/Bridge2[58, 59]. With this approach, an H-bond graph consists of nodes, i.e., the H-bonding protein groups included in the graph computation, and edges, i.e., the H-bond connections (direct or water mediated) between these NPY groups. We used the last 2,000 equally spaced coordinate snapshots of 40 ns of each of the CpHMD labeled as R#1 and R#2 and the last 1,500 equally snapshots of the 30ns CpHMD labeled as R#3. The Bridge computations were performed for the 3 replicas at once by uploading all trajectories for a given pH. To understand how the sidechains, backbone groups, and water-mediated bridges, contribute to the intra-molecular H-bond network of pNPY, we separately computed three sets of H-bond graphs:

1. direct H-bonds of sidechains and backbone groups, and water-mediated bridges of these groups;
2. direct sidechain-sidechain H-bonds and water-mediated bridges between sidechains;
3. direct H-bonds between the sidechains.

We considered water bridges between H-bond donor and H-bond acceptor formed by one, two or three H-bonded waters. To identify H-bonds we used standard geometric criteria: distance between the donor and acceptor heteroatoms ≤ 3.5 Å; H-bond angle lower than either 20° or 60°. Results for both cutoff values are reported. The *occupancy* of an H-bond is given by the percentage of coordinate sets, of the total number of coordinate sets used for analyses, in which the H-bond criteria are met. Here, we consider only H-bonds with an occupancy equal or higher than minimum thresholds reported in Table 1.

**Table 1.** H-bond criteria and occupancy thresholds used for the H-bond graphs.

| Set of H-bond graphs | H-bond angle criterion of $20^\circ$ | H-bond angle criterion of $60^\circ$ |
| --- | --- | --- |
| 1) | 10% | 50% |
| 2) | 10% | 25% |
| 3) | 10% | 15% |

Notice that Bridge does not distinguish between the H-bonds that a protein group donates or accepts.

### Secondary structure elements

These were determined using STRIDE (TIMELINE analysis) within Visual Molecular Dynamics (VMD)[32] graphics software.

## RESULTS

### 1. pK_a_ Predictions

S1-36 Figs show the evolution of coordinates, partial charges and the cumulative deprotonated fraction values of each titratable residues *i* of the peptide (*i* = 1 for Asp6, *i* = 2 for Glu10, *i* = 3 for Asp11, *i* = 4 for Glu15, *i* = 5 for Asp16, and *i* = 6 for His26), informing the convergence of the protonated states. For **Asp6** (S1-S6 Figs), at pH = 7, 6 the *λ*_1_values across the three simulation replicas are close to 0.8-1.0 (S1 and S2 Figs). The fraction of deprotonated residue *S*_1_ is close to 1 (S6 Fig), hence this residue is fully deprotonated. At pH = 5, 4, the *λ*_1_values are found both in the range of 0.8 - 1.0 (more at pH 5 than pH 4), and in the range of 0.0 - 0.2 (more at pH 4 than pH 5). As a result, Asp6 is mostly deprotonated at pH = 5 with *S*_1_∼0.8 (S3 and S6 Figs) and it is mostly protonated at pH = 4 with 0.2 ≤ *S*_1_ ≤ 0.4 (S4 and S6 Figs). At pH = 3, the *λ*_1_ values are primarily distributed in the range of 0-0.2 and *S*_1_∼0.1, indicating that Asp6 is fully protonated (S5 and S6 Figs).

**Glu10** is deprotonated at pH values 7, 6, and 5 (*S*_2_∼1.0, S7-S9 Figs). At pH = 4 and 3, it is mostly deprotonated or protonated (*S*_2_∼0.6 - 0.8, and *S*_2_∼0.2 - 0.4, respectively, S10-S12 Figs). **Asp11** is deprotonated at pH = 7 and 6 (*S*_3_∼1.0 and *S*_3_∼0.8, respectively, S13, S14 and S18 Figs). At pH = 5, it is mostly protonated (*S*_3_∼0.4, S15 and S18 Figs). At pH = 4 and 3, it is protonated (*S*_3_∼0.1 and S_3_∼0.0, respectively, S16-S18 Figs).

**Glu15** is fully deprotonated at pH = 7, 6, 5 (*S*_4_∼1.0, S19 - S21 and S24 Figs) while at pH = 4 it is mostly deprotonated (*S*_4_∼0.6). It is protonated at pH = 3 (*S*_4_∼0.2, S22-S24 Figs). **Asp16** is deprotonated at pH = 7 - 4 (*S*_5_∼0.8 - 1.0, S25-S28 and S30 Figs), and it can be either deprotonated or protonated at pH = 3 (*S*_5_∼0.4 - 0.6, S29-S30 Figs). **His26** is mostly neutral at pH 7 (*S*_6_∼0.8, S31 and S36 Figs). At pH = 6, it is both mostly positively charged and neutral (*S*_6_∼0.4, S32 and S36 Figs). At lower it is totally protonated (*S*_6_∼0.0 - 0.1) (S33-36 Figs).

The pK_a_ values, estimated by the titration curves (S37-S38 Figs, S1 Table), turned out to be converged within 10ns of constant pH dynamics. The largest shifts in pK_a_ (ΔpK_a,i_) of residues *i* (X_i_) relative to its “standard” value in the CH3CO-AAXAA-NH2[44] peptide are observed for Asp16 (ΔpK_a,5_ < -1) and Asp11 (ΔpK_a,3_ > 1) (see the Fig 1 and S1 Table). The negative pK_a_ shift of Asp16 might be caused, at least in part, by its salt bridge with Arg19 (Fig 3A and S39-S63 Figs). At pH = 5 or lower, Asp11 interacts with Glu10, that is mostly deprotonated, as seen in the above analysis. The ionized state of the residue may be stabilized by its salt bridge with Arg25 (Fig 3B and S55 Fig). The Asp11-Glu10 interaction may cause, at least in part, by the large positive shift in pK_a_.

**Fig 2.**
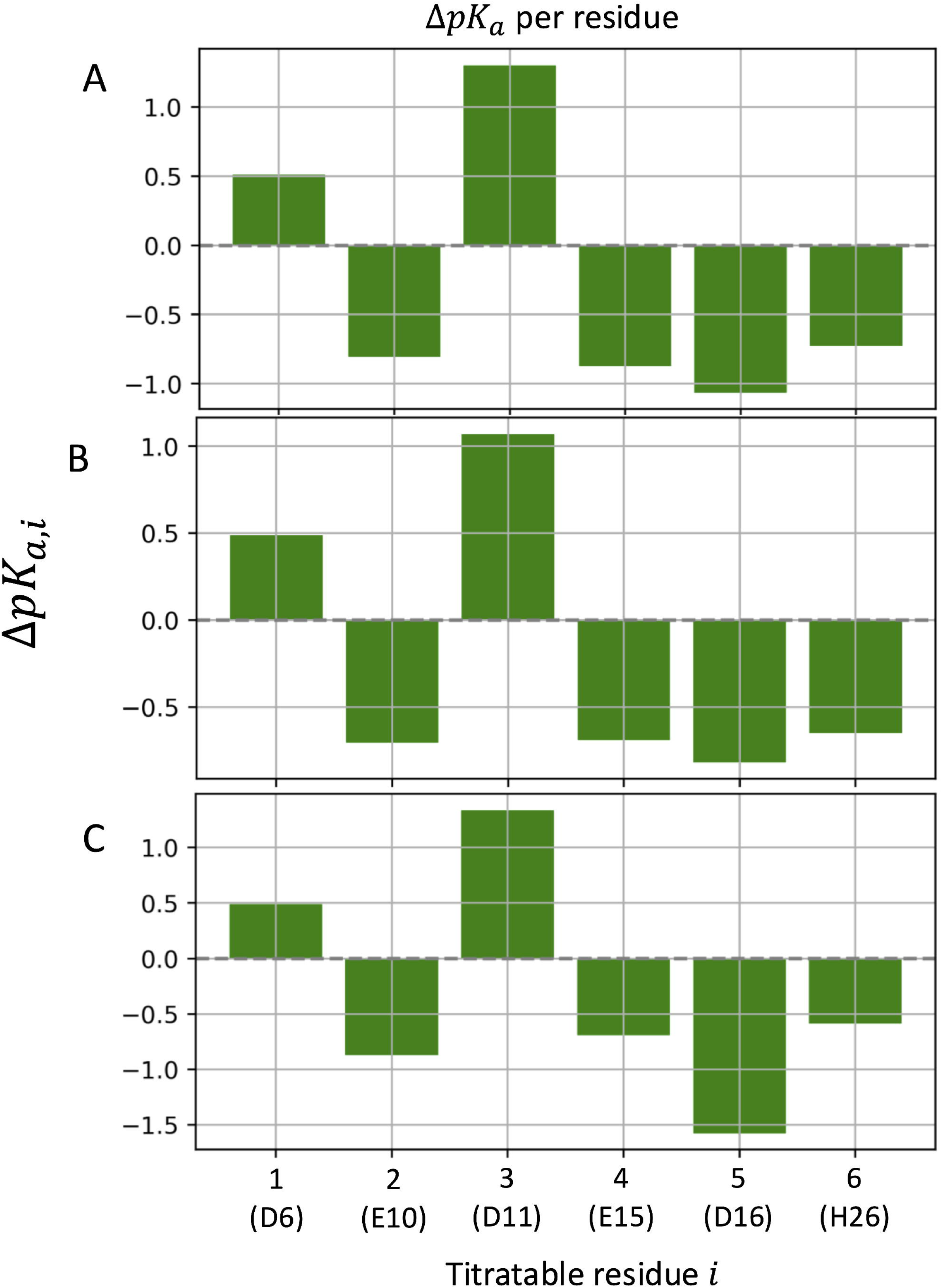
ΔpK_a,i_ **values of the titratable residues**. (A-C) refer to simulations R#1-3.

**Fig 3.**
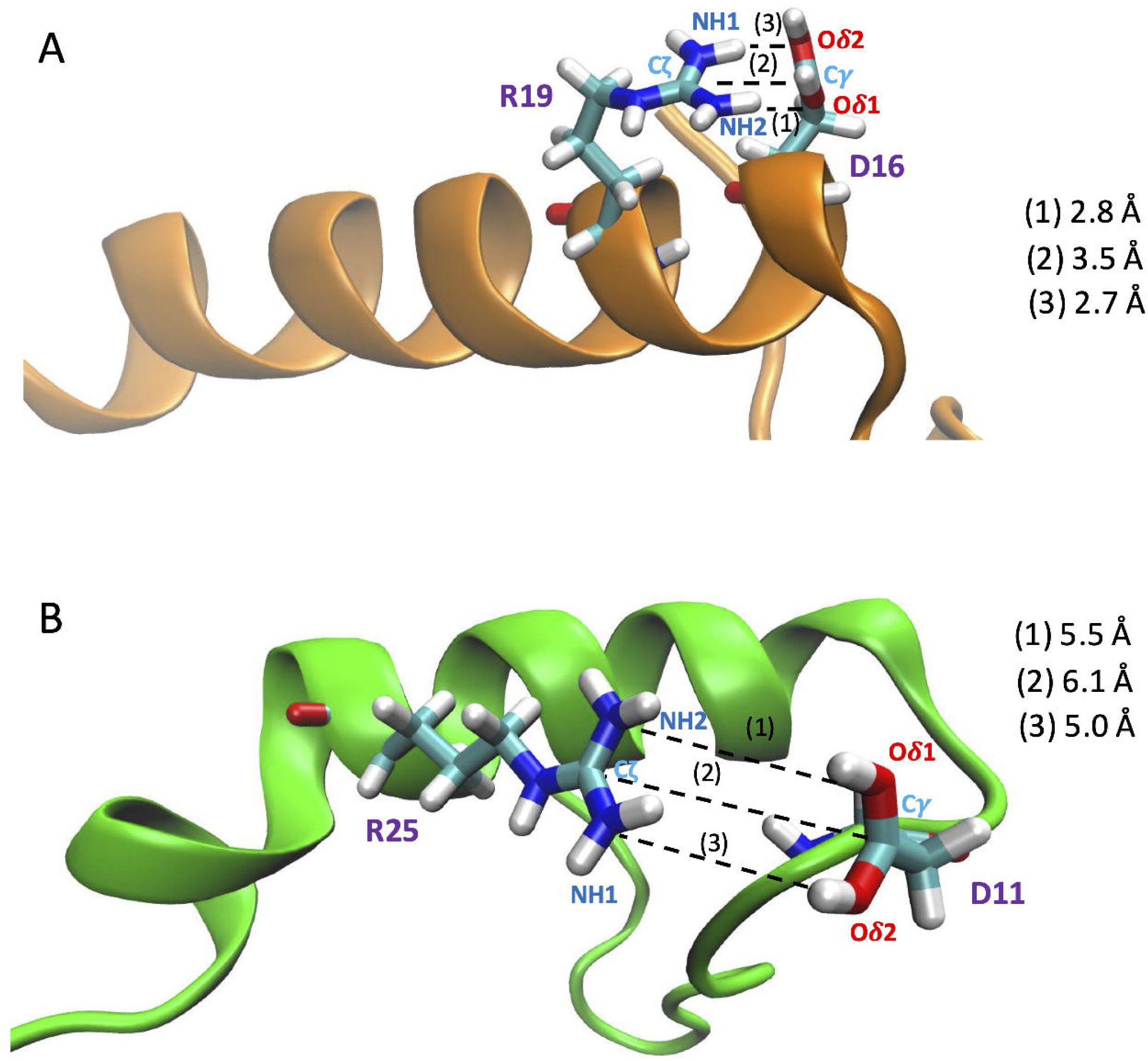
Asp16 - Arg19 (A) and Asp11 – Arg25 (B) salt bridge from our CpHMD simulations at pH 7 (R#1, 2, respectively). In these snapshots (at 91.8 ns and 135.0 ns, respectively), the (1) Oo1 (Asp) – NH2(Arg), (2) Cy(Asp) - C((Arg), and (3) Oo2(Asp) – NH1(Arg) distances are the shortest. The first salt bridge is also observed in R#2 and R#3, whereas the second only in R#2 at pH 7, 6.

### 2. H-bond networks

We used the Bridge/Bridge2 graph-based algorithm and graphical user interface[58, 59] to identify H-bond network at different pH investigated here (Figs 4-6, S39-S63). We performed separate computations for H-bond networks for protein groups (sidechains and backbone) focus on either direct or water mediated H-bonds (up to three water molecules, Fig 4), side chain-side chain on either direct or water mediated H-bonds (Fig 5), and for side chain-side chain on direct - bonds (Fig 6). All graph computations were performed with a distance H-bond criterion of 3.5Å. To test how the choice of the H-bond angle (between acceptor, hydrogen atom and donor) influences the results used two different angle criteria, 20^0^ and 60^0^. As expected, using the stricter 20^0^ H-bond angle criterion reduces the number of H-bonds in the graphs (Figs 4-6B).

**Fig 4.**
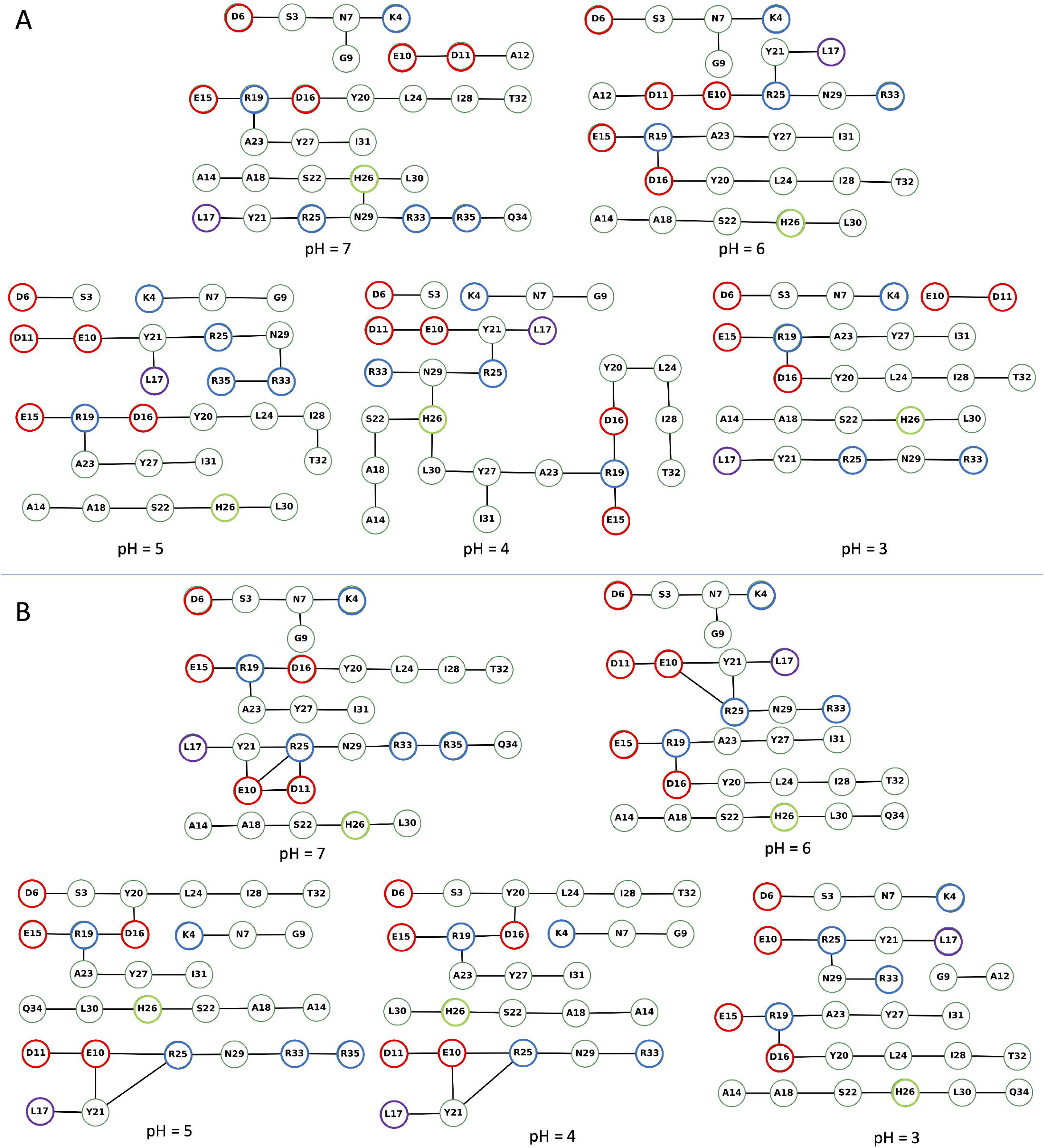
H-bond graphs computed for the backbone, sidechains, and three water bridges at different pH values. We used the complete dataset of three replicas at each simulated pH value to compute the H-bond graphs with Bridge2[59]. All graph computations used an H-bond distance criterion of 3.6Å. The color coding of the nodes is as follows: red, Asp and Glu residues, blue, Arg and Lys, violet, Leu17, lime, His26, and green, all other residues. (**A-B**) H-bond graphs computed with an H-bond angle criterion of 60^0^ (panel A) vs. 20^0^ (panel B). The H-bond occupancy threshold used, 50% and 10% for the angle criterion of 60^0^ and 20^0^, respectively. Note that the nodes and edges of the graph were manually arranged in the Bridge2 graphical interface to minimize the space taken by each panel; thus, nodes representing the residues of the peptide are not arranged according to their relative location in the peptide.

**Fig 5.**
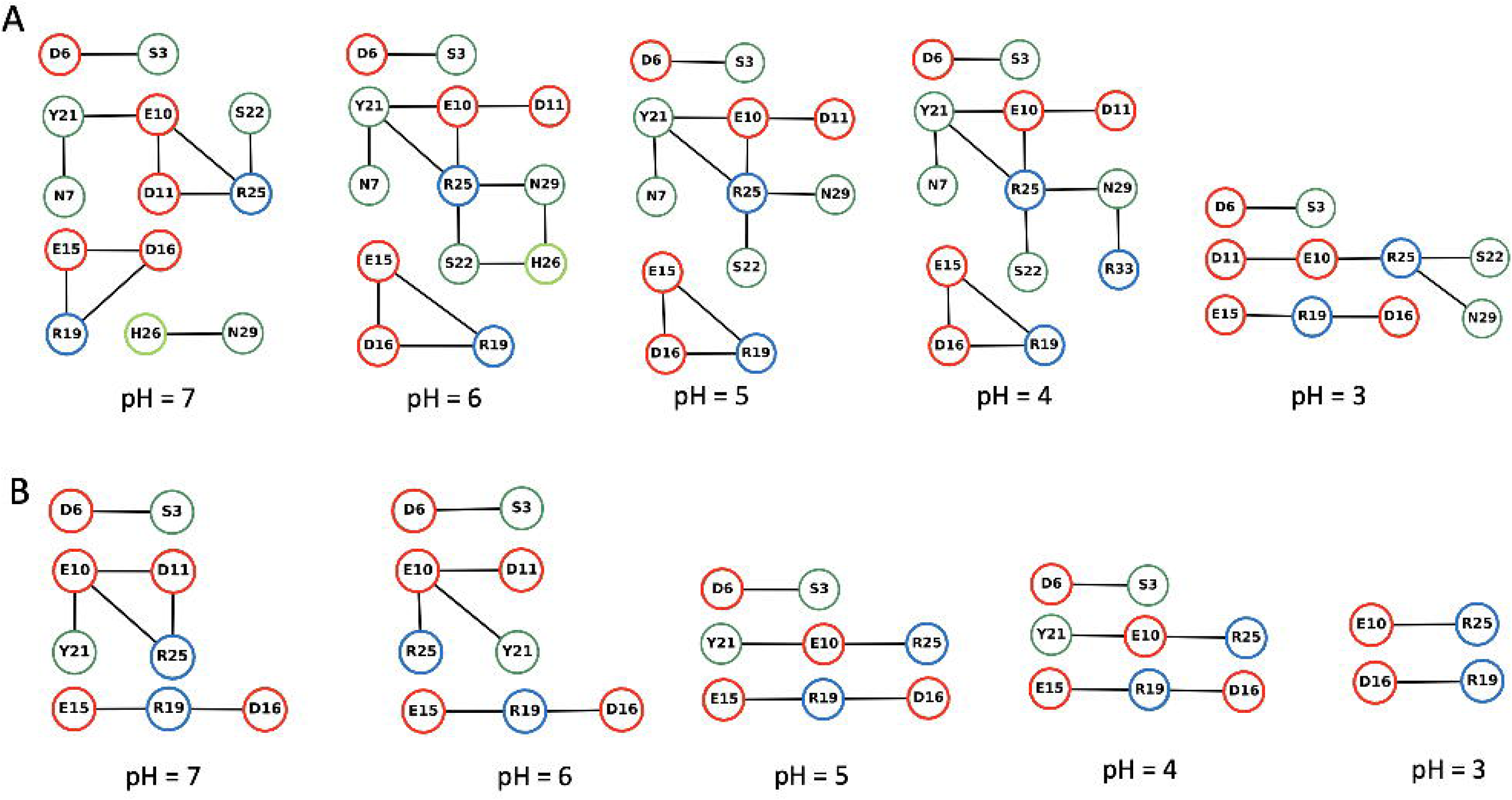
H-bond graphs for direct sidechain-sidechain H-bonds and water-mediated bridges, without the protein backbone groups. We used the complete dataset of all replica simulations at each pH value. (A, B) H-bond graphs with an H-bond angle criterion of 60^0^ in (A) and 20^0^ (B). The H-bond occupancy threshold used, 25% and 10% for the angle criterion of 60^0^ and 20^0^, respectively.

**Fig 6.**
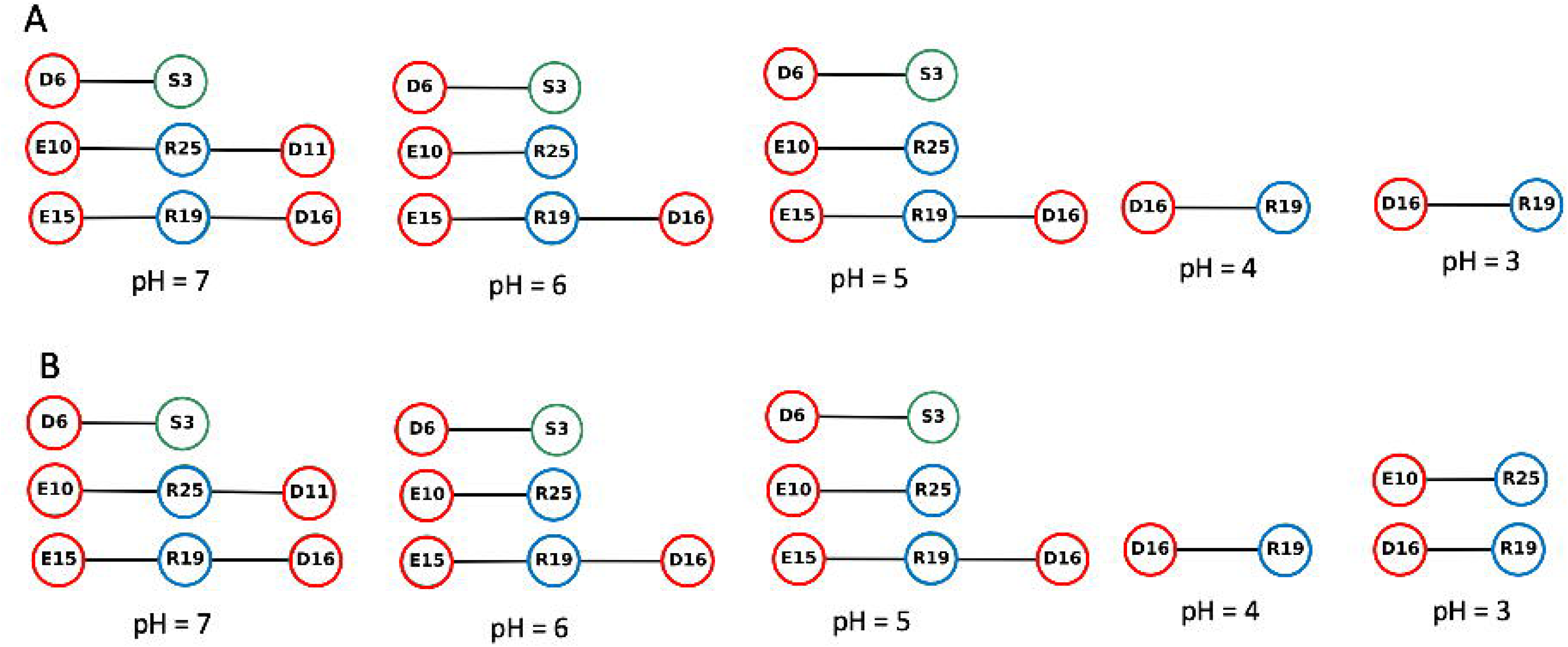
Same as Fig.4, but only for direct H-bonds of sidechains. The H-bond occupancy threshold used, 15% and 10% for the angle criterion of 60^0^ and 20^0^, respectively.

The number of direct H-bonds (Fig 6A) is lower than that of water-mediated H-bonds, whether one considers only sidechains (Fig. 5A) or sidechains and protein backbone groups (Fig. 4A). Interestingly, at all pH values and with either H-bond angle criterion, Asp16 side chain H-bonds to Arg19 side chain at all pH values (Fig 6A), while the side chains of Asp6, of Glu10, and of Glu15 form direct H-bonds with Ser3, Arg25, Arg19 at pH 7-5 (Fig 6A).

H-bonds and water-mediated bridges between sidechains are as follows. **Asp6** samples water-mediated H-bonds with Ser3 at all pH values, and direct H-bonds at pH 7-5 (Figs. 5A, 6A). At all pH values, the **Glu10** has direct or water mediated H-bonds with Arg25 and water-mediated H-bonds with Asp11 (Fig. 5A). **Asp11** has direct or water mediated H-bonds with Arg25 at pH 7 (Fig. 5A). **Glu15** has direct or water-mediated H-bonds with Arg19 at all pH values (Fig. 5A), and water- mediated H-bonds with Asp16 at pH 7-4 (Fig. 5A). **Asp16** connects to Arg19 side chain at all pH values (Fig. 5A).

### 3. Conformation dynamics

Our model of pNPY in solution is obtained from the hNPY NMR structure at pH 3.2 and a temperature of 310K[26]. Its a-helix extends from residue Asp11 to Tyr36[26]. It is mostly maintained during the equilibration phase (by 500-ns long MD simulations at 303.15K, see S64 Fig, S5 Table) and during the constant pH simulations at 303.15K (S65-S69 Figs). However, residues Arg33-Tyr36 at the C-term and Asp11-Ala14 at the N-term lose their secondary structure (S65-S69 Figs). Thus, the helix becomes shorter. A decrease of a -helical content relative to the human one has been observed also in NMR studies of pNPY in solution[29]: at pH 3.2 (temperature not specified in the study: the helix extends from Pro13 to Tyr36.

The conformational fluctuations of the peptide are quantified by PAD[60] values (Fig 7). The higher the value for a backbone unit, the larger its fluctuation. Overall, as expected, residues at the N- and C-termini (mostly coils) have larger fluctuations than the helix at all values of pH, and even more so at higher and low pH values (S70-S73 Figs). An unanticipated finding is that only a smaller fragment of the peptide, from Glu15 to Arg25, has very small PAD values that are similar to those of an α-helical segment of a folded protein, as seen in refs[61–65].

**Fig 7.**
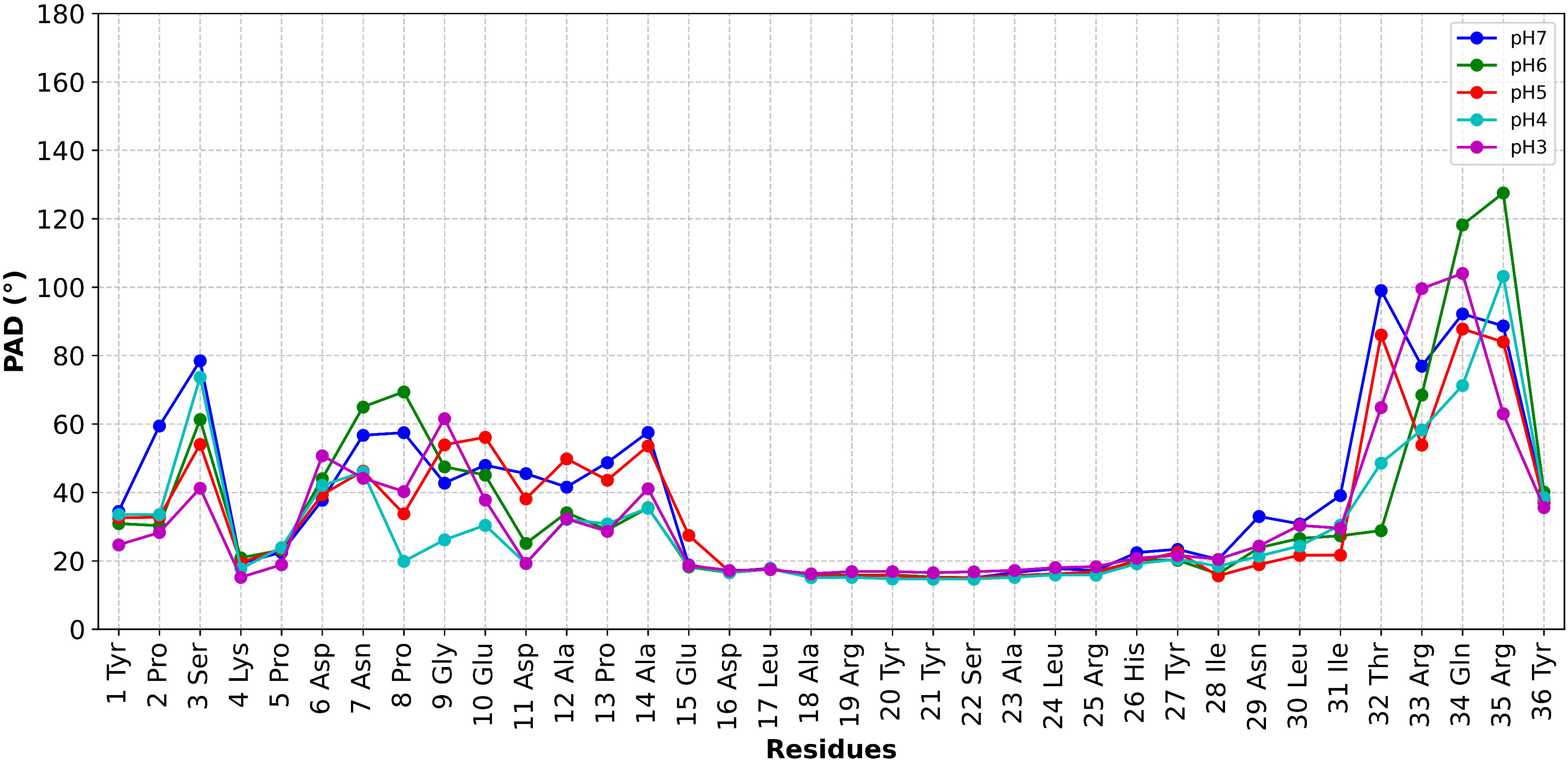
The backbone fluctuations during the CpHMD simulations as quantified by PAD values. The larger the PAD values, the larger the fluctuations of the residue backbone atoms[60]. The central helical core has very similarly low PAD values, indicating similar rigidity, whereas the N- and C-termini have larger fluctuations with distinct PAD values in each of the three replicas.

## DISCUSSIONS AND CONCLUSIONS

We have carried out constant pH simulations to study the protonation patterns of the pNPY peptide in aqueous solution, upon acidification, from pH 7 to 3. Three independent simulations, for a total of 110 ns, indicate that the peptide keeps most but not all of its long a-helical structure present at pH 3.1 and pH 7.4 for the same peptide, as observed by NMR[29], while few residues at both the N-terminus and the C-terminus side assume a random coil conformations (S65-S69 Figs). As expected, the helix is relatively rigid at all pH values (Fig 7). The termini are disordered, as observed experimentally in both the human[26] and porcine[29] NMR structural ensembles.

The H-bond graph computations indicate that the intra-molecular H-bond network of the peptide is strongly pH dependent. The H-bond graph computations indicate that the intra-molecular H-bond network of the peptide is strongly pH dependent. As anticipated from the NMR structure, there are relatively few H-bonds directly between the sidechains, and Asp16 and Arg19 directly connect to each other at each pH value (Fig 6). Asp6 and Glu15 lack direct sidechain H-bonds at pH = 3 and pH = 4. H-bonds between Asp carboxyl groups and Pro backbone carbonyl groups are instead present only in the hNPY NMR ensemble.

The pKa values of the six ionizable residues of the peptide estimated from their calculated titration curves in aqueous solution (S37-S38 Figs) converged within about 10ns. Two residues experience rather large changes in pKa (more than 1 pKa unit) relative to their standard values, as assumed to be in the CH3CO-AAXAA-NH2[44] peptide, where X is the residue we focus on[44]. Asp16 has low pKa, possibly because of its close interactions with Arg19 (Fig 3A). As a result, Asp 16 is always deprotonated except at pH = 3. The opposite effect is observed for Asp11, which is adjacent to Glu10, and which Asp 11 is protonated even at mildly acidic solutions. The only histidine residue of the peptide. His26, is mostly neutral only at pH 7, and is protonated at lower pH values.

The hNMR structure (model#1) shares one intermolecular H-bonds involving side chain present in our simulation of pNPY (Glu10 (N) with Asp11 (O*δ*1)). However, as expected, several other interactions are only present in the simulated structural ensemble (including Asp16 (O*δ*1) with Arg19 (NH1), Asp6 (N) with Ser3 (Oγ), Asp6 (N) with Ser3 (Oγ), Glu15 (O*ε*1) with Arg19 (N)) and only in the NMR structure (Asp11 (O*δ*1) with Asn7(N*δ*2), Asp16(O*δ*2) with Tyr20(OH)).

In conclusion, we have presented here a CpHMD study on NPY in neutral and acidic solutions, ranging from pH 7.0 to pH 3.0. The pK_a_ values of Asp residues and the protonation states of ionizable residues vary significantly upon decrease of pH. We suggest that performing CpHMD simulations might prove critical to accurately describe NPY binding with its cellular partners (including the Y1R/Y2R receptors and cellular membranes) at physiological and acidic pH.

## ASSOCIATED CONTENT

### Data Availability Statement

All simulations and data analyses were performed using openly available topology, parameter, and software. The NAMD 2.14 package (https://www.ks.uiuc.edu/Research/namd/) was used to run MD simulations and CHARMM 15.3.2022 package (https://academiccharmm.org) was used to run constant pHMD simulations for NPY in water. Plots were make using Gnuplot package (http://www.gnuplot.info), and Bridge2 program (https://github.com/maltesie/bridge2). Molecular structures were visualized using VMD (https://www.ks.uiuc.edu/Research/vmd/). Data, including input and parameter files for Molecular Dynamics simulations and Constant pH Molecular Dynamics simulations can be found at Zenodo repository: https://doi.org/10.5281/zenodo.14069542.

## Funding Sources

RTG 2416: MultiSenses-MultiScales: Novel approaches to decipher neural processing in multisensory integration Deutsche Forschungsgemeinschaft (DFG) - Project number 368482240

### Notes

The authors declare no competing financial interest.

## ACKNOWLEDGMENT

Hoa Thi Nguyen was supported by DFG (RTG 2416). Open Access publication funded by the Deutsche Forschungsgemeinschaft (DFG, German Research Foundation) – 491111487. The authors gratefully acknowledge the Juelich Supercomputing Centre for providing computing time.

## Supporting Information

1. Prediction of the pKa values of titratable residues.

Here, we present our calculations of the protonation coordinate *i*, partial charge, and deprotonated fraction S*_i_* of residue *i*, *i* = 1 for Asp6, *i* = 2 for Glu10, *i* = 3 for Asp11, *i* = 4 for Glu15, *i* = 5 for Asp16, and *i* = 6 for His26.

**S1 Fig**. **The** 1_1_ **coordinate and partial charges of Asp6 during simulated time at pH 7.** (A) Replica 1 (R#1), (B) Replica 2 (R#2) and (C) Replica 3 (R#3).

**S2 Fig**. Same as **S1 Fig**, but here calculations are carried out for Asp6 at pH 6.

**S3 Fig**. Same as **S1 Fig**, but here calculations are carried out for Asp6 at pH 5. **S4 Fig**. Same as **S1 Fig**, but here calculations are carried out for Asp6 at pH 4. **S5 Fig**. Same as **S1 Fig**, but here calculations are carried out for Asp6 at pH 3.

**S6 Fig**. **Time series of the deprotonated fraction values S of Asp6 at different pH values. S7 Fig**. Same as **S1 Fig**, but here calculations are carried out for Glu10 at pH 7.

**S8 Fig**. Same as **S1 Fig**, but here calculations are carried out for Glu10 at pH 6. **S9 Fig**. Same as **S1 Fig**, but here calculations are carried out for Glu10 at pH 5. **S10 Fig**. Same as **S1 Fig**, but here calculations are carried out for Glu10 at pH 4. **S11 Fig**. Same as **S1 Fig**, but here calculations are carried out for Glu10 at pH 3. **S12 Fig**. Same as **S6 Fig** but calculated for Glu10.

**S13 Fig**. Same as **S1 Fig**, but here calculations are carried out for Asp11 at pH 7. **S14 Fig**. Same as **S1 Fig**, but here calculations are carried out for Asp11 at pH 6. **S15 Fig**. Same as **S1 Fig**, but here calculations are carried out for Asp11 at pH 5. **S16 Fig**. Same as **S1 Fig**, but here calculations are carried out for Asp11 at pH 4. **S17 Fig**. Same as **S1 Fig**, but here calculations are carried out for Asp11 at pH 3. **S18 Fig.** Same as **S6 Fig** but calculated for Asp11.

**S19 Fig**. Same as **S1 Fig**, but here calculations are carried out for Glu15 at pH 7.

**S20 Fig**. Same as **S1 Fig**, but here calculations are carried out for Glu15 at pH 6.

**S21 Fig**. Same as **S1 Fig**, but here calculations are carried out for Glu15 at pH 5. **S22 Fig**. Same as **S1 Fig**, but here calculations are carried out for Glu15 at pH 4. **S23 Fig**. Same as **S1 Fig**, but here calculations are carried out for Glu15 at pH 3. **S24 Fig**. Same as **S6 Fig** but calculated for Glu15.

**S25 Fig**. Same as **S1 Fig**, but here calculations are carried out for Asp16 at pH 7. **S26 Fig**. Same as **S1 Fig**, but here calculations are carried out for Asp16 at pH 6. **S27 Fig**. Same as **S1 Fig**, but here calculations are carried out for Asp16 at pH 5. **S28 Fig**. Same as **S1 Fig**, but here calculations are carried out for Asp16 at pH 4. **S29 Fig**. Same as **S1 Fig**, but here calculations are carried out for Asp16 at pH 3. **S30 Fig**. Same as **S6 Fig** but calculated for Asp16.

**S31 Fig**. Same as **S1 Fig**, but here calculations are carried out for His26 at pH 7. **S32 Fig**. Same as **S1 Fig**, but here calculations are carried out for His26 at pH 6. **S33 Fig**. Same as **S1 Fig**, but here calculations are carried out for His26 at pH 5. **S34 Fig**. Same as **S1 Fig**, but here calculations are carried out for His26 at pH 4. **S35 Fig**. Same as **S1 Fig**, but here calculations are carried out for His26 at pH 3. **S36 Fig**. Same as **S6 Fig** but calculated for His26.

S37 Fig. Titration curves of titratable residues after 40ns for R#1-2, and 30ns for R#3.

**S38 Fig. Comparison between mean of calculated pK_a_ values of titratable residues and model pK_a_ values.** The model pK_a_ values are taken from the penta-peptide CH_3_CO-AA*X*AA-NH_2_, where *X* represents a titratable amino acid. (A-C) R#1-3.

**2.** H-bond networks

**S39 Fig. Intramolecular H-bond networks involving the entire residues (that is, backbone and side chain), averaged over all replica’s simulations at pH 7, using the H-bond angle criteria of 60^0^ or less (A) and of 20^0^ or less (B).** The H-bonds may be either direct or involve 1,2, or 3 water molecules. *Left:* H-bond occupancies (expressed in percentages) shown in a residue heat map . *Right*: H-bond network. The number of H-bonds (reported either if the occupancy is 50% or more (A) or 10% or more (B)) is shown in purple, whereas the number of water molecules is in orange.

**S40 Fig**. Same as **S39 Fig** but calculate for pH 6. **S41 Fig**. Same as **S39 Fig** but calculate for pH 5. **S42 Fig**. Same as **S39 Fig** but calculate for pH 4. **S43 Fig**. Same as **S39 Fig** but calculate for pH 3.

**S44 Fig.** Same as **S39 Fig - S43 Fig (A),** *right* but calculated only for R#1. **S45 Fig.** Same as **S39 Fig - S43 Fig(A),** *right* but calculated only for R#2. **S46 Fig.** Same as **S39 Fig - S43 Fig (A),** *right* but calculated only for R#3**. S47 Fig.** Same as **S39 Fig - S43 Fig (B),** *right* but calculated only for R#1. **S48 Fig.** Same as **S39 Fig - S43 Fig (B),** *right* but calculated only for R#2.

**S49 Fig.** Same as **S39 Fig - S43 Fig (B),** *right* but calculated only for R#3.

**S50 Fig.** Same as **S39 Fig,** but only for the side chains, and using an occupancy of 25% for bond angle criterion of 60^0^ or less instead of 50% (pH 7). Water mediated and direct H-bonds included.

**S521 Fig.** Same as **S50 Fig** but calculate for pH 6. **S52 Fig.** Same as **S50 Fig** but calculate for pH 5. **S53 Fig.** Same as **S50 Fig** but calculate for pH 4. **S54 Fig.** Same as **S50 Fig** but calculate for pH 3.

**S55 Fig.** Same as **S50 Fig - S54 Fig** (A), *right* but calculated only for R#1, R#2 and R#3.

**S56 Fig.** Same as **S50 Fig - S54 Fig** (B), *right* but calculated only for R#1, R#2 and R#3.

**S57 Fig.** Same as **S50 Fig**. except that only direct H-bonds are considered and an occupancy of 15% is set for the criterion of H-bond angle of 60^0^. The pH is 7.

**S58 Fig.** Same as **S57 Fig** but calculated for pH 6. **S59 Fig.** Same as **S57 Fig** but calculated for pH 5. **S60 Fig.** Same as **S57 Fig** but calculated for pH 4. **S61 Fig.** Same as **S57 Fig** but calculated for pH 3.

**S62 Fig**. Same as **S57 Fig - S61 Fig** (A), *right* but calculated only for R#1, R#2 and R#3.

**S63 Fig**. Same as **S57 Fig - S61 Fig** (B), *right* but calculated only for R#1, R#2 and R#3, except for pH 3, where the minimum occupancy is 6%.

### 3. NPY conformation dynamics

Standard MD simulations. Here all the titratable side chains were in their standard protonation states, as chosen at the beginning of the simulation. During the final 110 ns of each of the three MD simulations (lasting 500 ns), the α-helical segment (Glu15–Ile31), was conserved whereas the N- and C-terminal regions were rather disordered. A summary of the three analyses is offered in **S5 Table** and **S64–S69 Fig**s.

**S64 Fig**. Secondary structure content of pNPY, plotted as a function of time, in the three standard MD simulations, for the last 110 ns (A: R#1, B: R#2, C: R#3). The a-helix is colored in magenta, turn in teal the extended configuration in yellow, 3-10 helix in blue and coil in white.

Figure prepared with VMD[32].

**S65 Fig**. Same as **S64 Fig** but for constant pH simulations at pH 7, for all the timescale explored in replicas R#1-3.

**S66 Fig**. Same as **S64 Fig** but at pH 6. **S67 Fig**. Same as **S64 Fig** but at pH 5. **S68 Fig**. Same as **S64 Fig** but at pH 4. **S69 Fig**. Same as **S64 Fig** but at pH 3.

**S70 Fig. Top:** Fluctuations as backbone as quantified by PAD[60] values, calculated for CpHMD simulations at pH 7 and pH 6. Bottom: Bottom: mean PAD values for the 3 replicas and standard deviation.

**S71 Fig.** Same as **S70 Fig**, calculated for CpHMD simulations at pH 6 and pH 5.

**S72 Fig.** Same as **S70 Fig**, calculated for CpHMD simulations at pH 5 and pH 4.

**S73 Fig.** Same as **S70 Fig**, calculated for CpHMD simulations at pH 4 and pH 3.

S1 Table. Mean pKa values obtained from our simulations.

**S2 Table.** Occupancies of either direct H-bonds or water-mediated bridges among entire residues (that is, backbone and side chains), averaged over the constant pH simulations at different pH values. This and the next two tables have as occupancies as those in S39-S49 Figs (50% and 10% for criteria of 60^0^ and 20^0^, respectively), S50-S56 (25 % and 10%, **Table S3**), and S57-S63 (15% and 10%, **Table S4**). The atoms involved in direct H-bonding are shown. They have the same color code as the amino acids they belong to. As expected, they belong to residues close by.

**S3 Table.** Same as **S2** but only for side chains and the occupancy of 25% for the criterion of 60^0^ and 10% for the criterion of 20^0^.

**S4 Table.** Same as **S3** but only for direct H-bonds and the occupancy of 15% for the criterion of 60^0^ and 10% for the criterion of 20^0^.

**S5 Table**. Helical secondary structure content in the last 110 of standard MD simulations (R#1-3) and for the last 40ns (R#1 and R#2) and 30ns (R#3) of the constant pH MD simulations.

S1 Text. H-Bond networks.

**ABBREVIATIONS**
NPY: Neuropeptide Y;
h(p)NPY: human (porcine) NPY;
MD simulation: Molecular dynamics simulation;
CpHMD simulations: Constant pH molecular dynamics simulations,
T-pad: The protein angular dispersion.

