## Supplementary figures and images for "pH-dependent structural dynamics of the neuropeptide Y in aqueous solution"

### S1.Fig

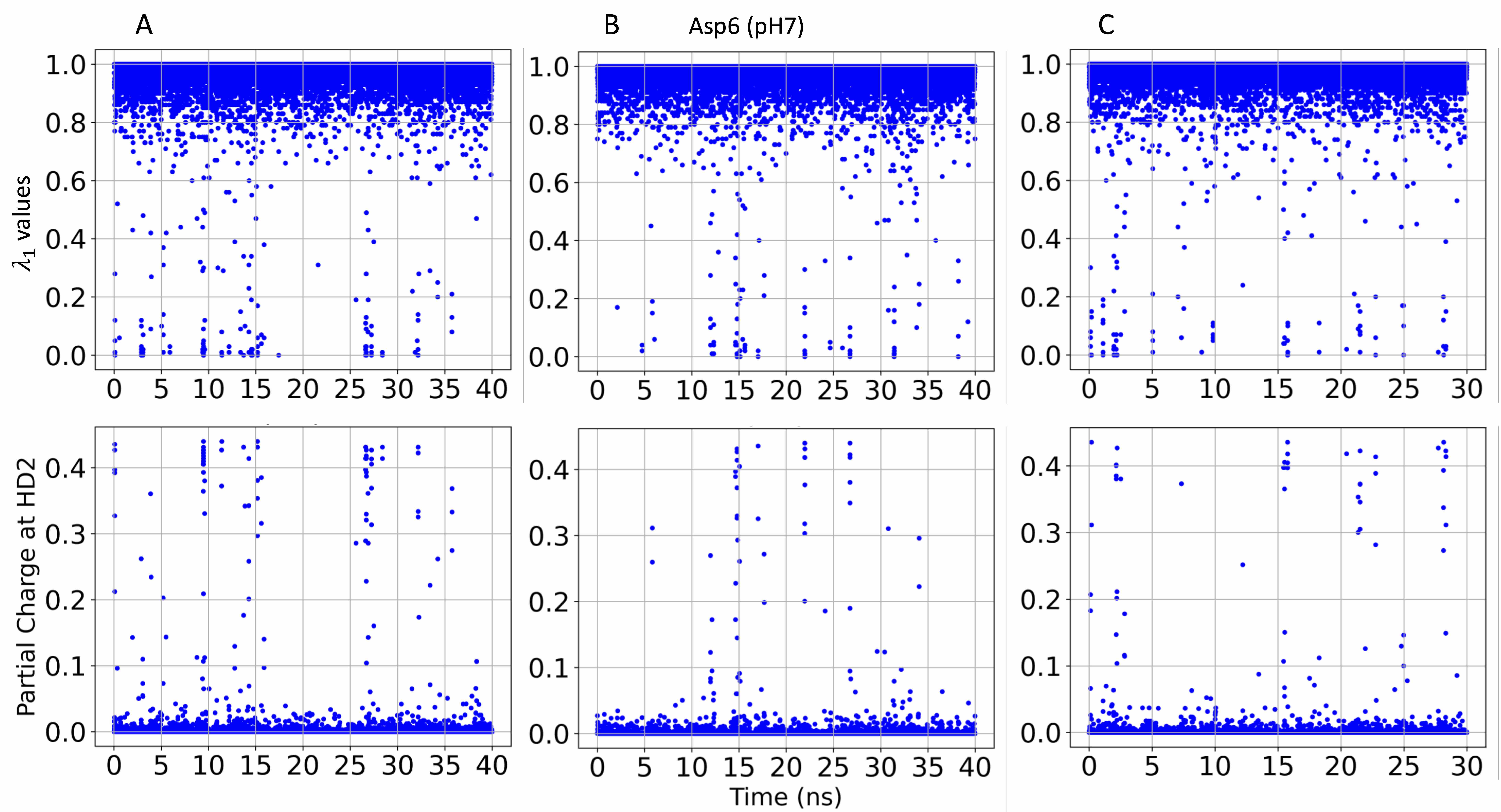

### S2.Fig

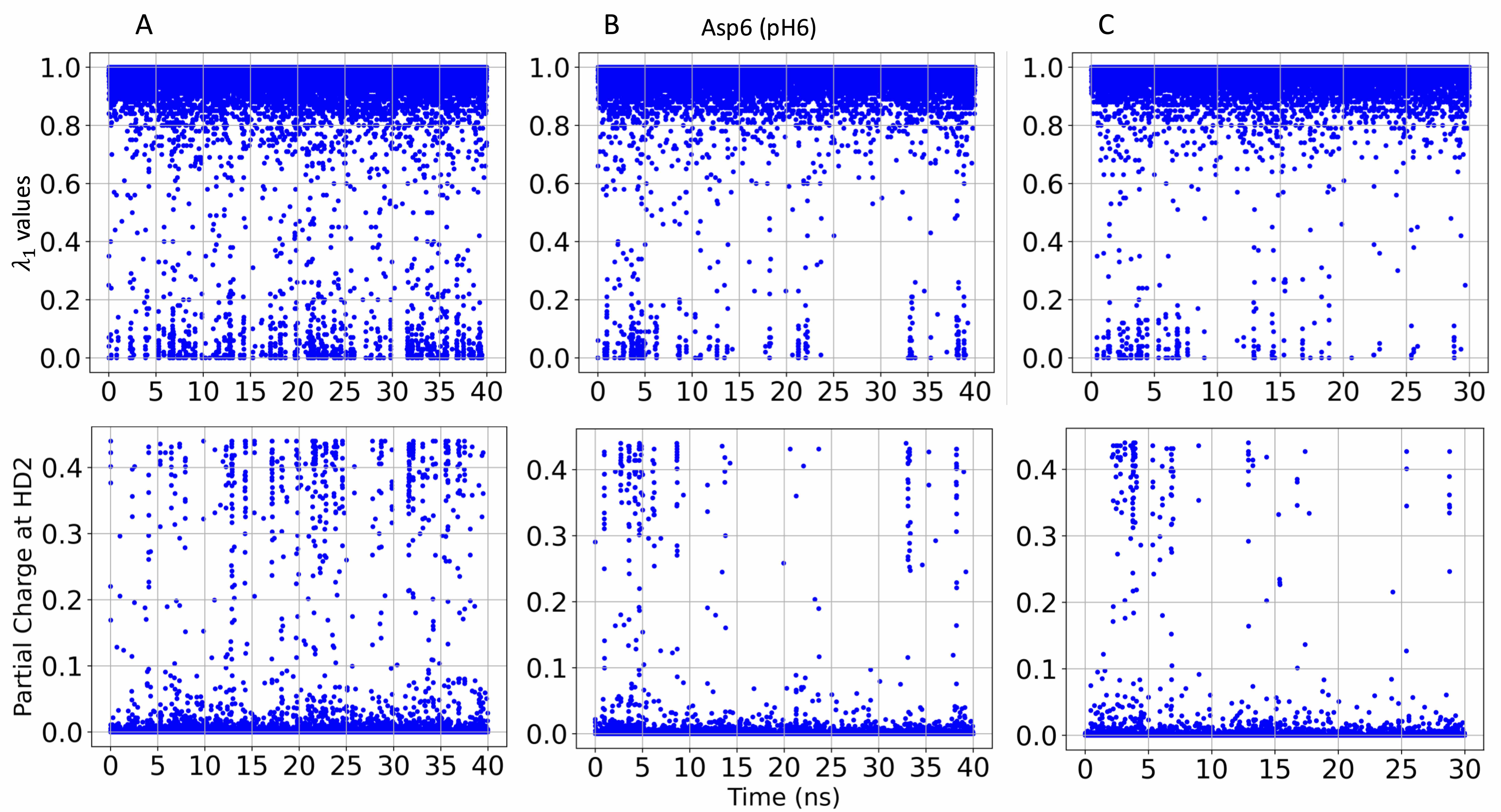

### S3.Fig

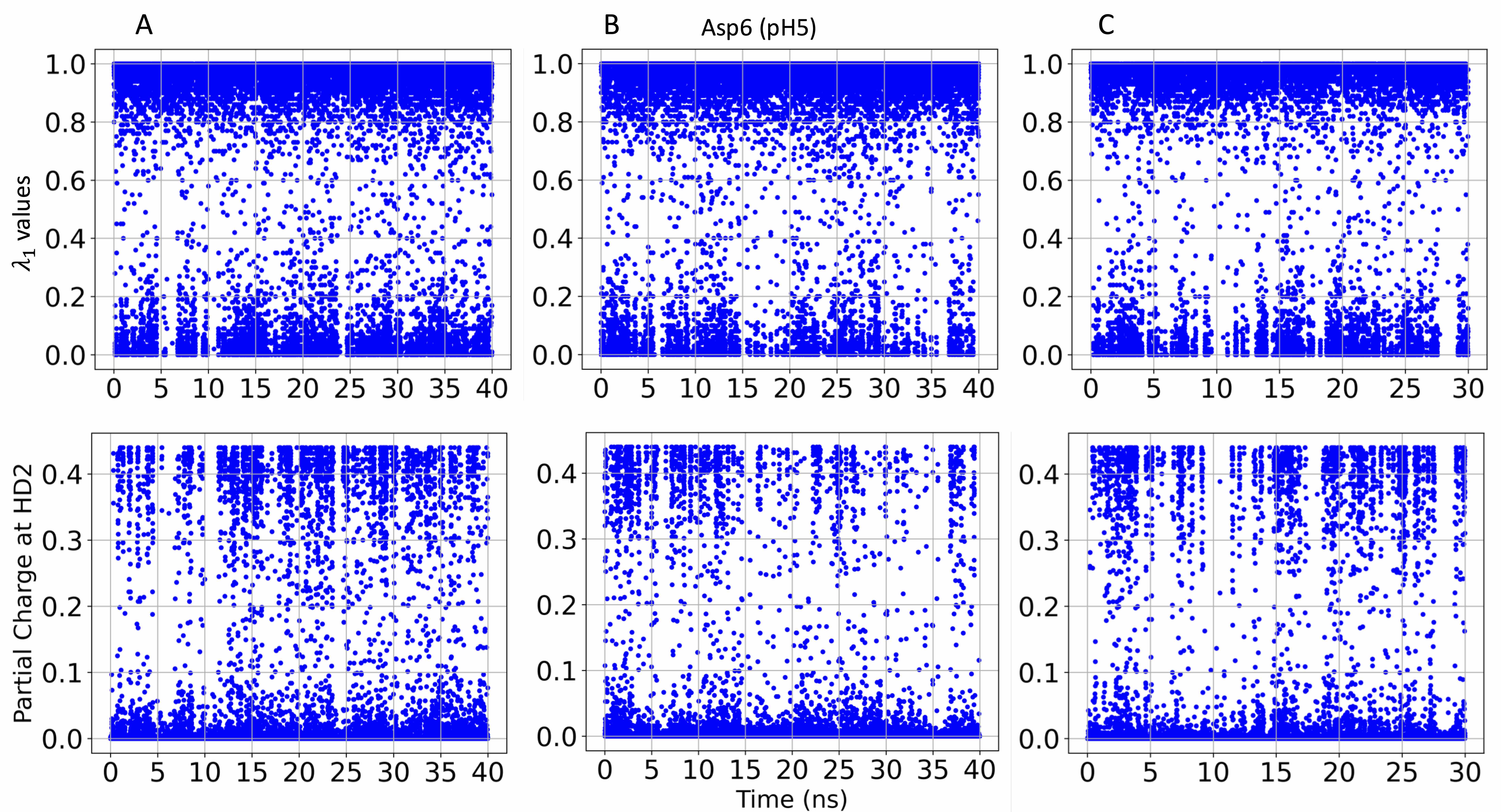

### S4.Fig

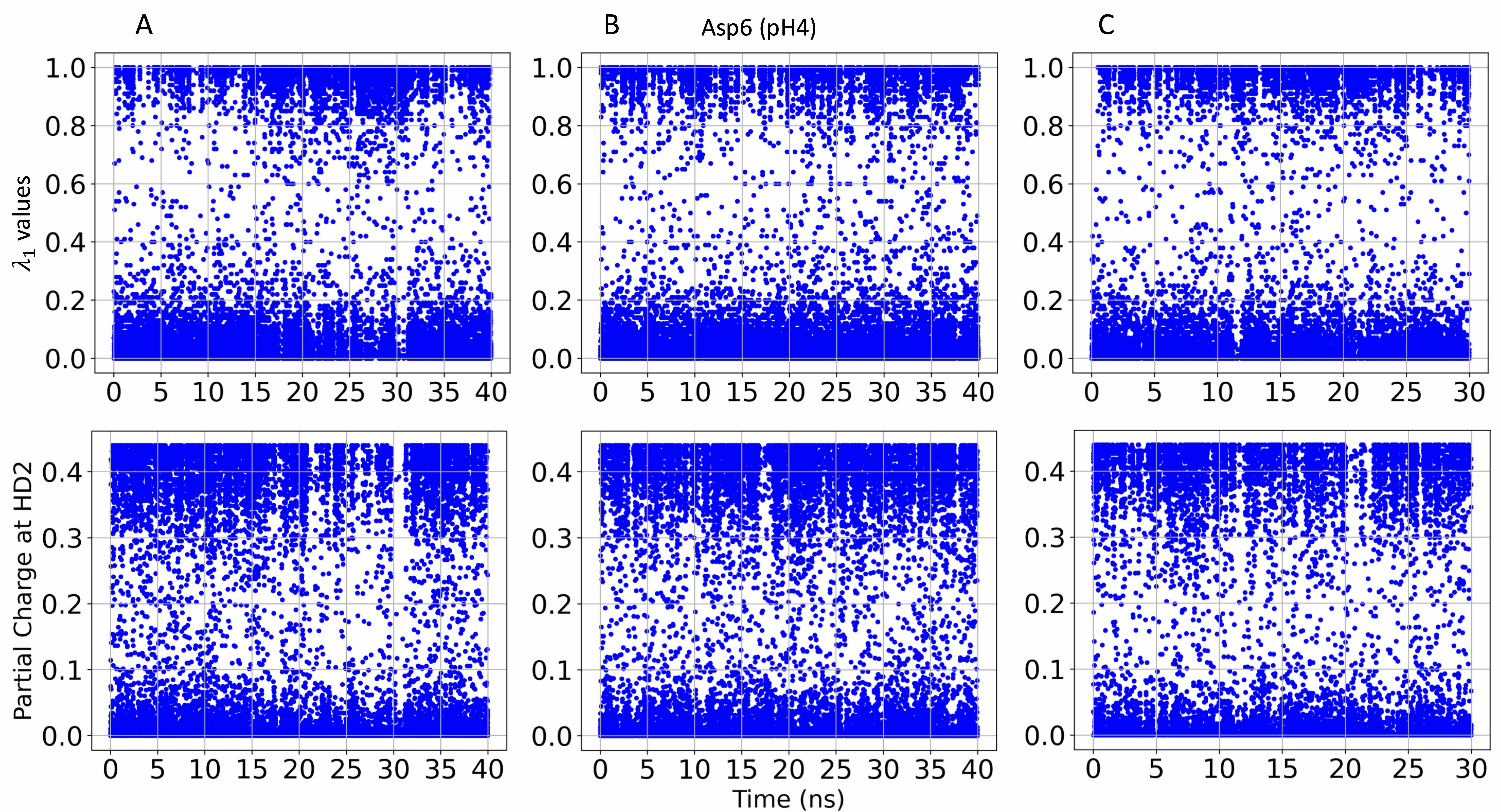

### S5.Fig

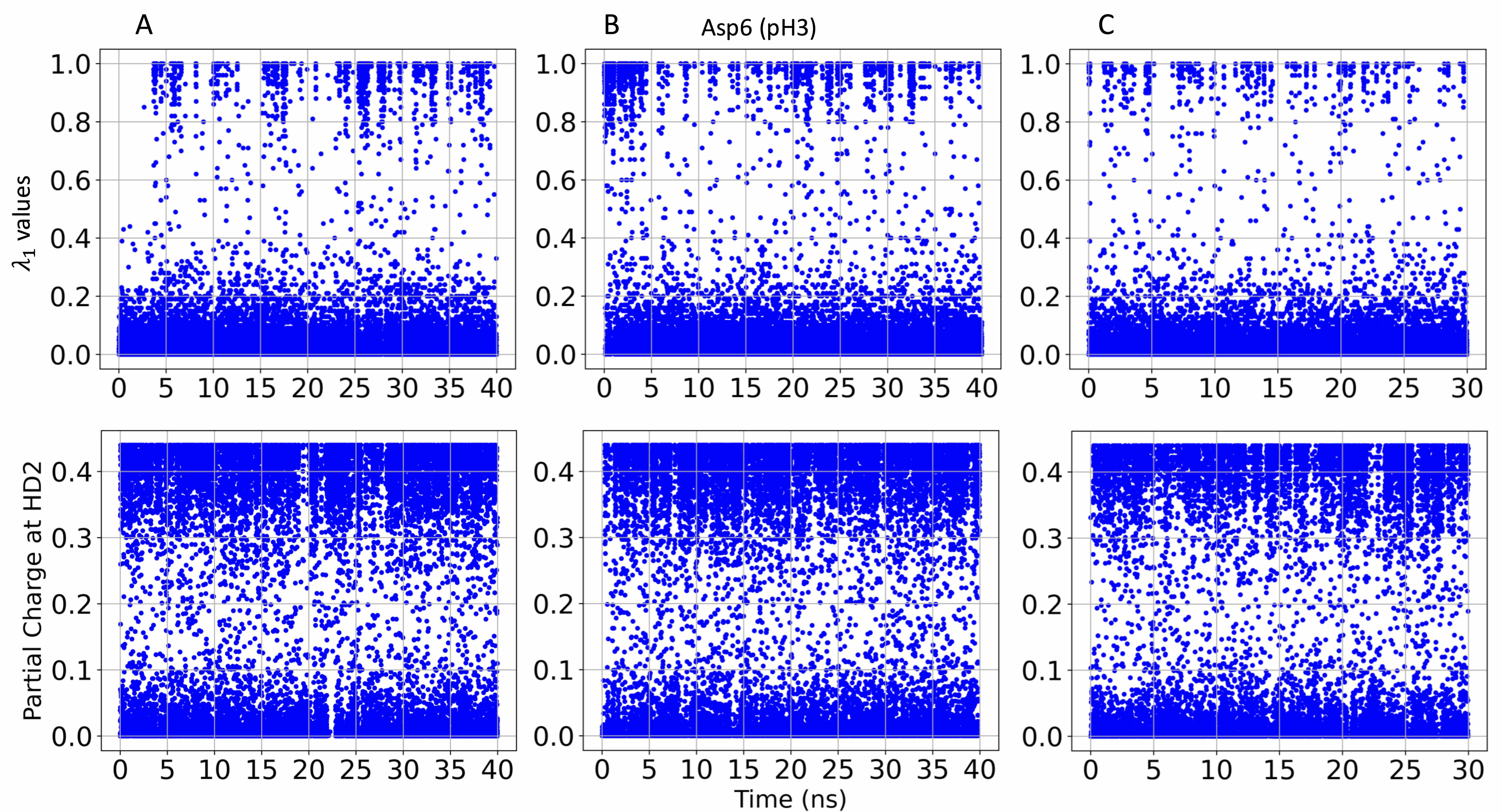

### S6.Fig

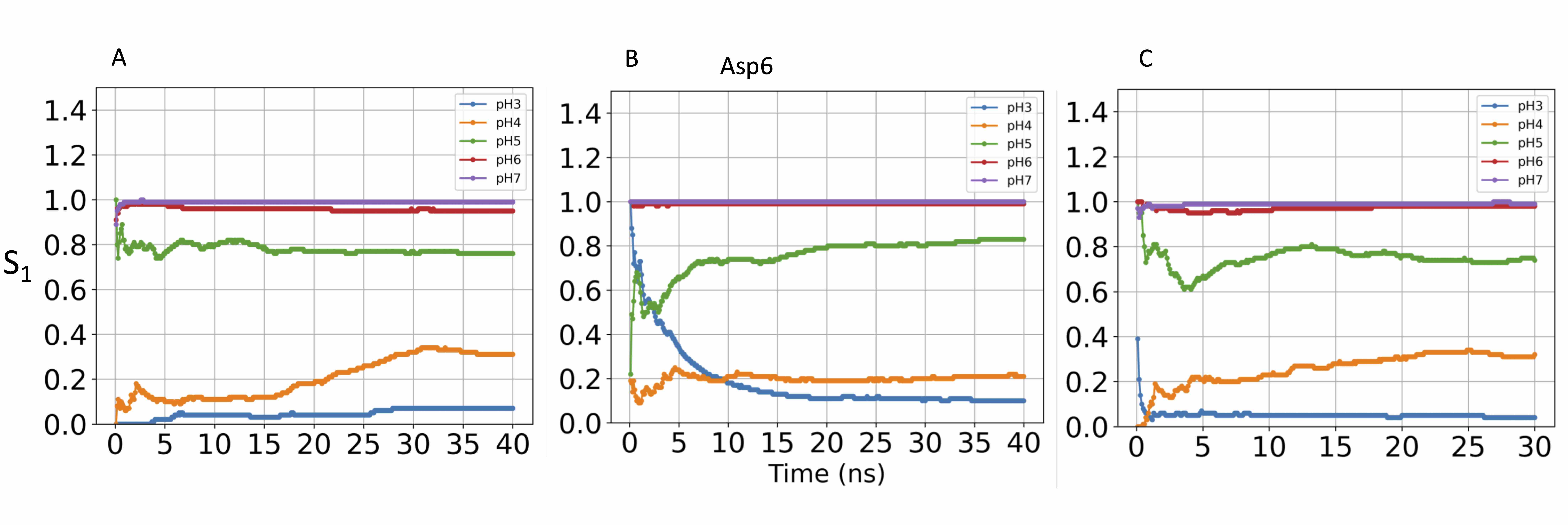

### S7.Fig

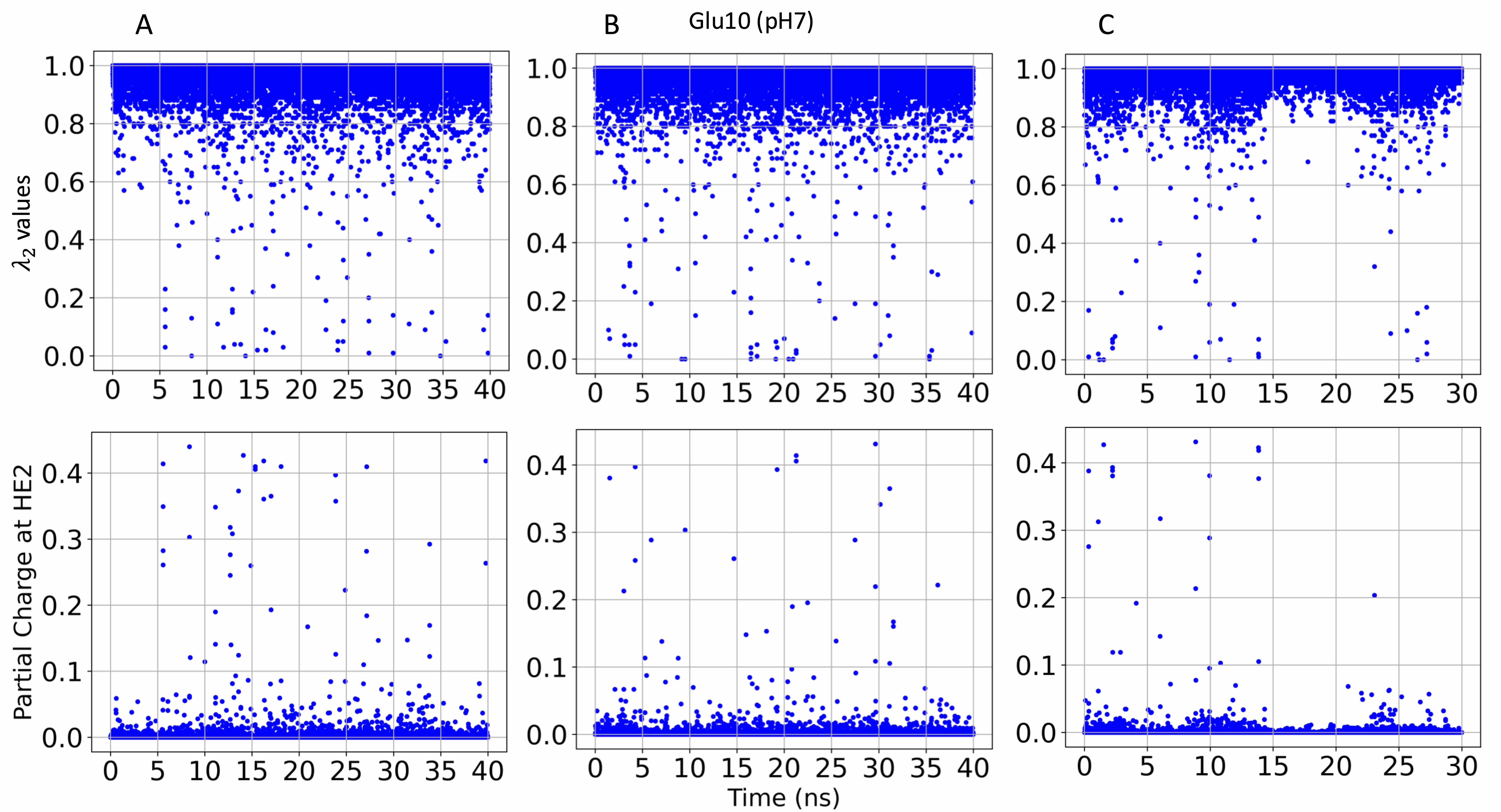

### S8.Fig

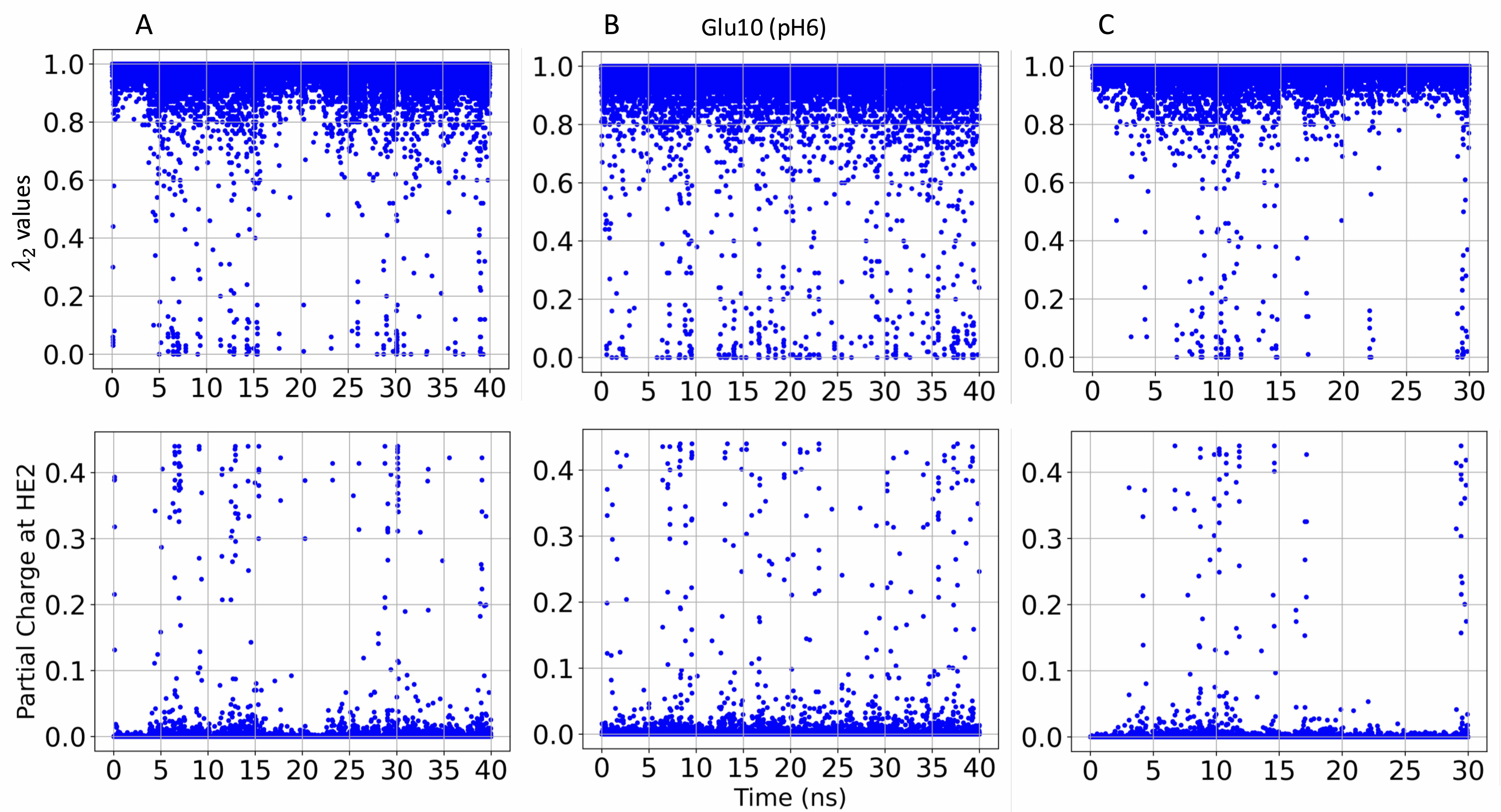

### S9.Fig

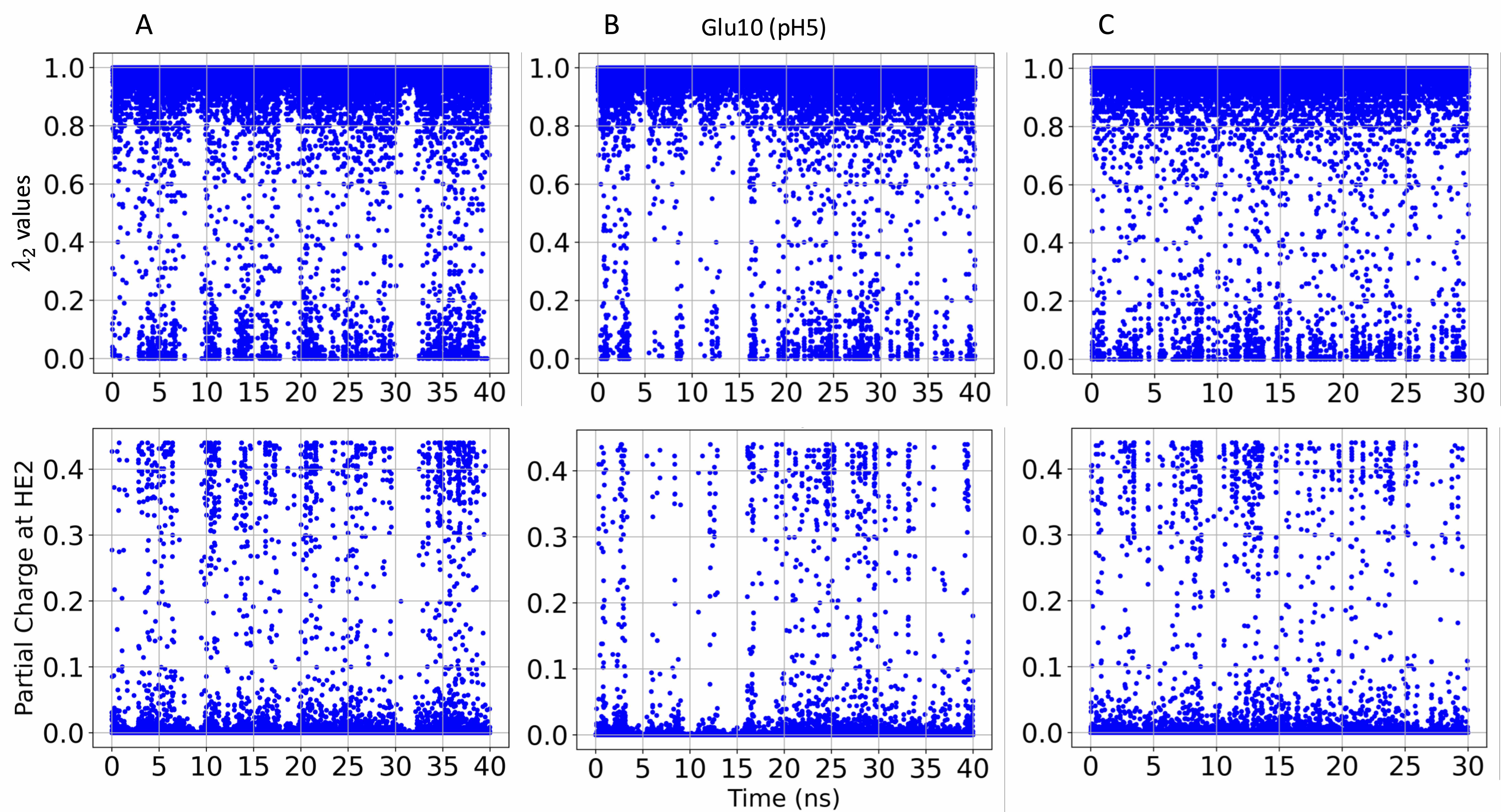

### S10.Fig

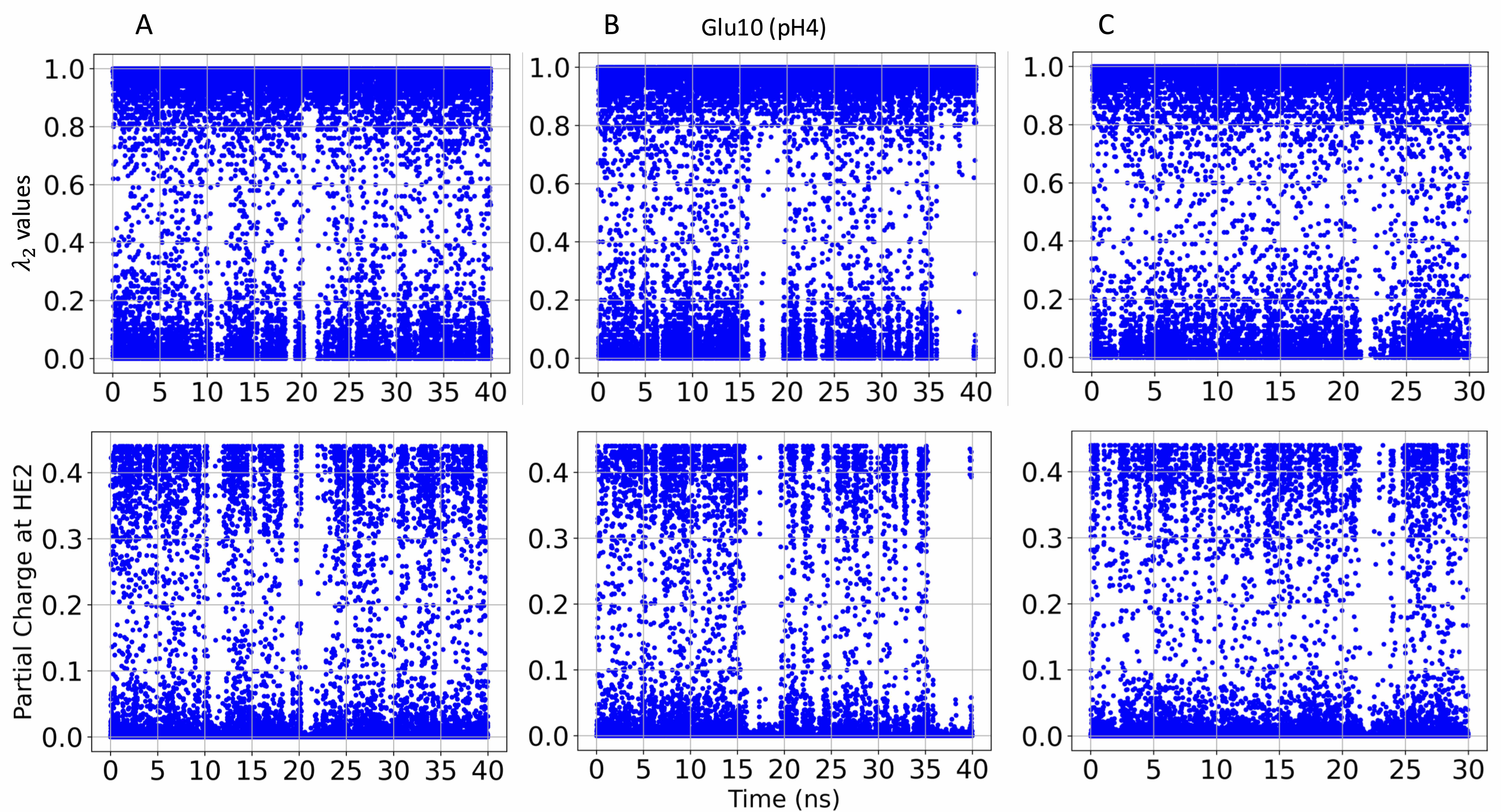

### S11.Fig

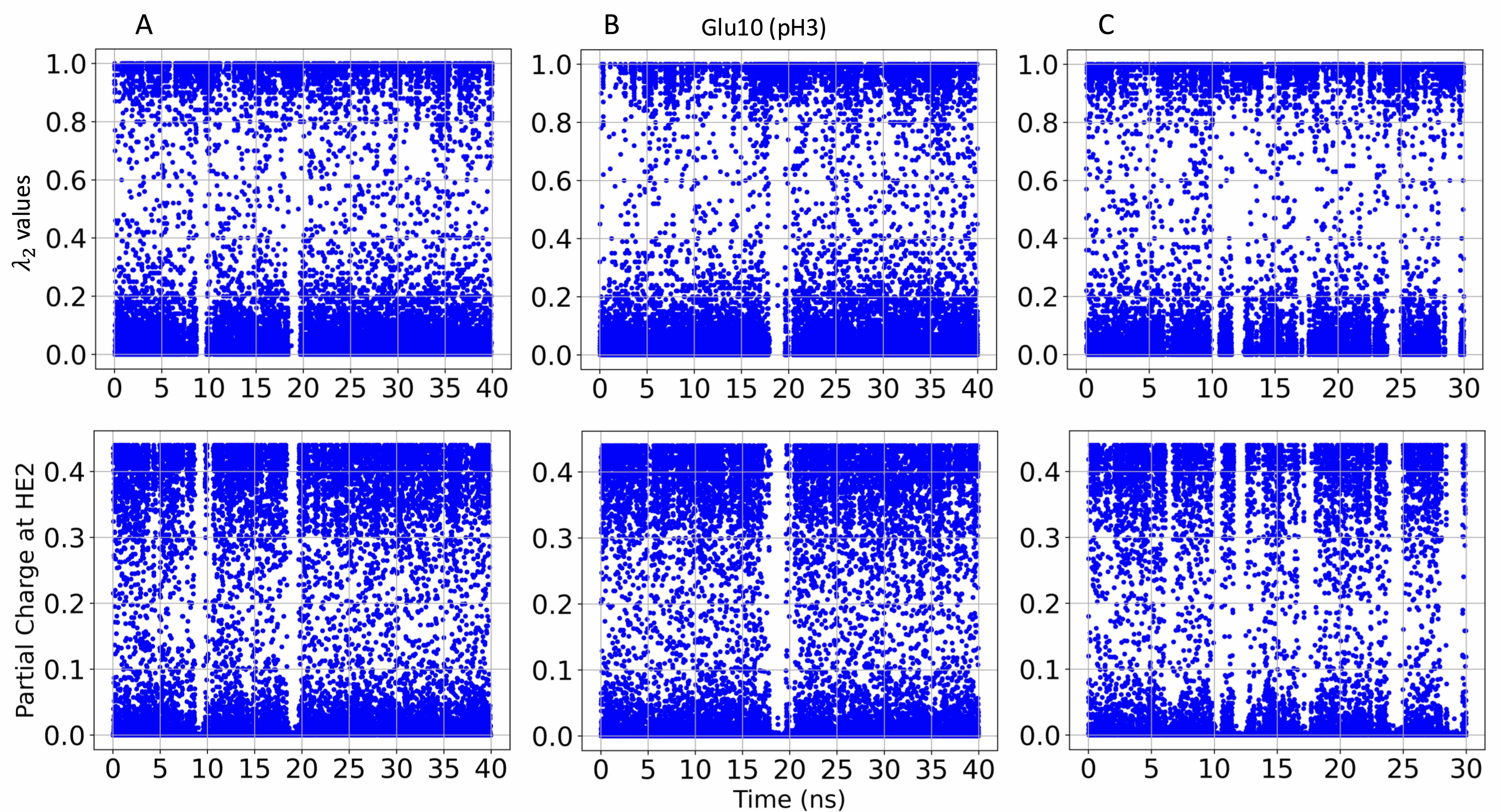

### S12.Fig

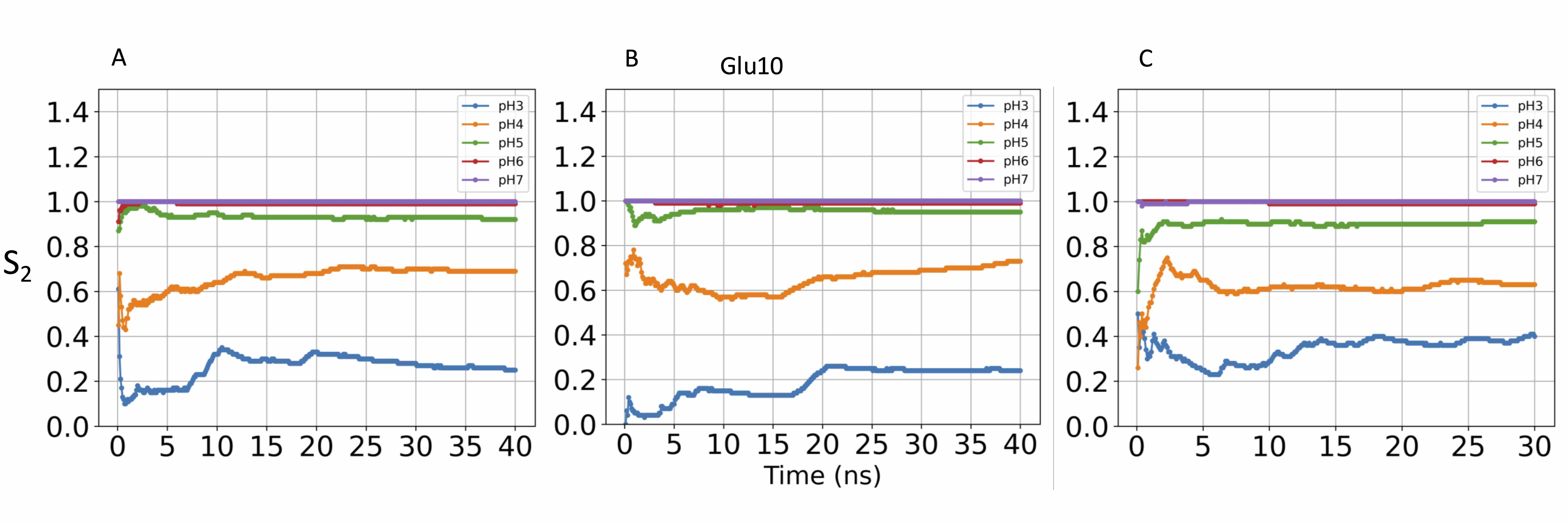

### S13.Fig

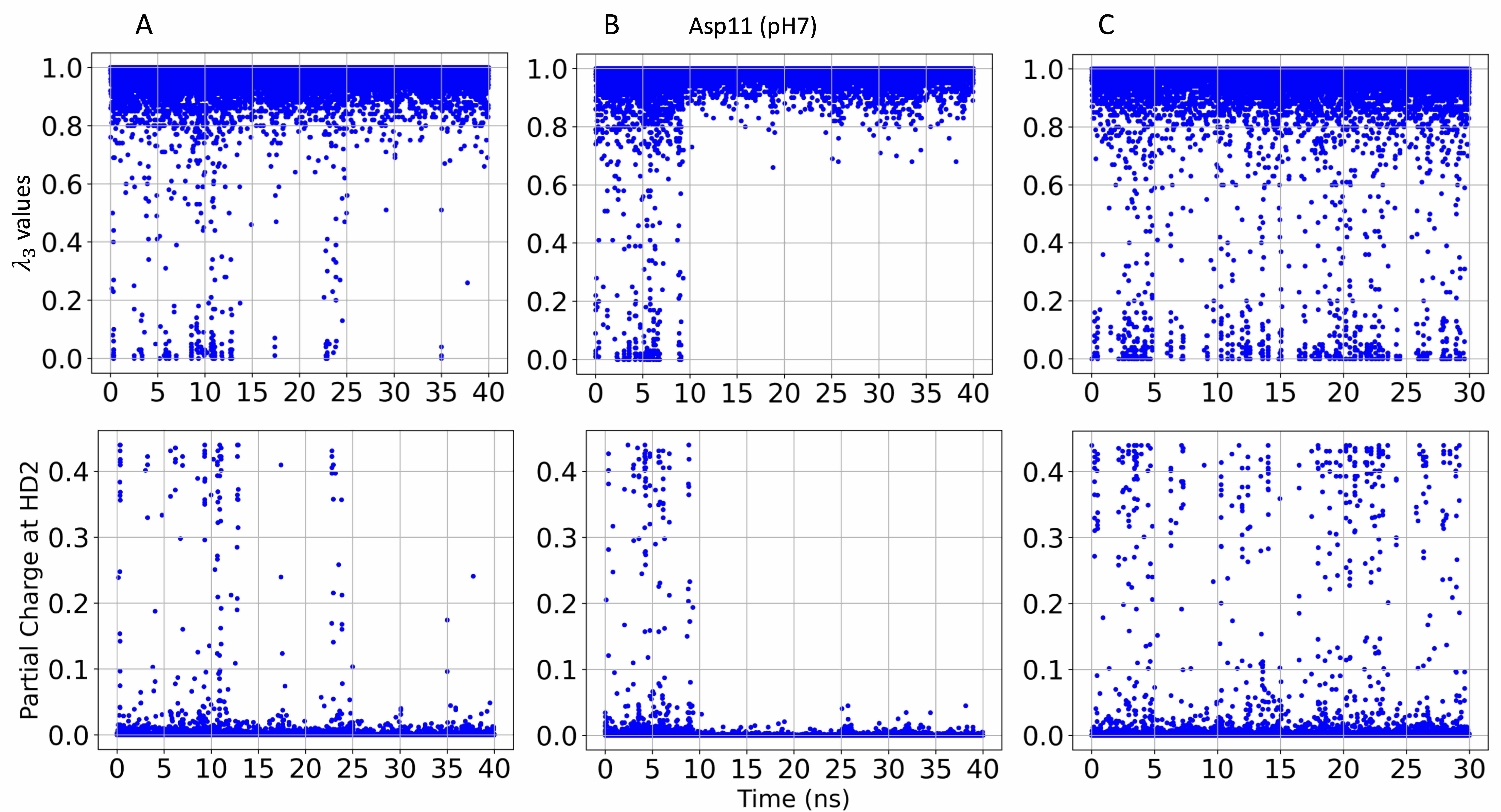

### S14.Fig

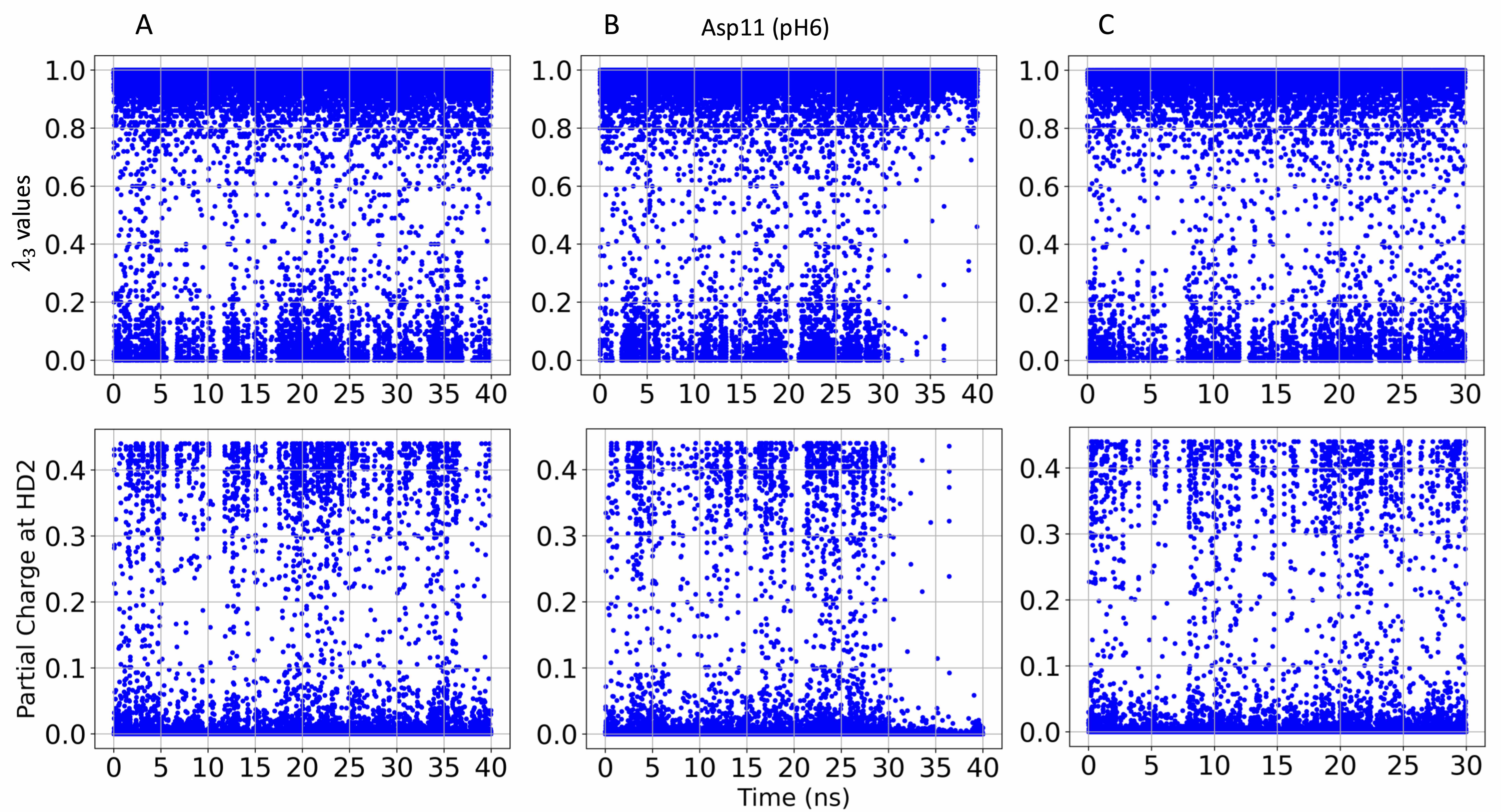

### S15.Fig

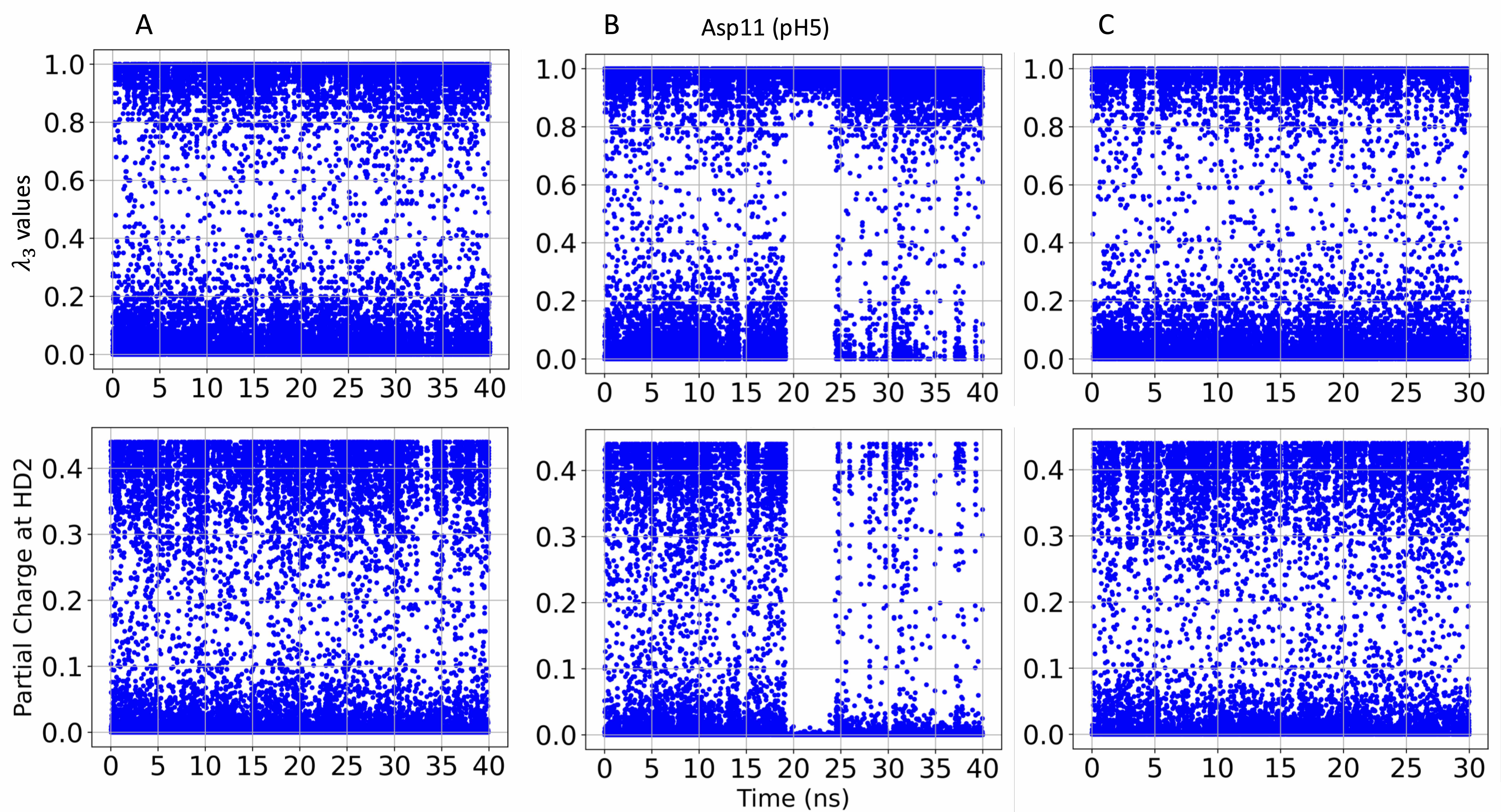

### S16.Fig

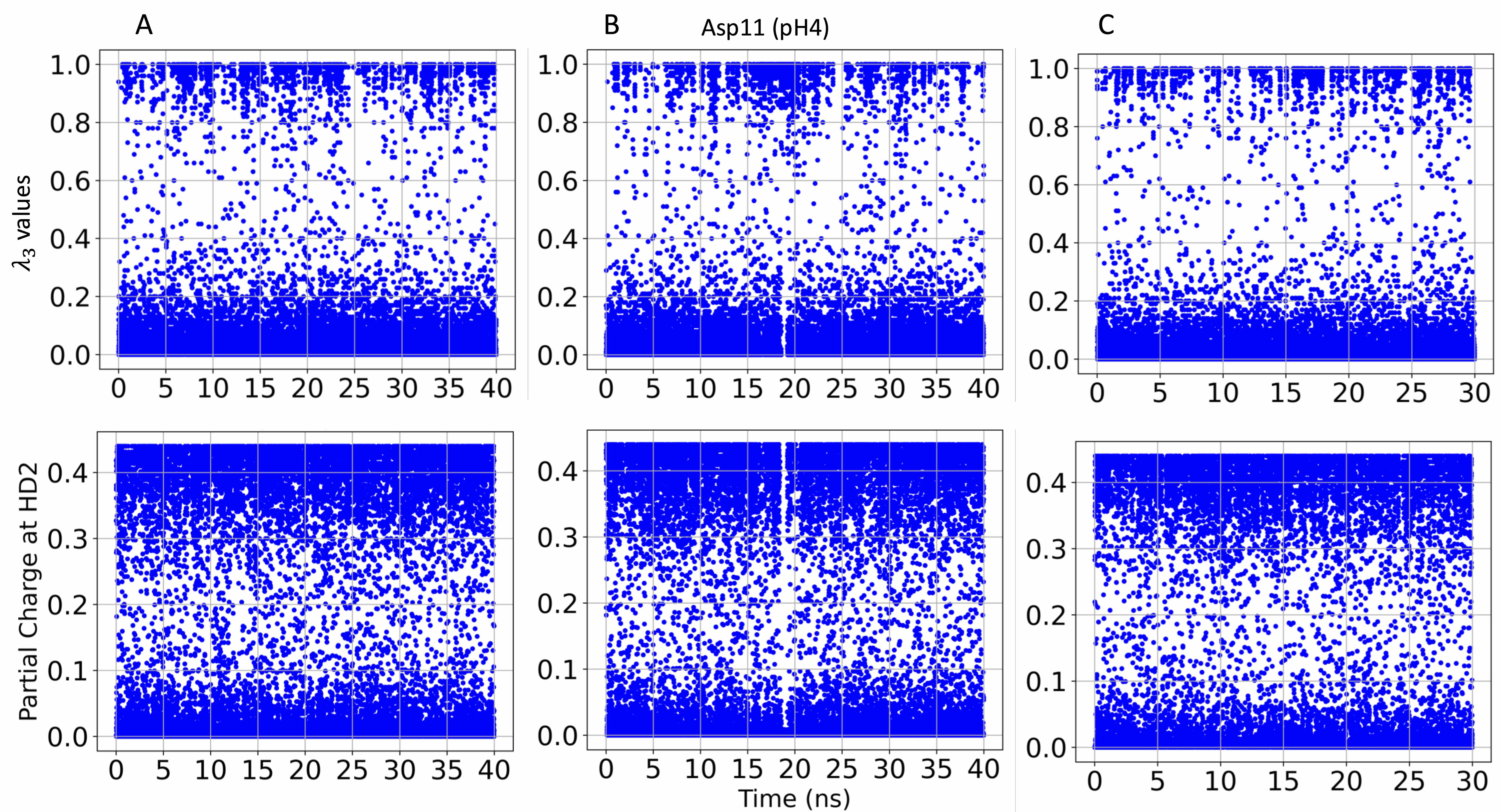

### S17.Fig

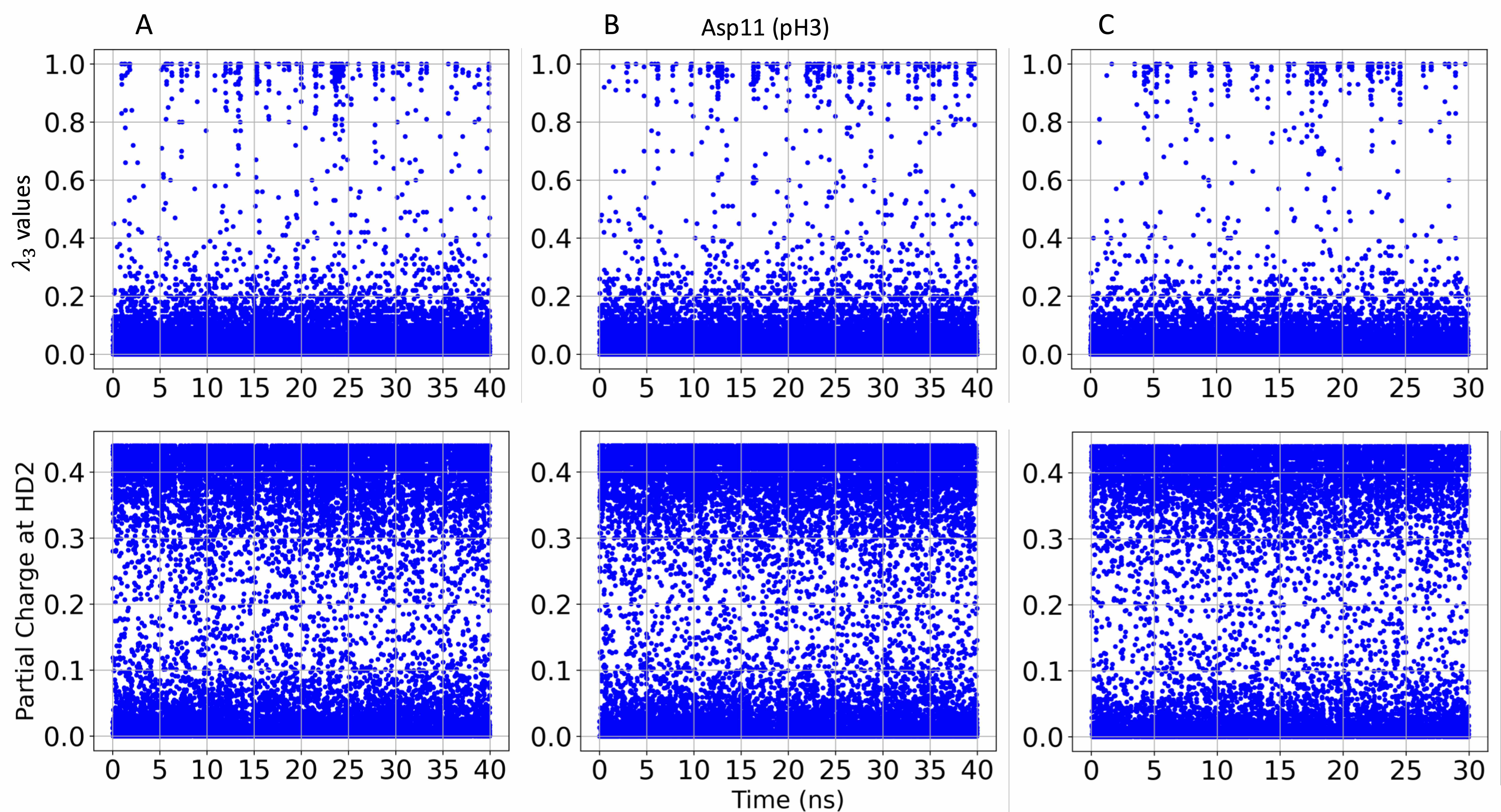

### S18.Fig

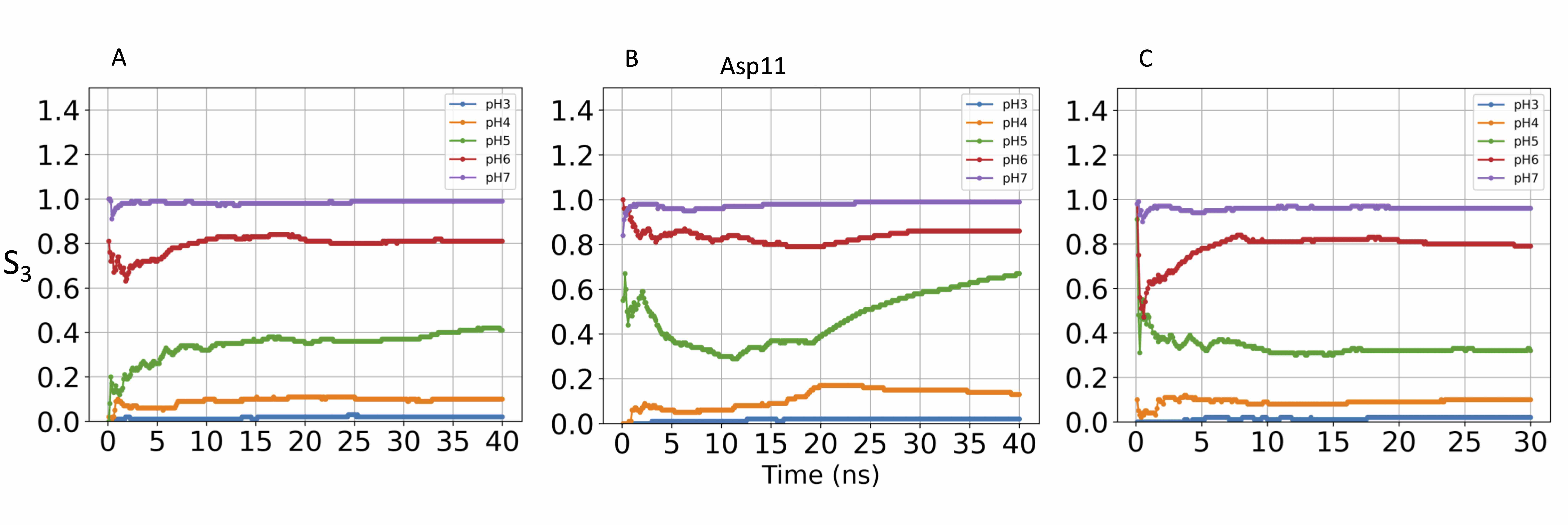

### S19.Fig

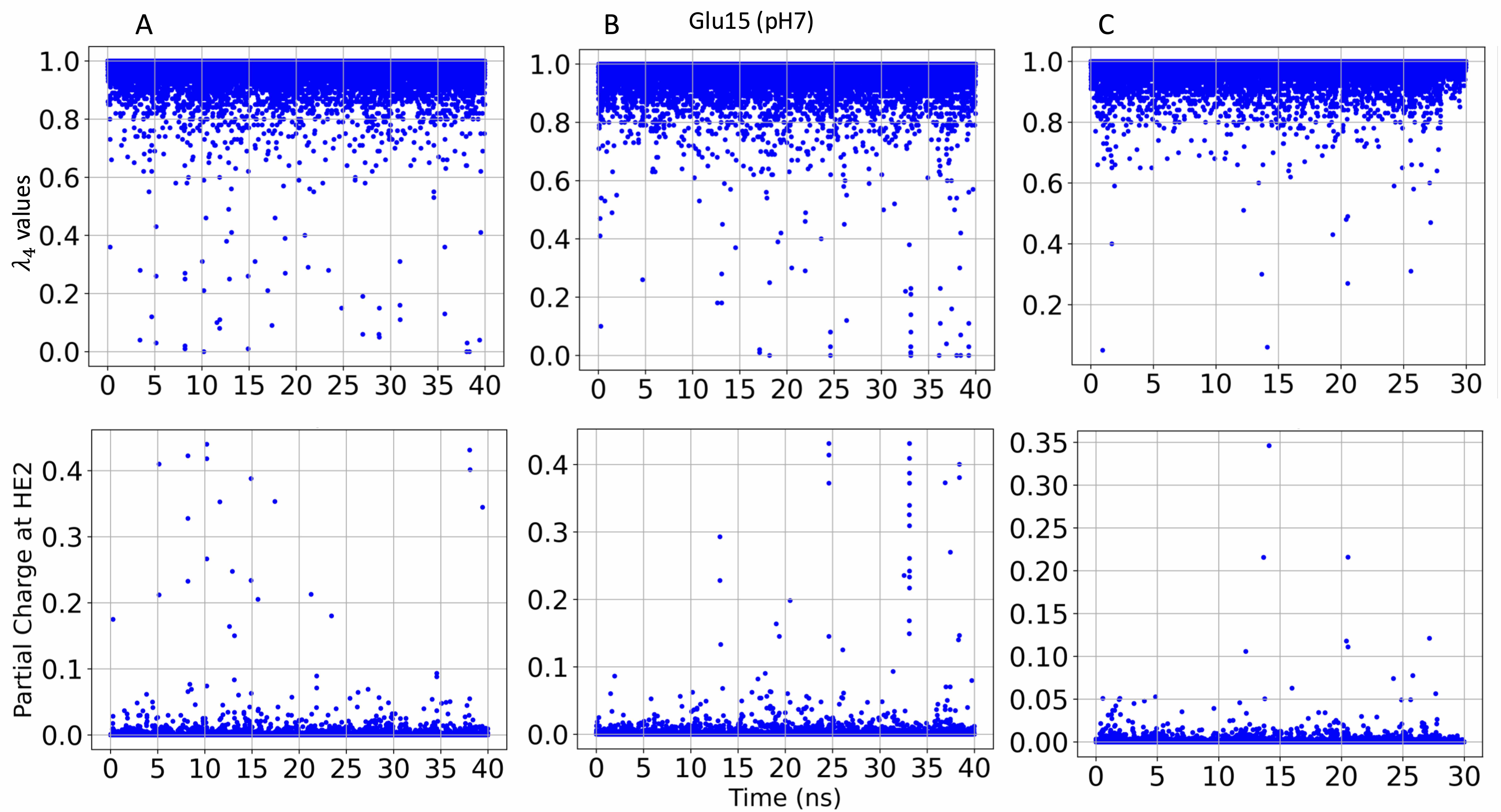

### S20.Fig

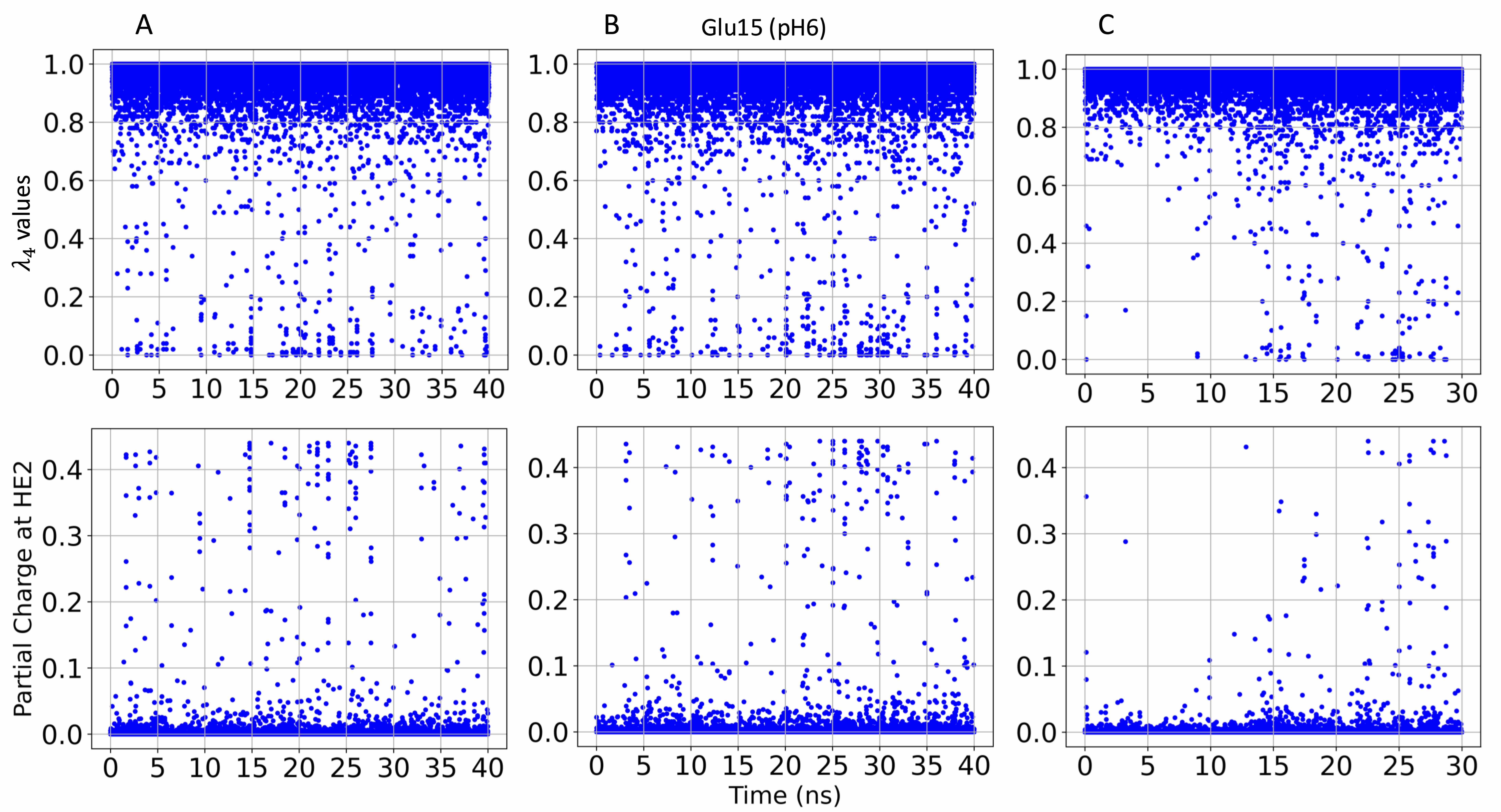

### S21.Fig

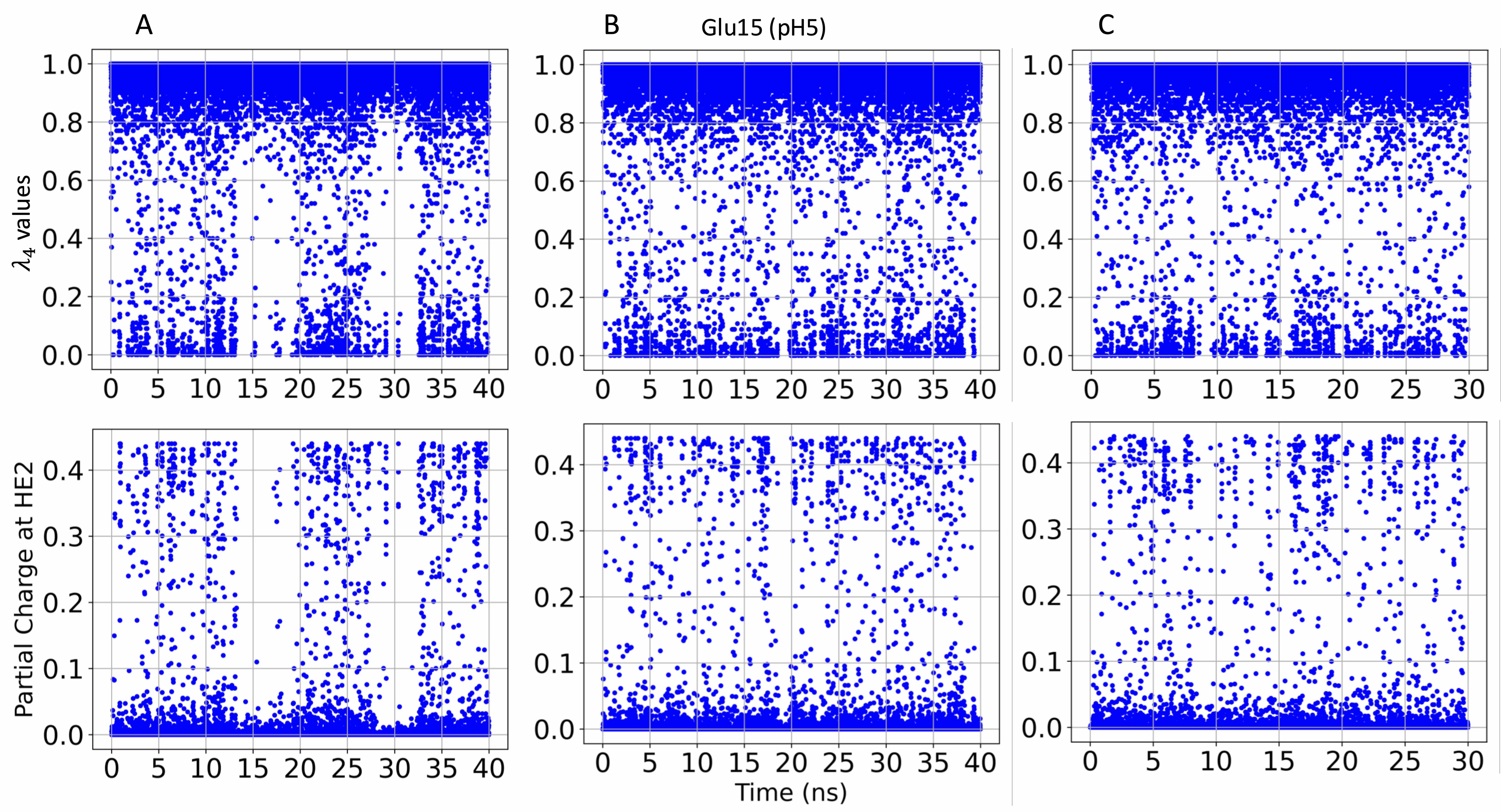

### S22.Fig

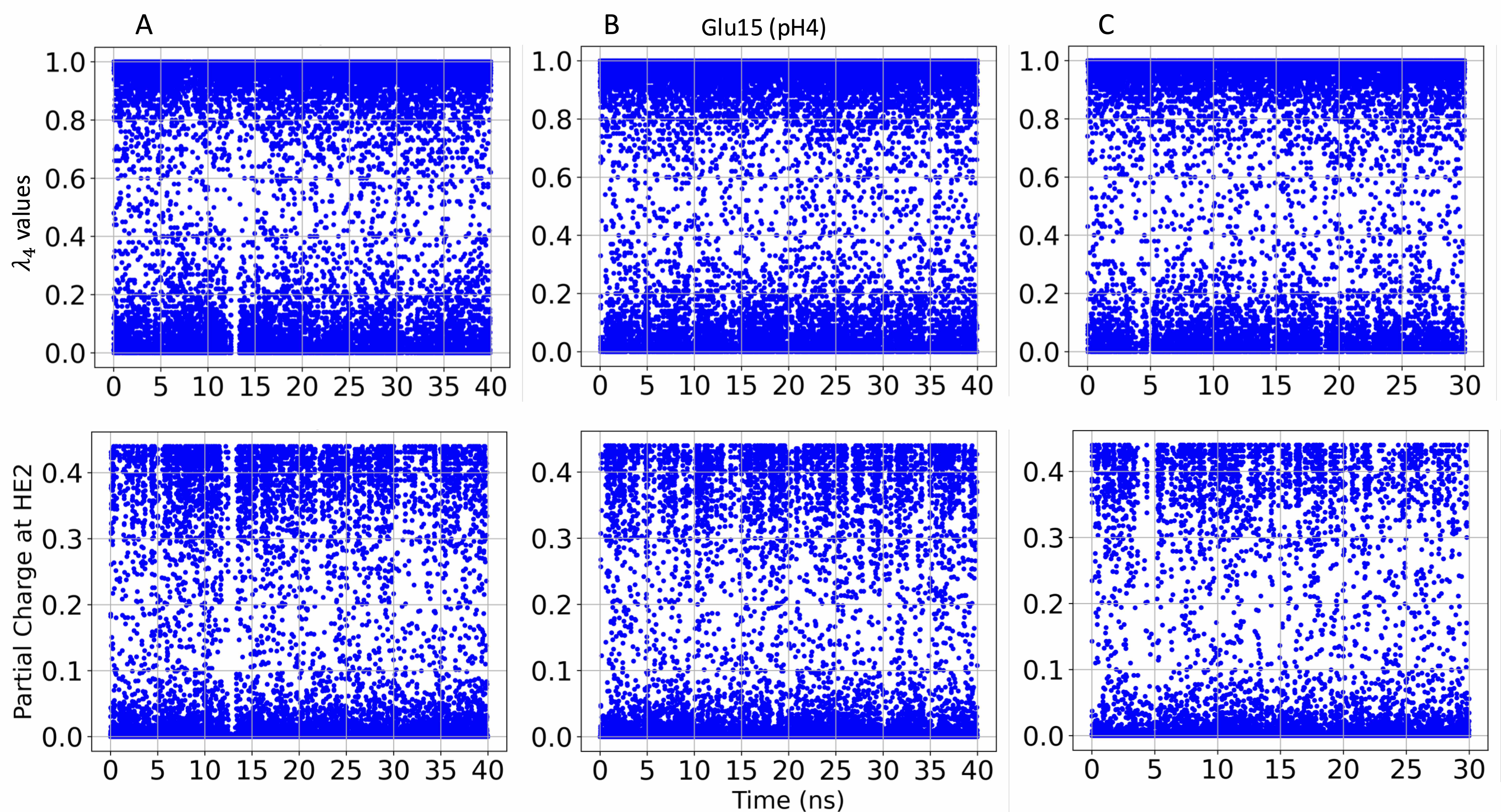

### S23.Fig

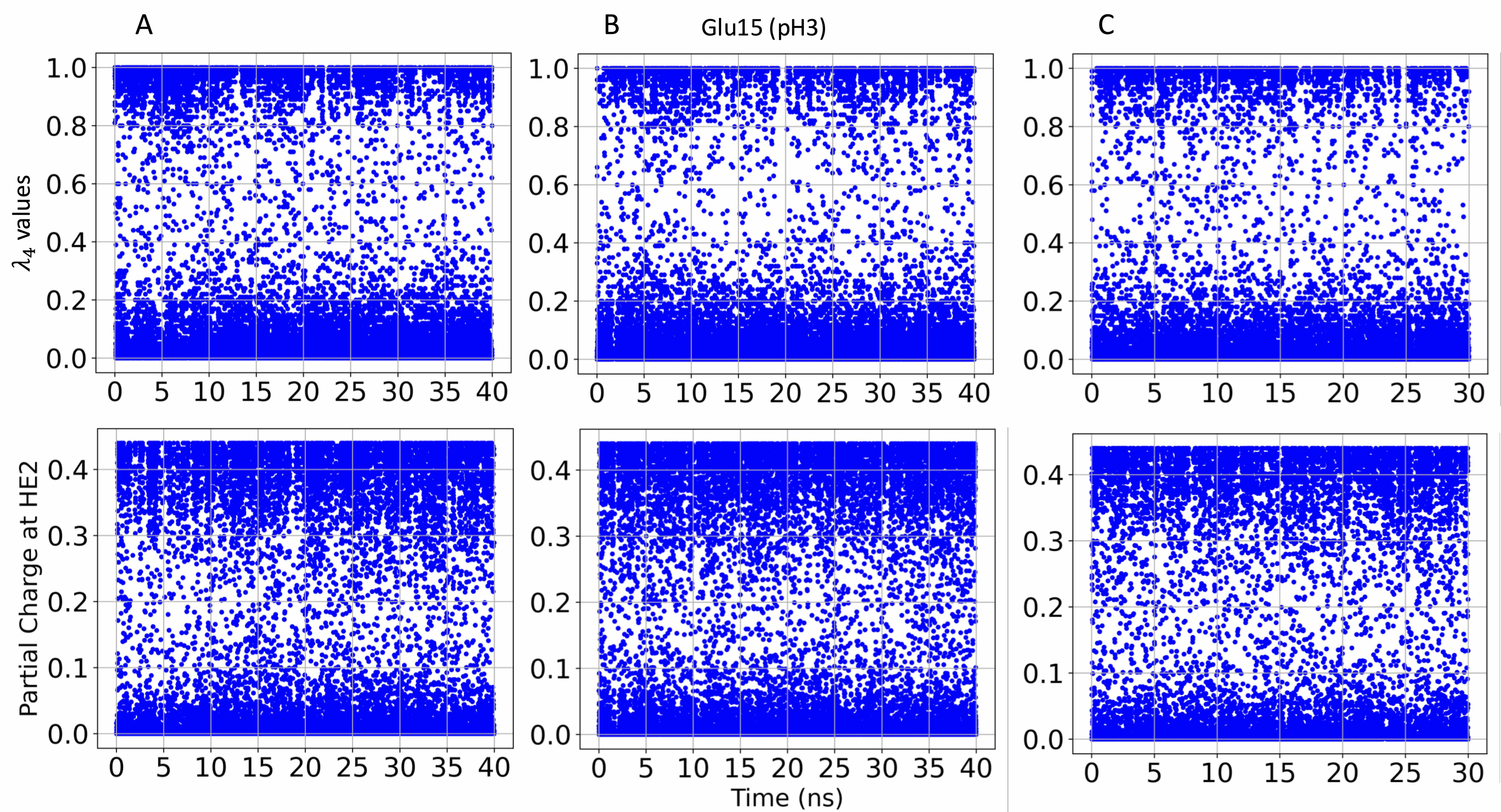

### S24.Fig

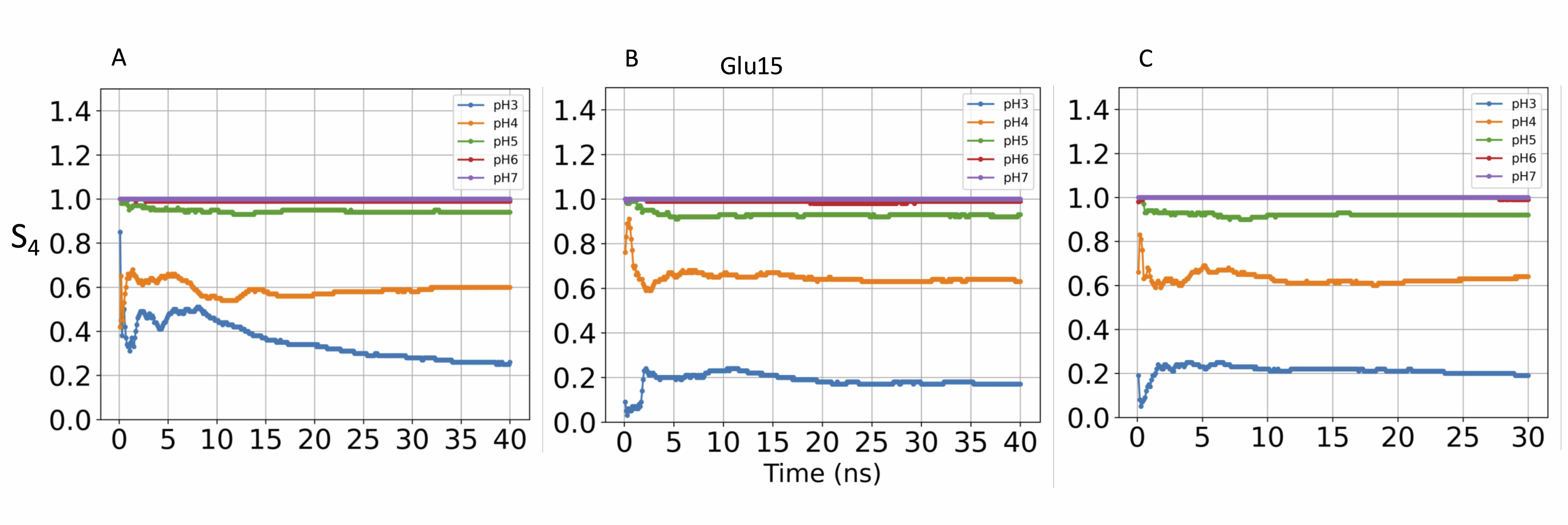

### S25.Fig

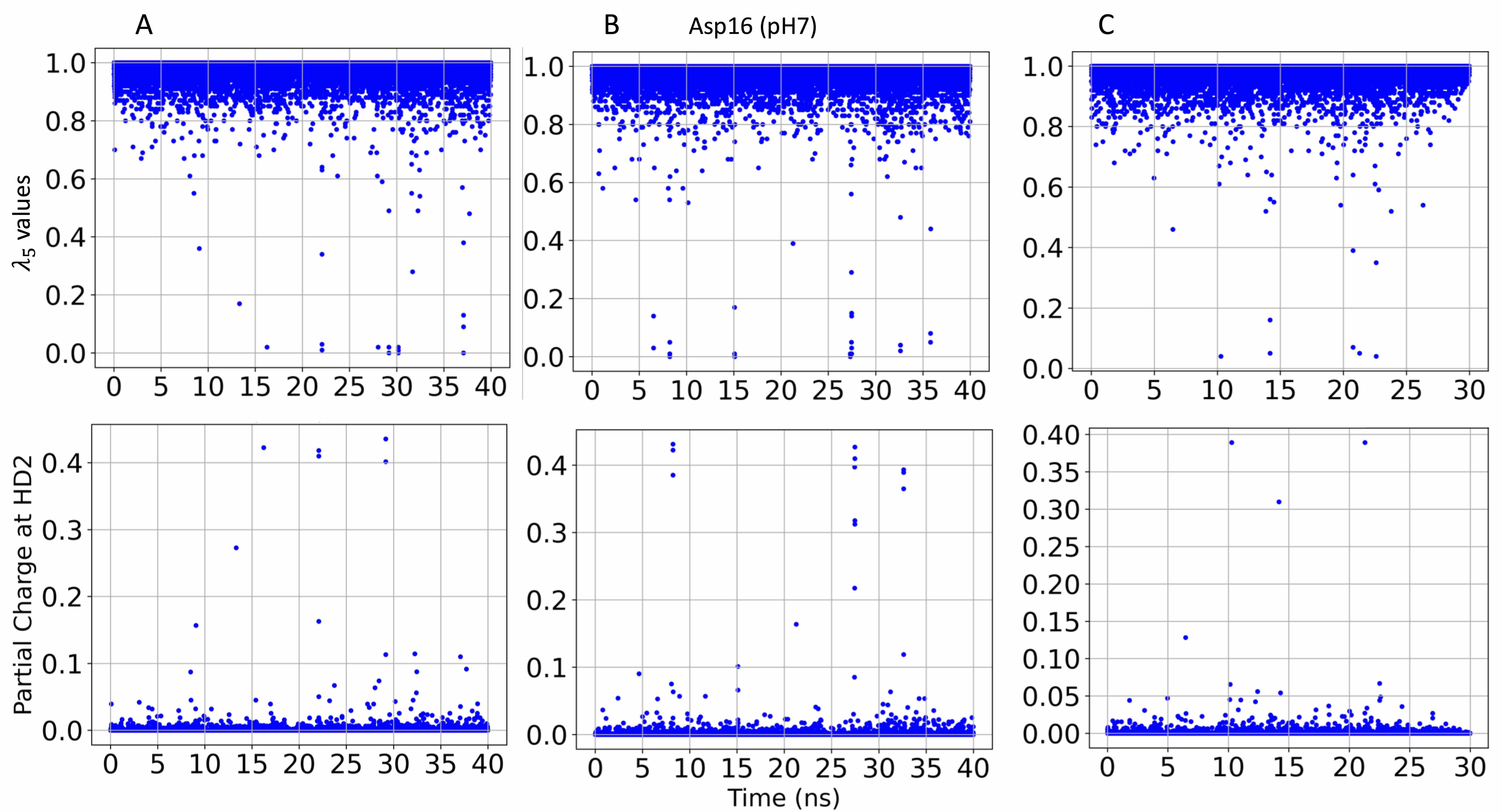

### S25.Fig

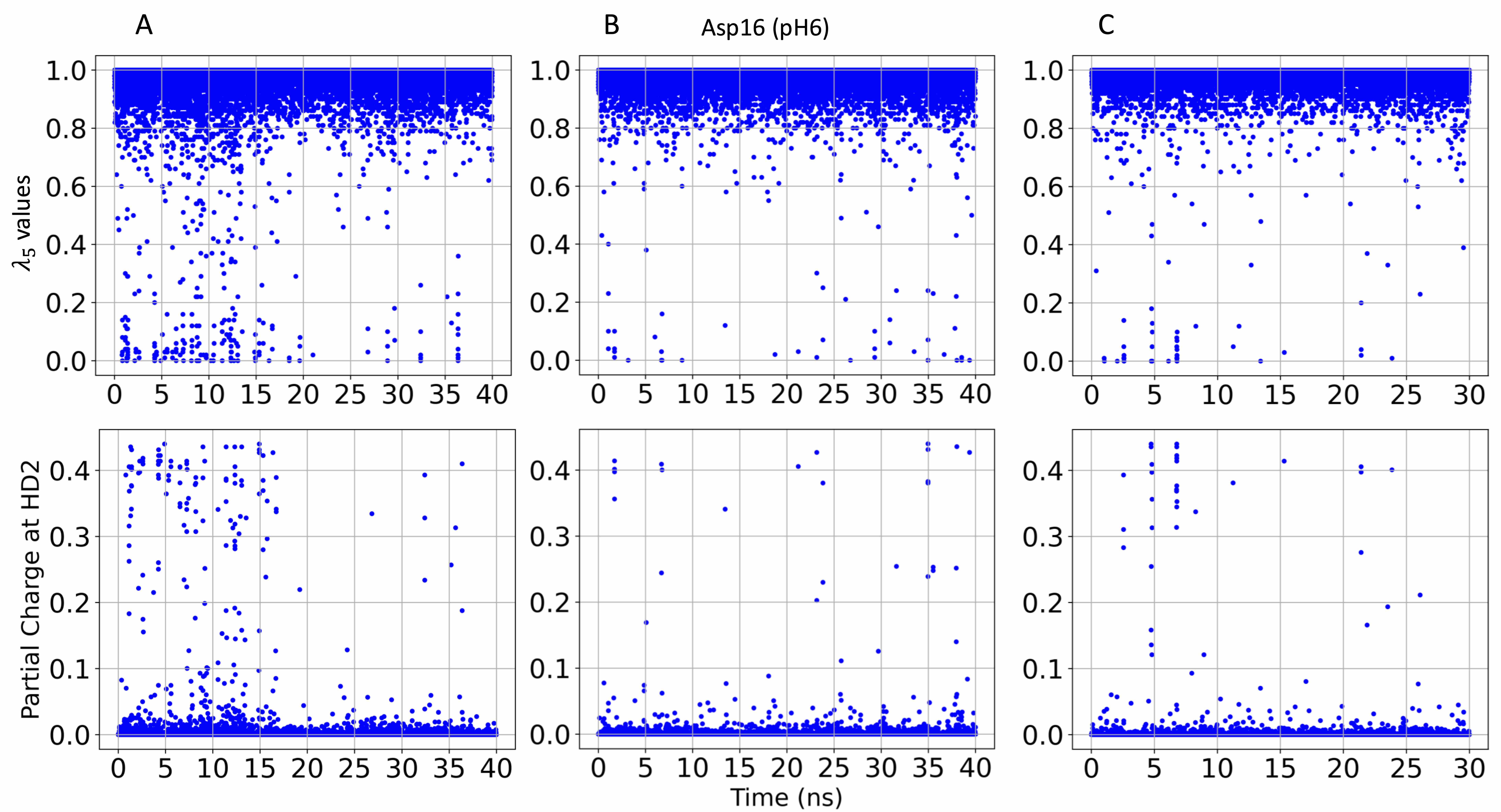

### S27.Fig

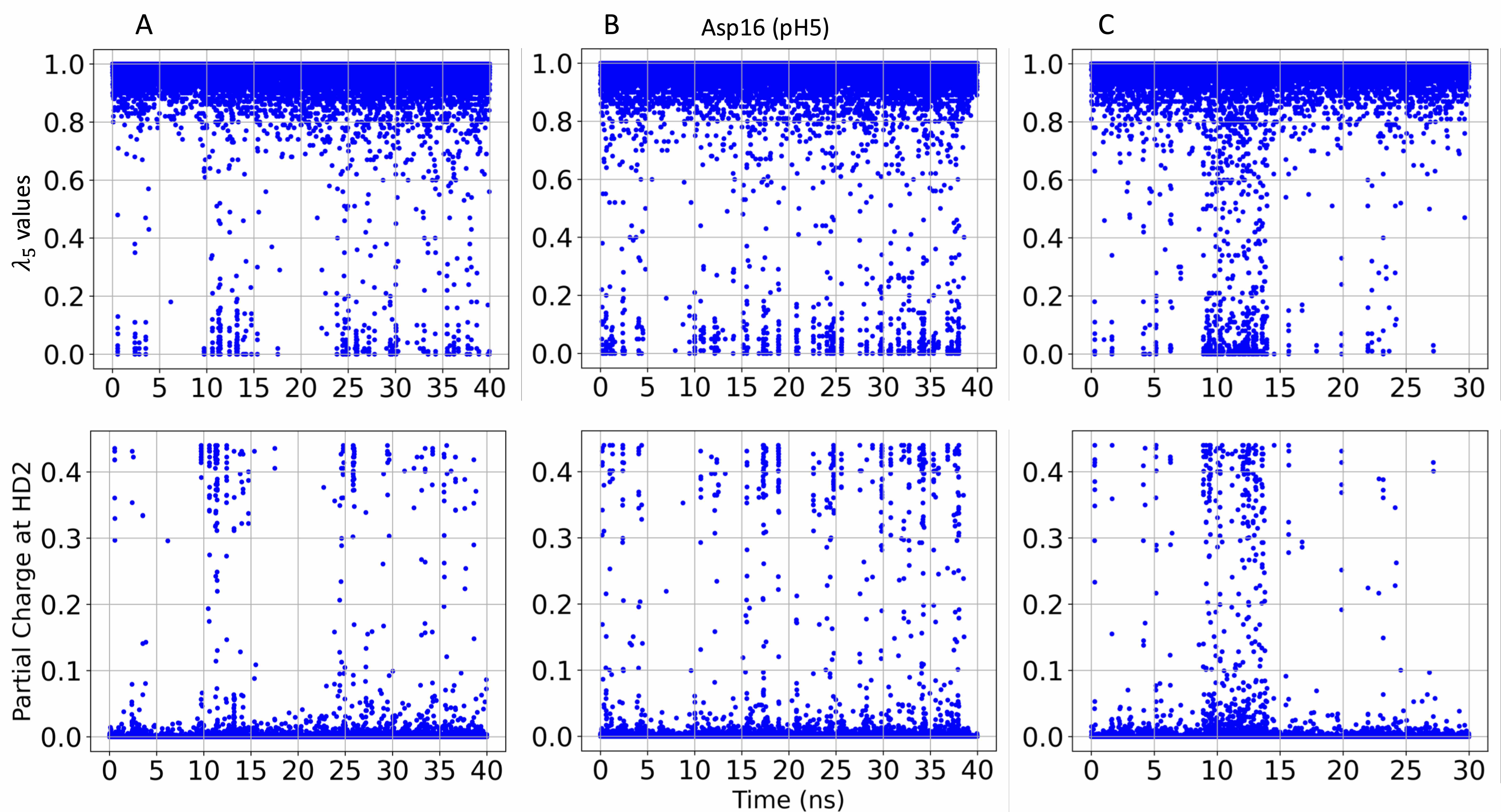

### S28.Fig

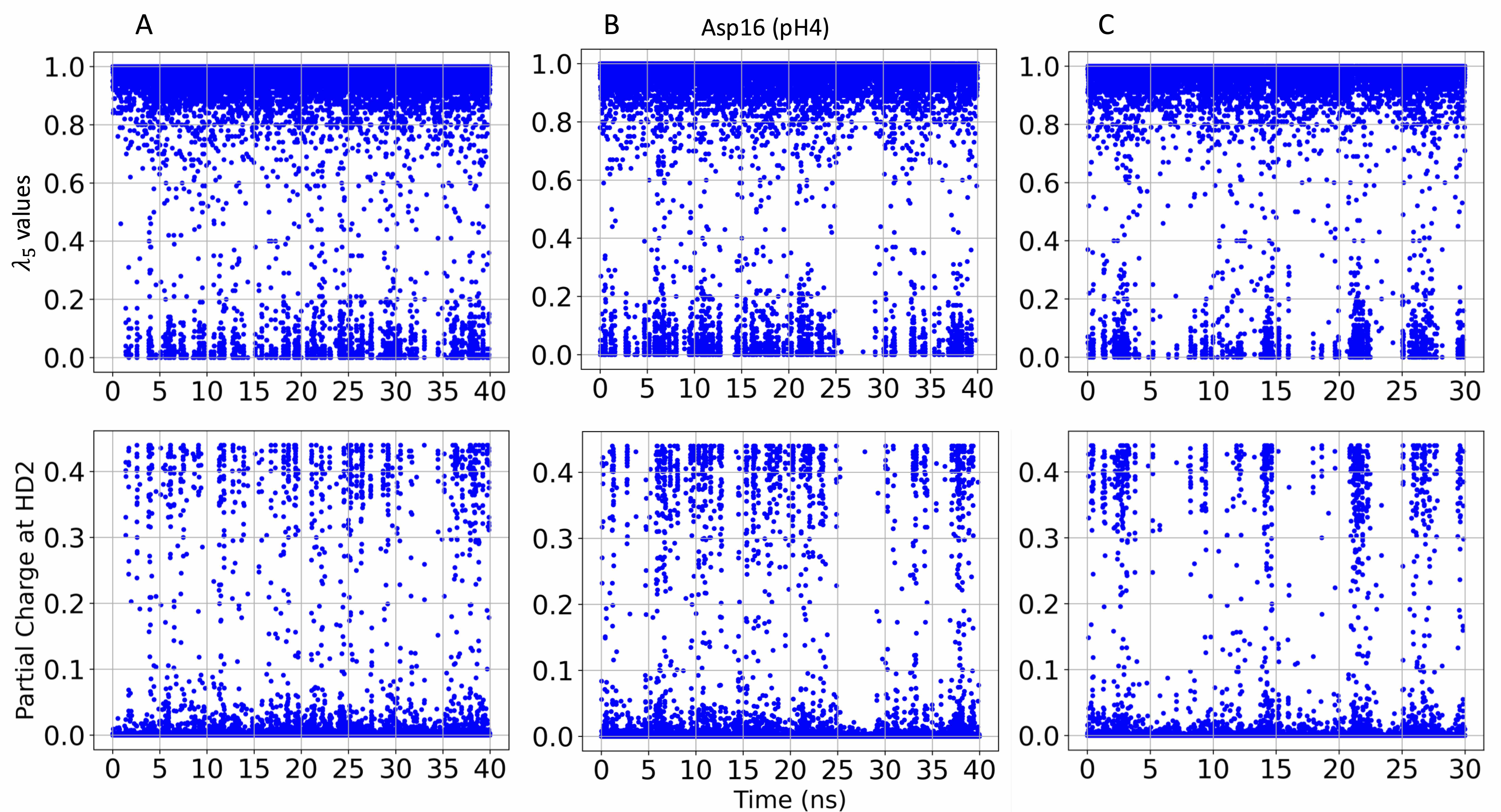

### S29.Fig

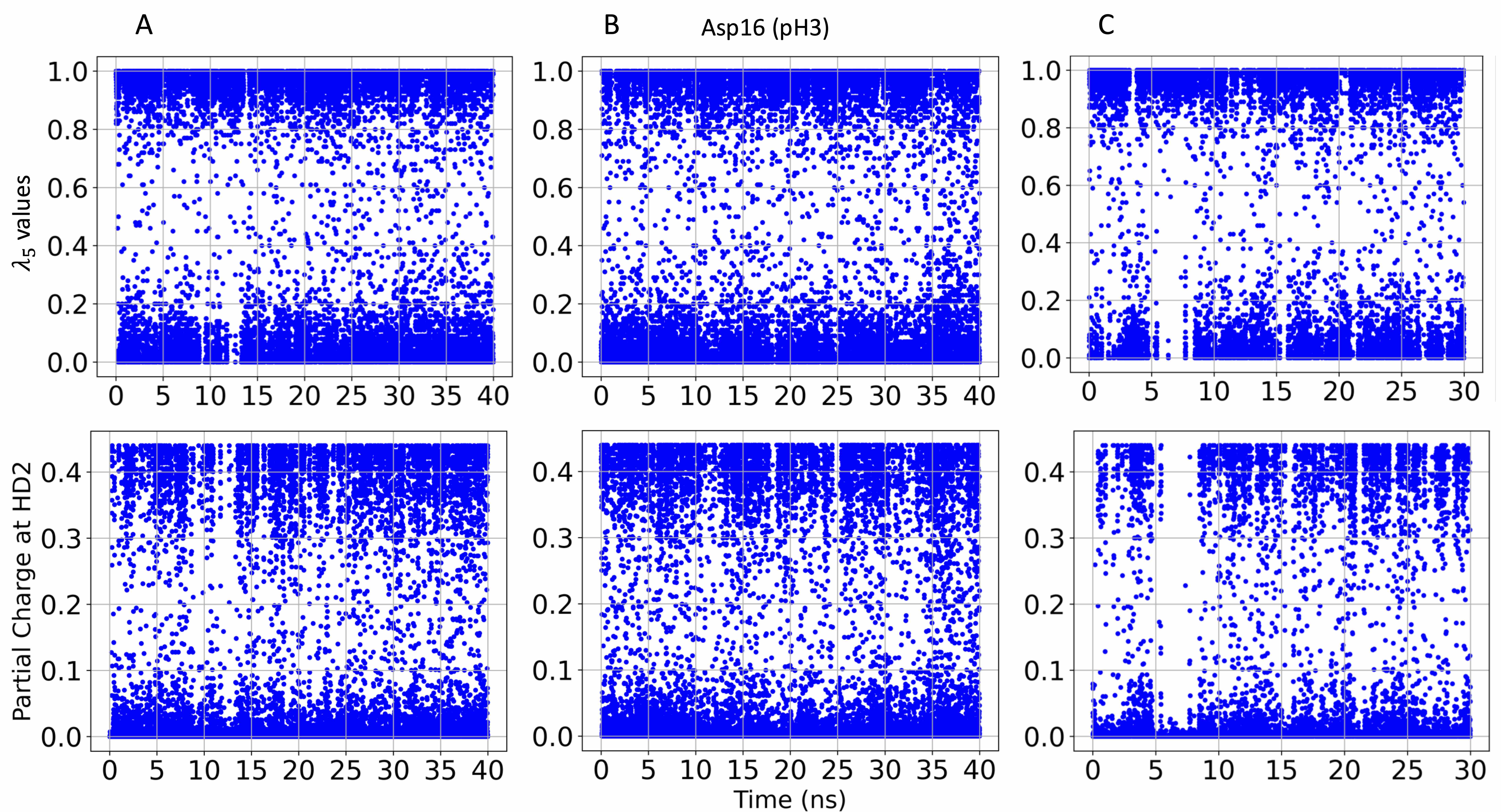

### S30.Fig

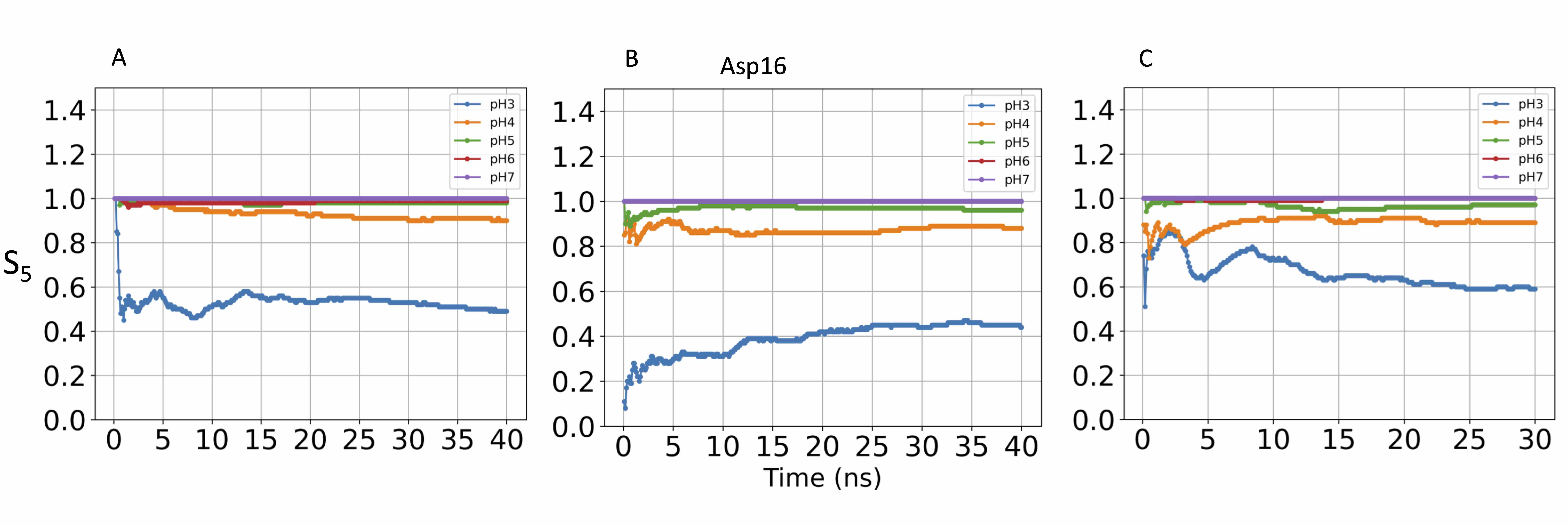

### S31.Fig

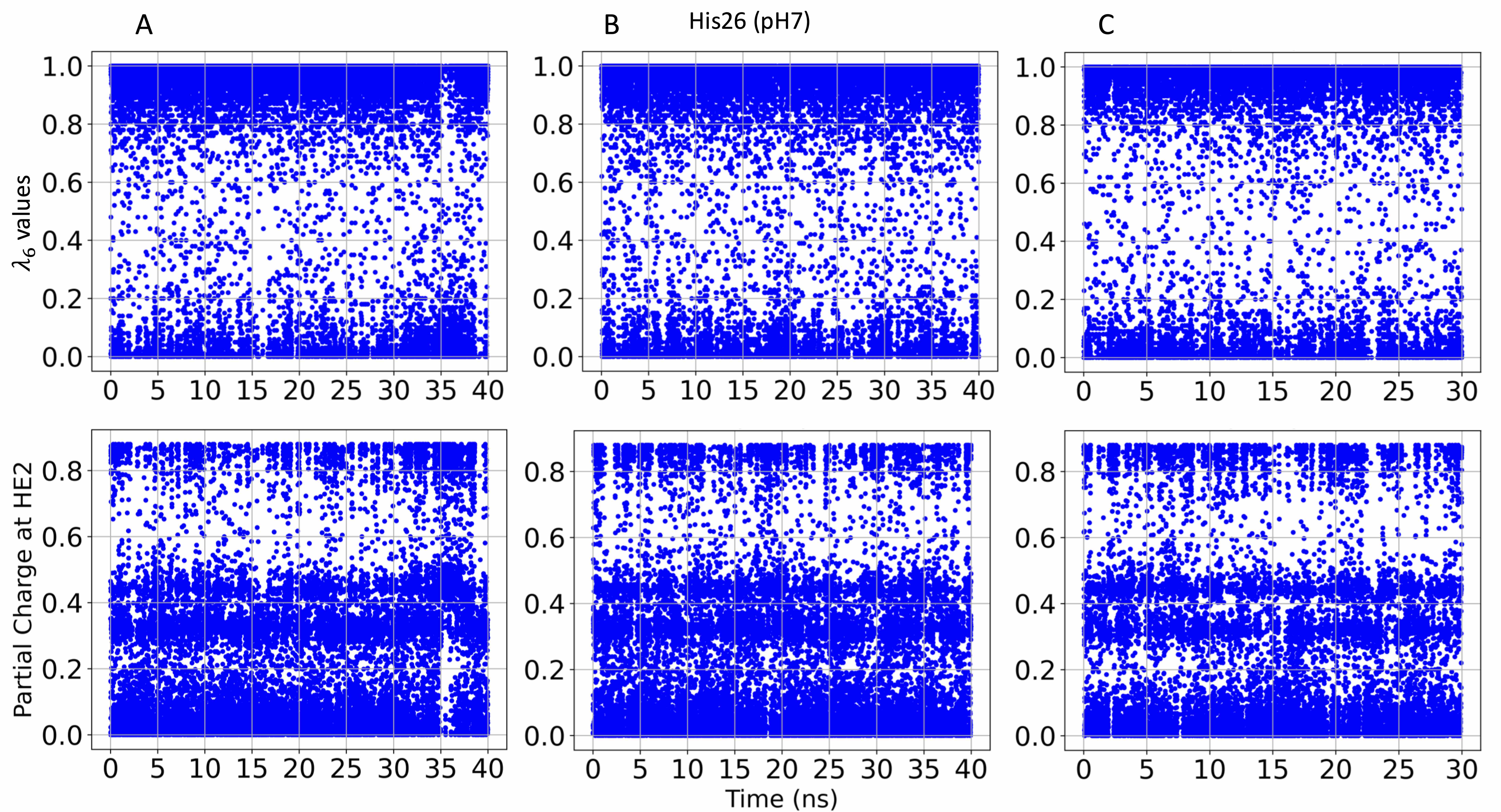

### S32.Fig

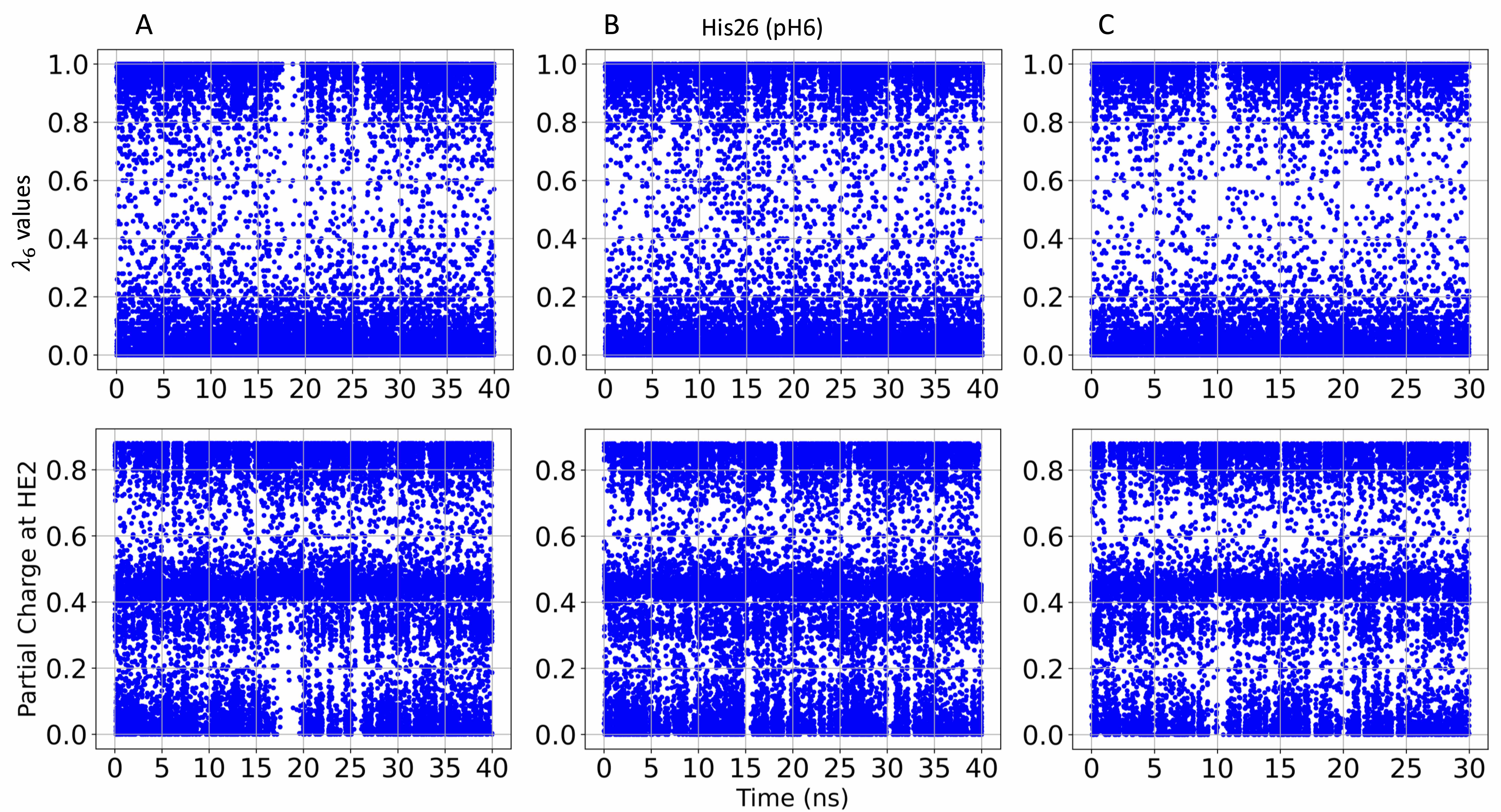

### S33.Fig

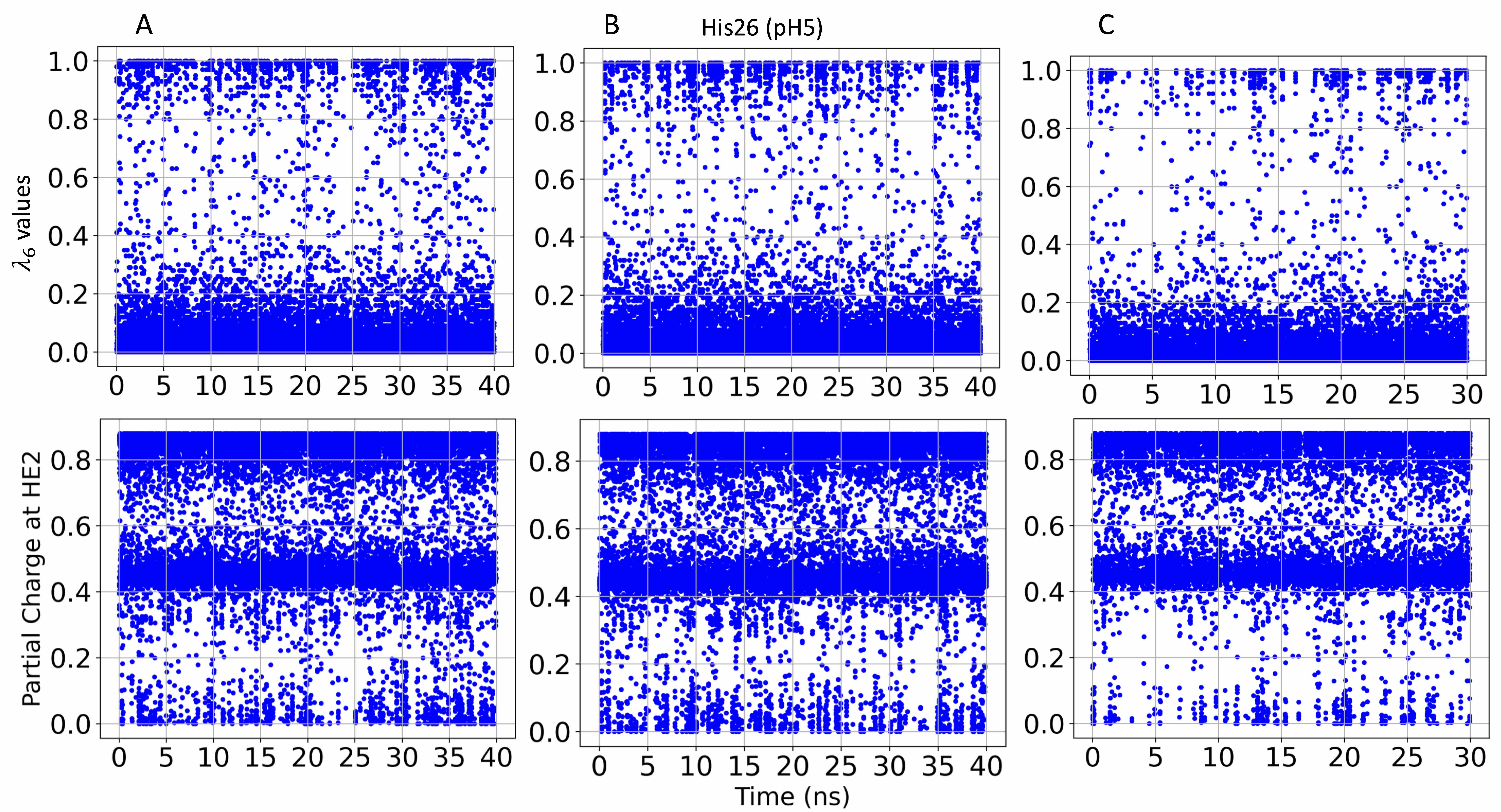

### S34.Fig

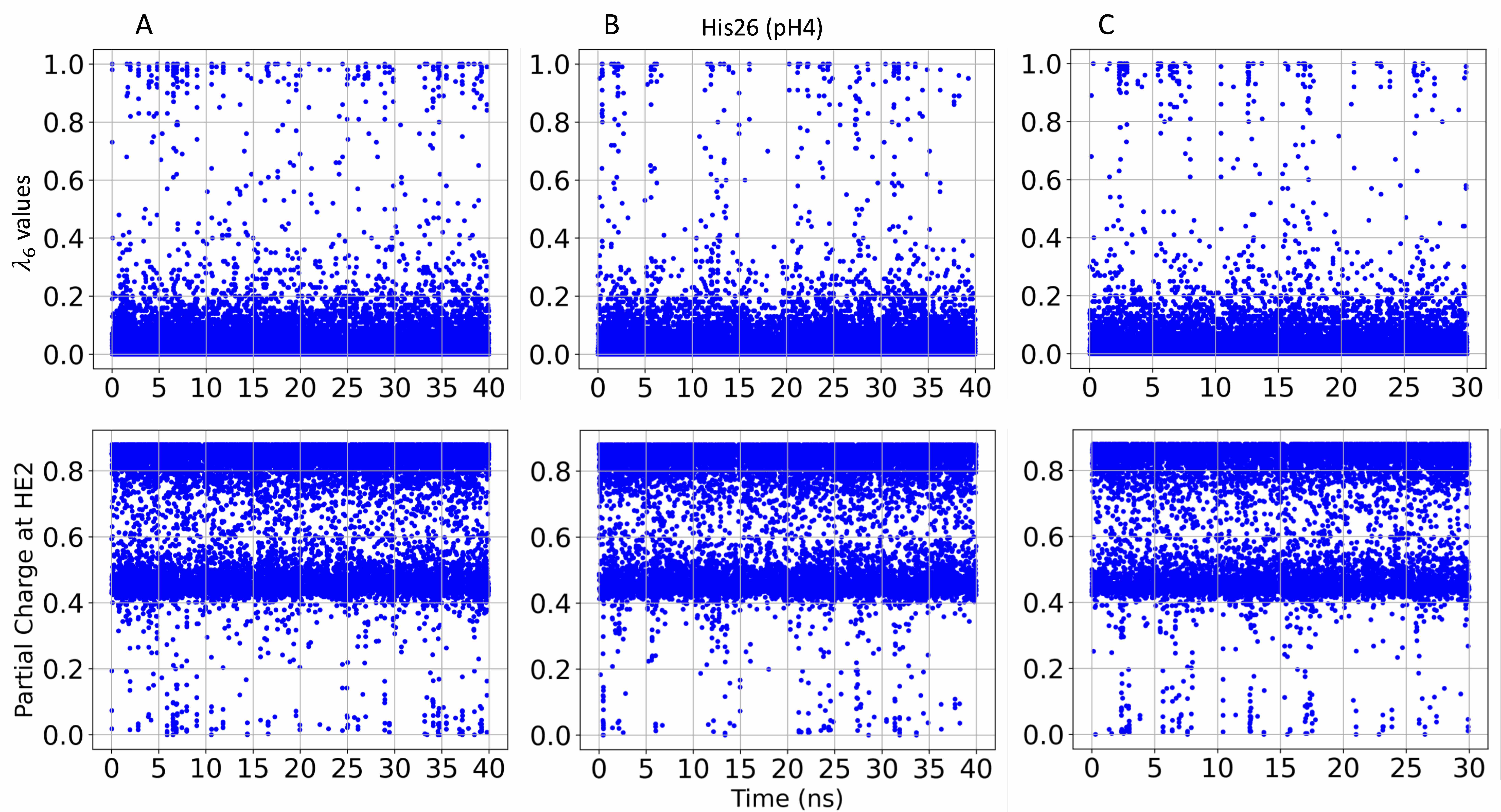

### S35.Fig

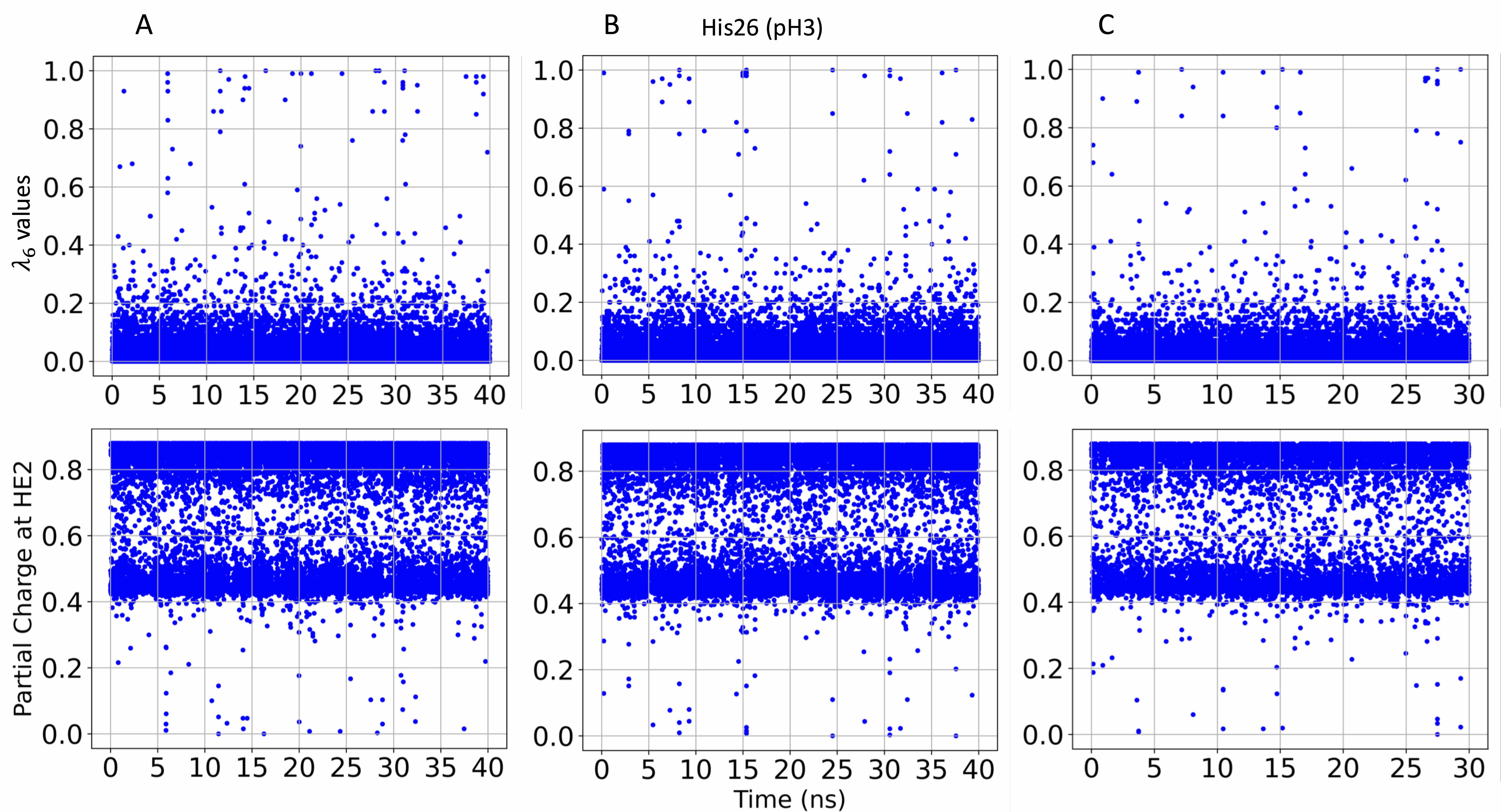

### S36.Fig

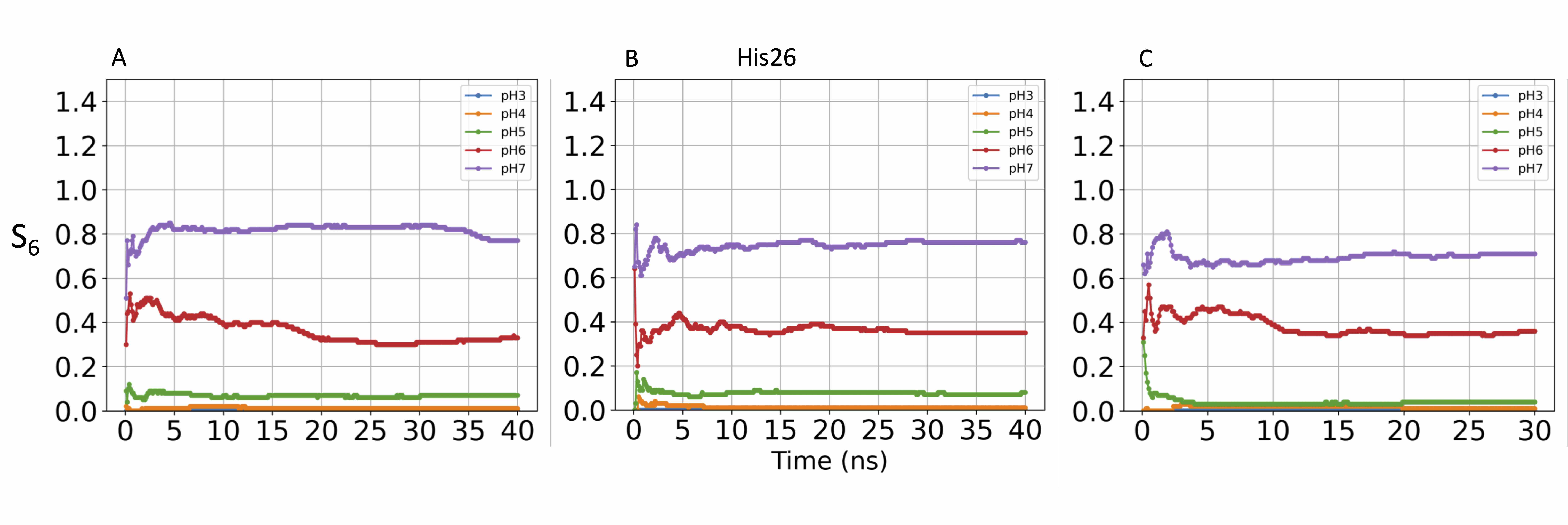

### S37.Fig

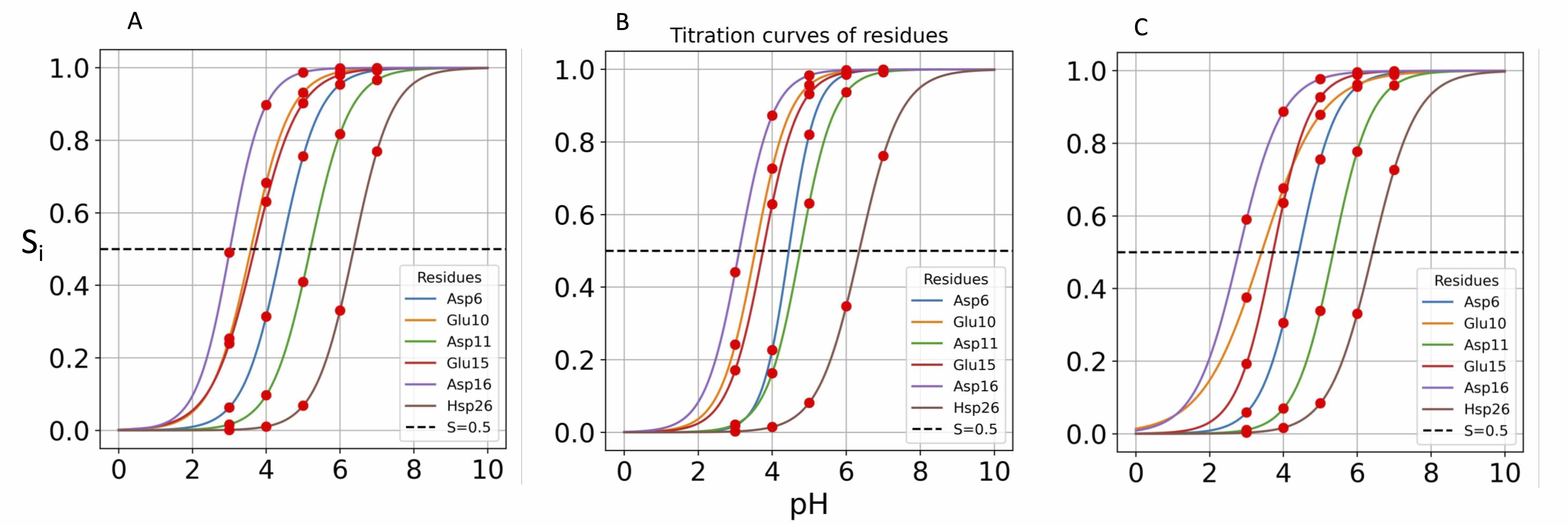

### S38.Fig

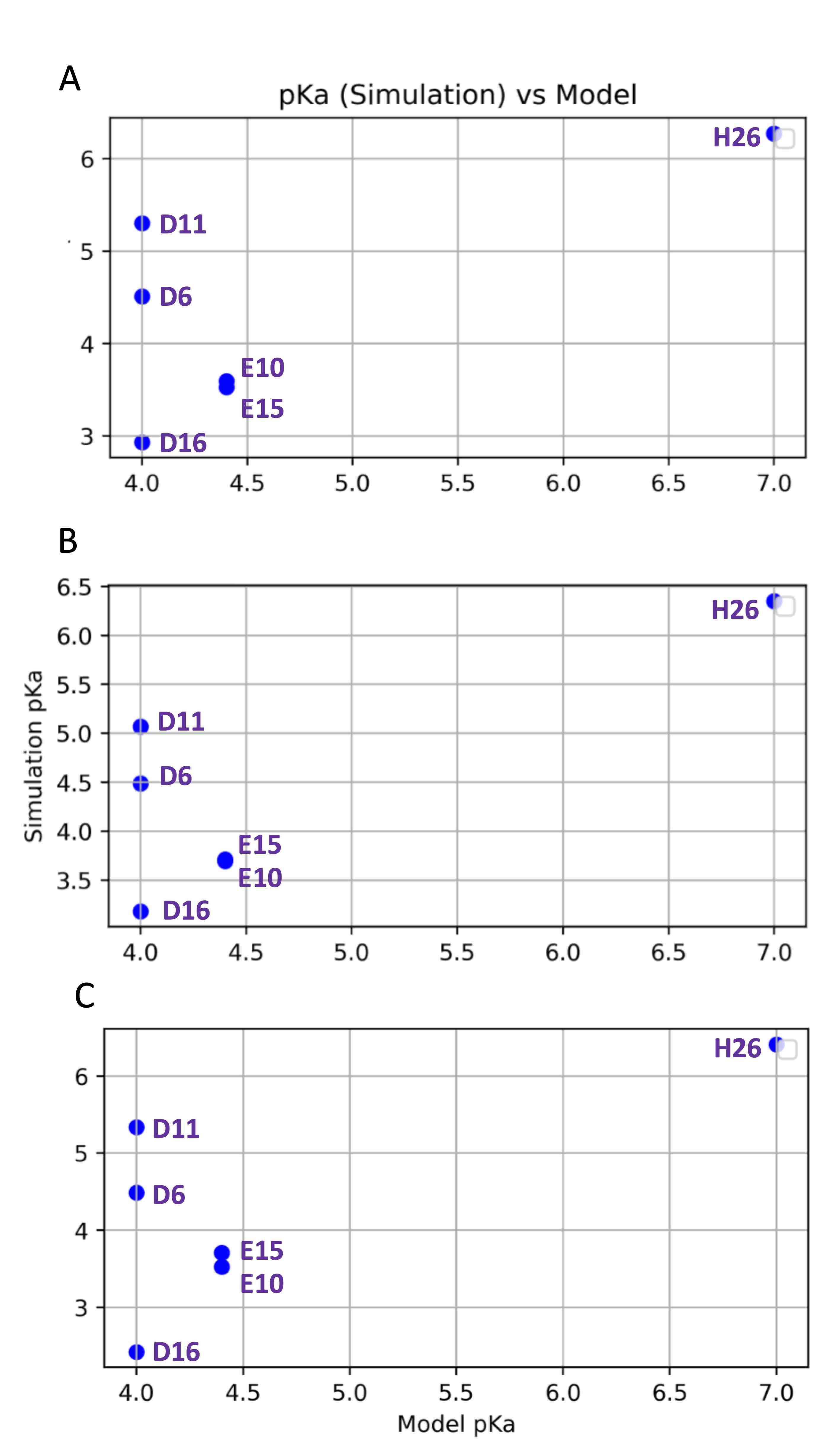

### S39.Fig

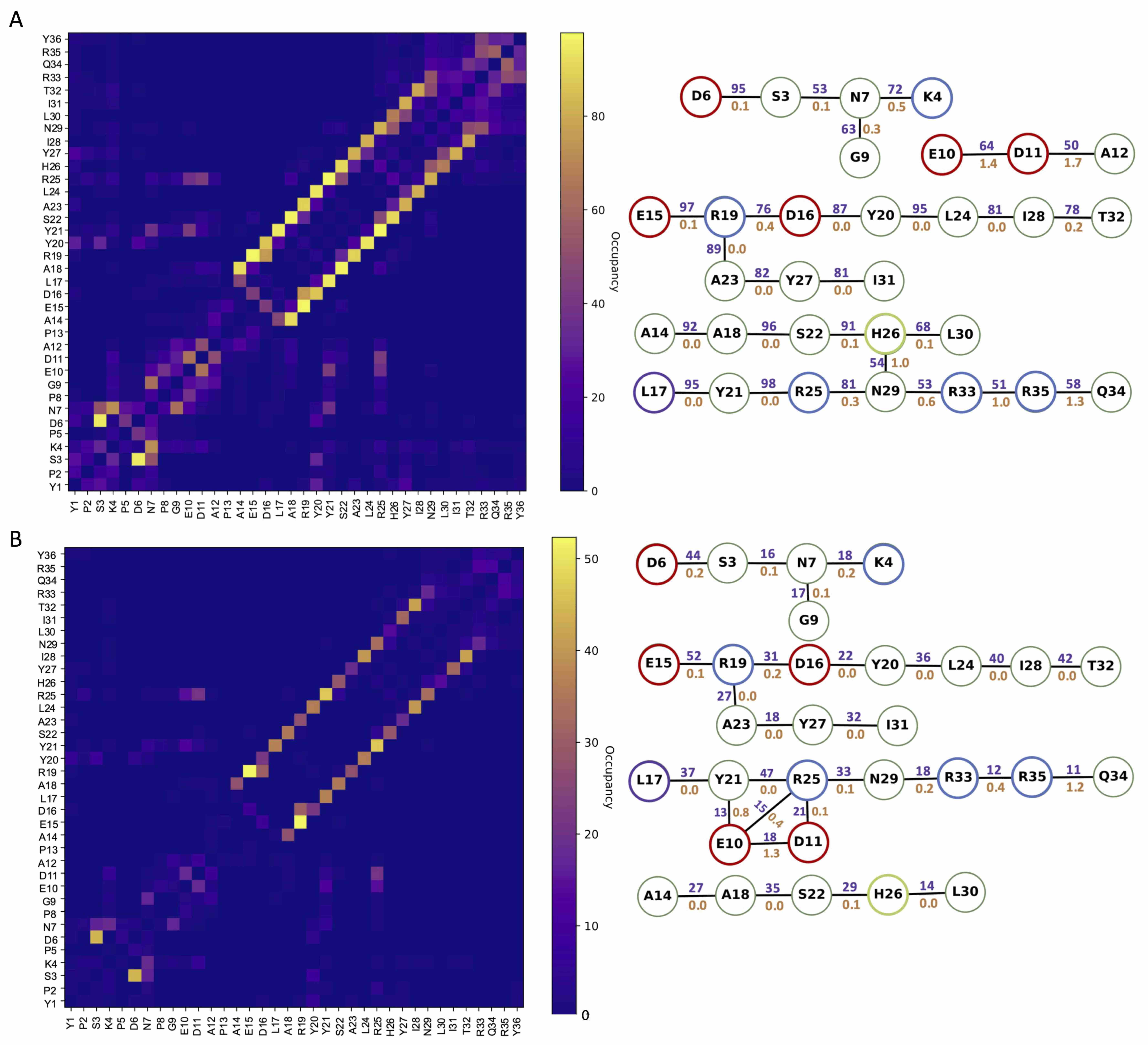

### S40.Fig

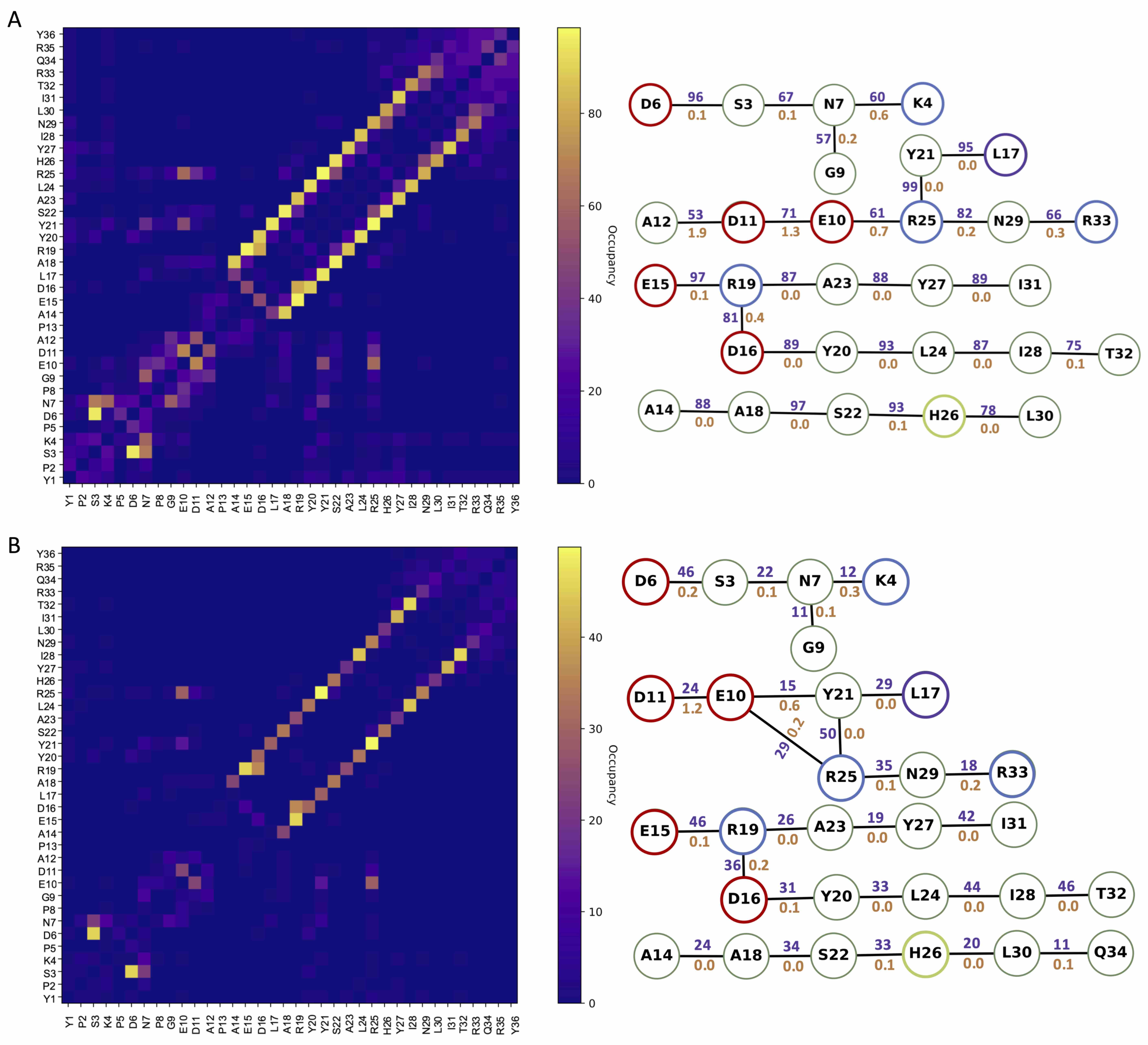

### S41.Fig

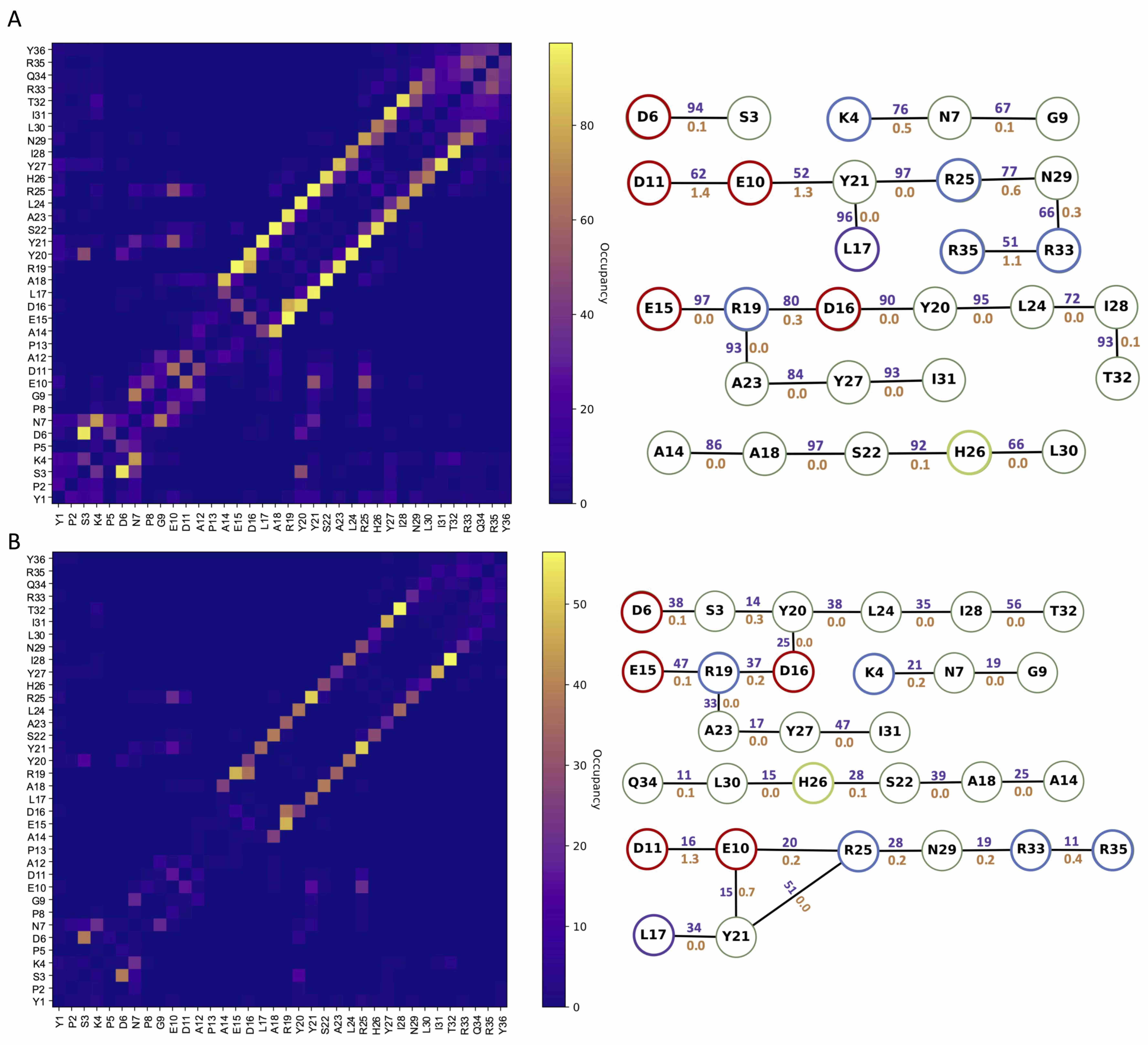

### S42.Fig

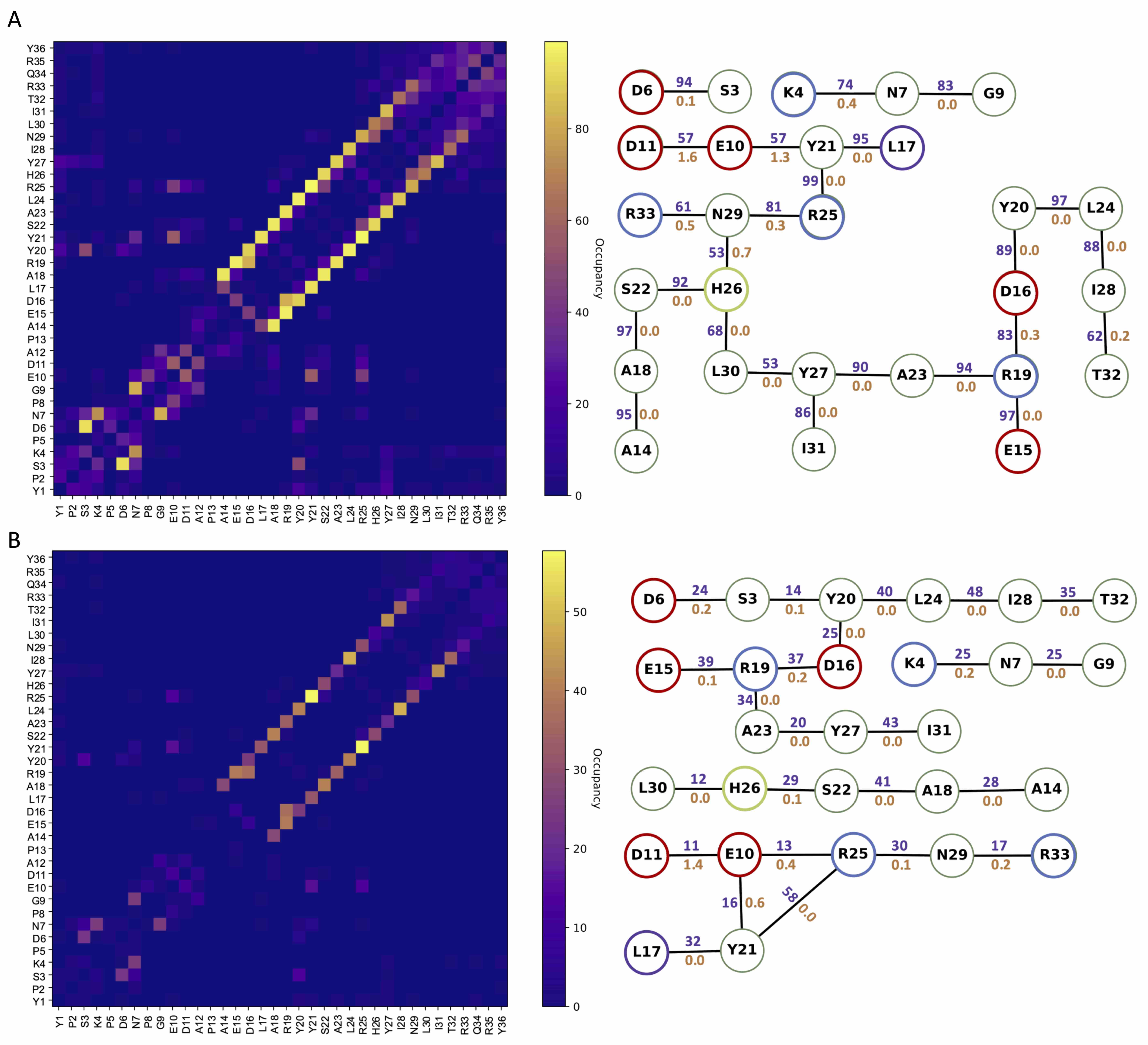

### S43.Fig

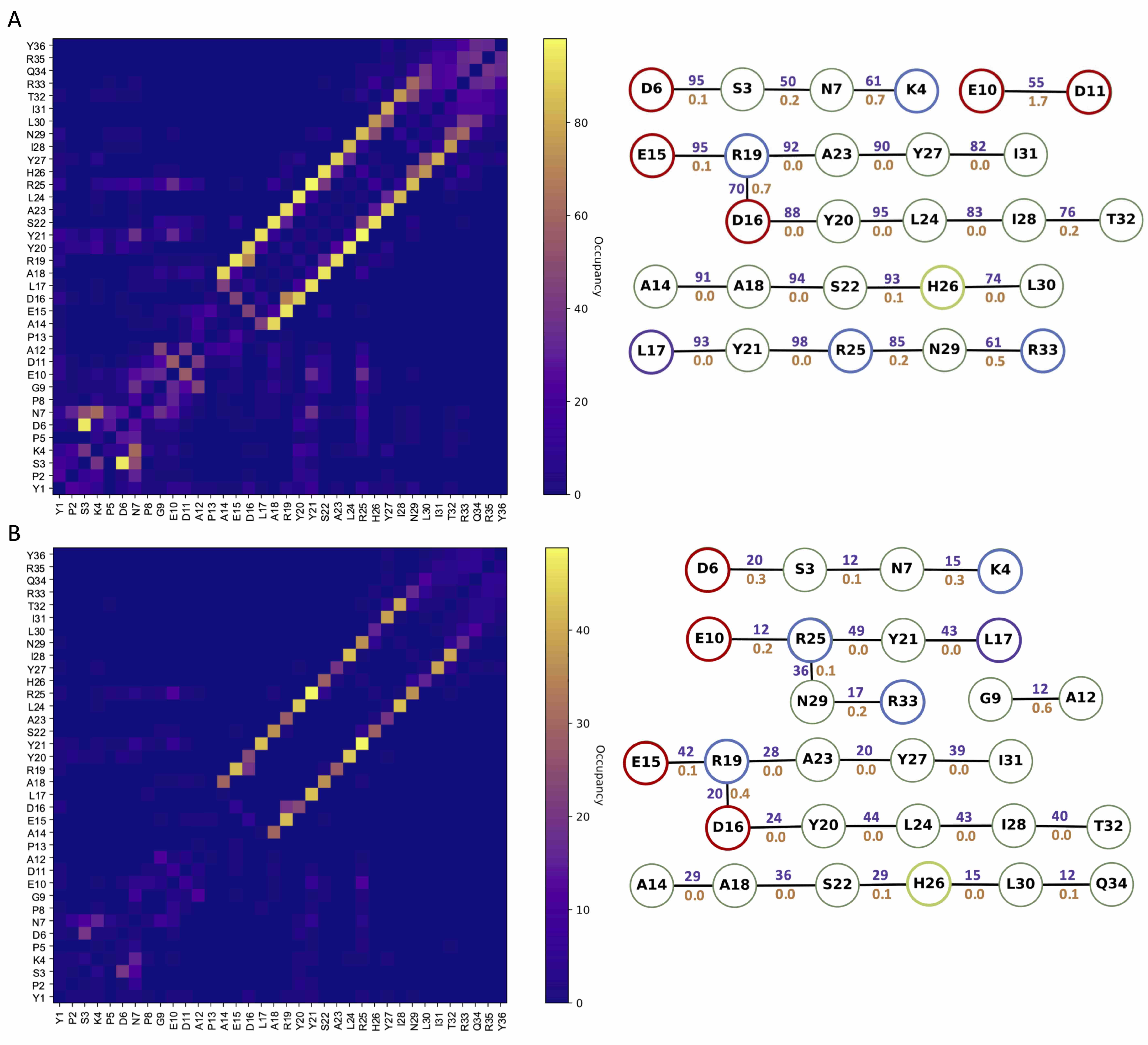

### S44.Fig

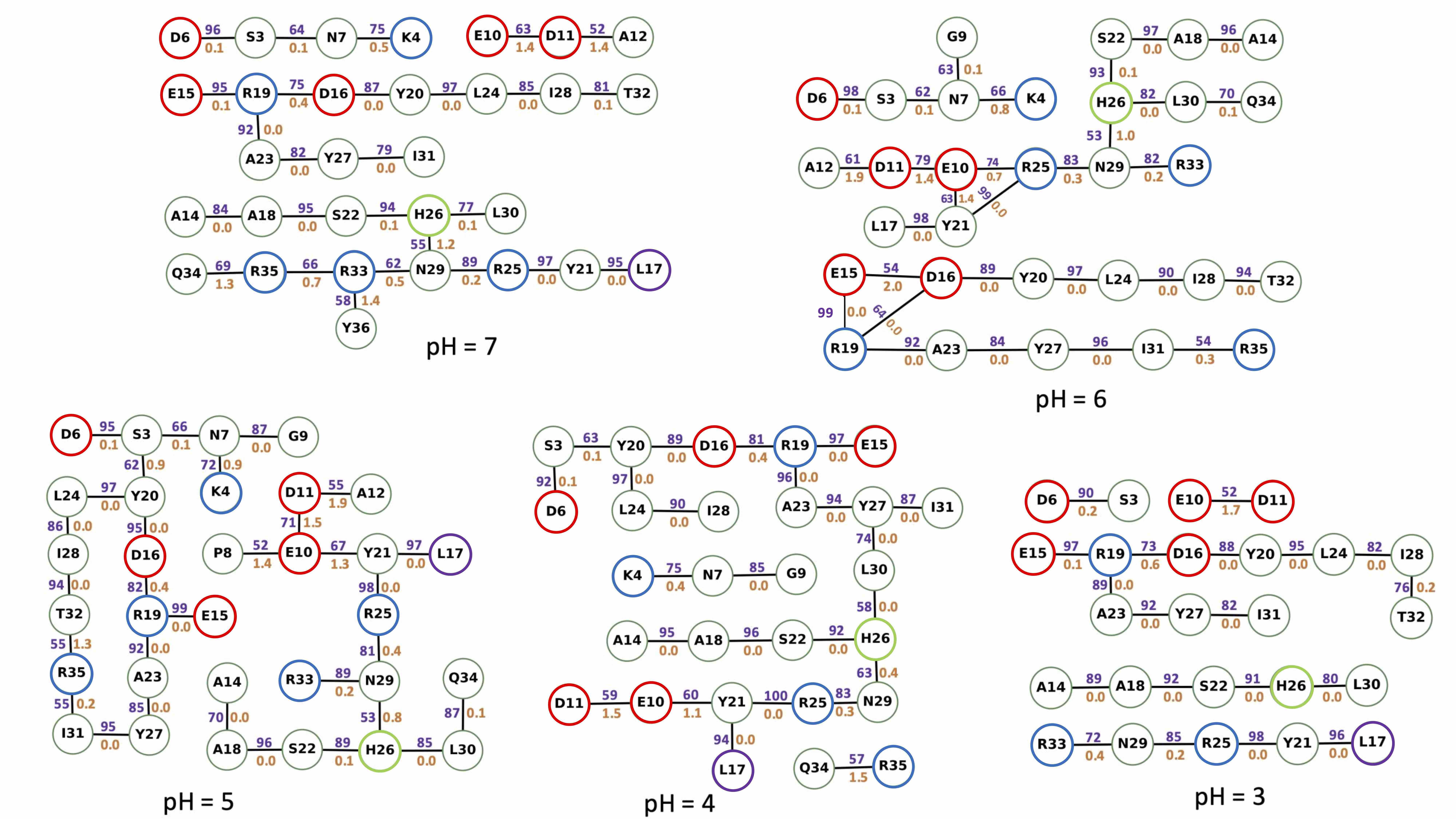

### S45.Fig

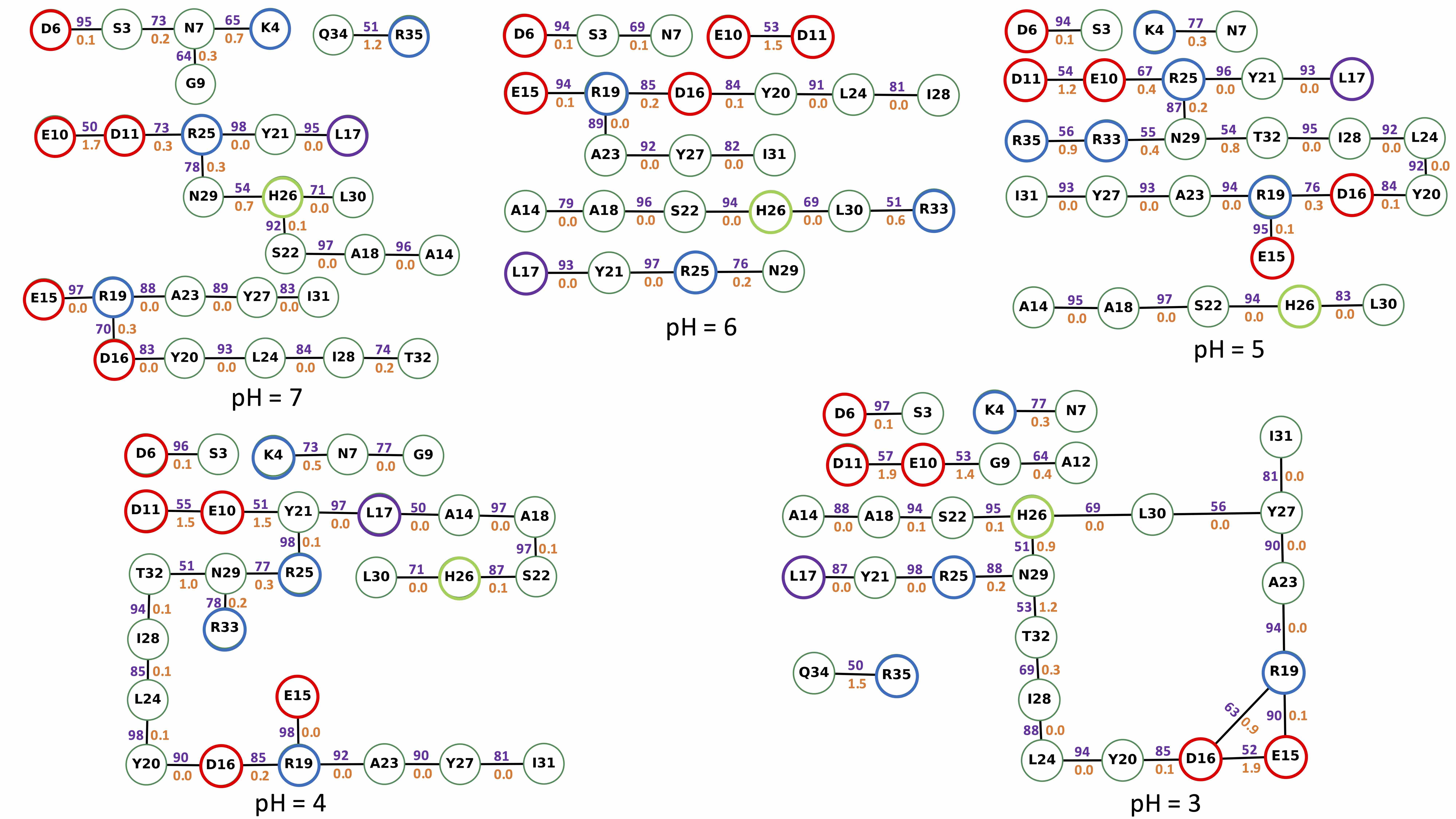

### S46.Fig

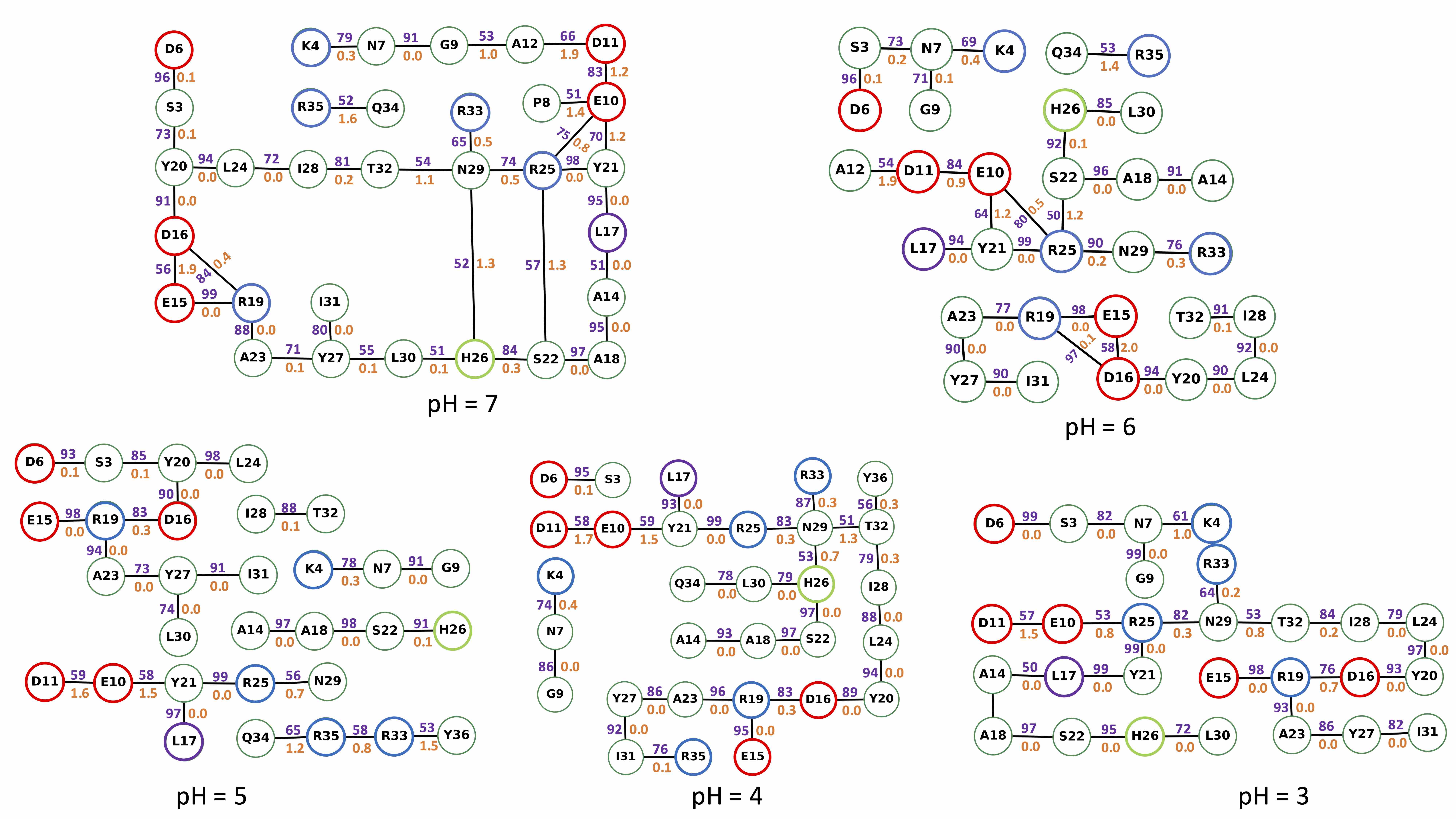

### S47.Fig

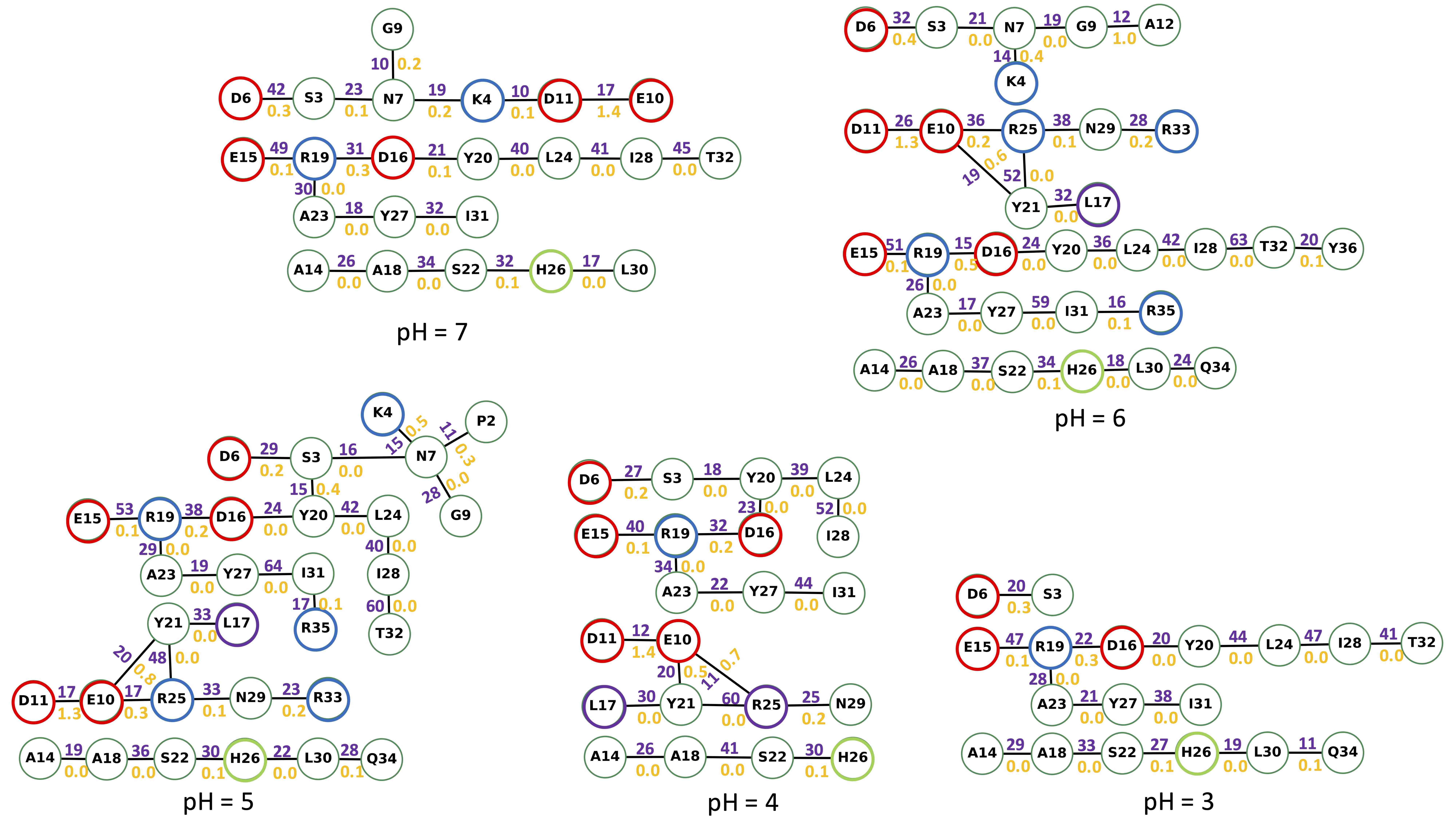

### S48.Fig

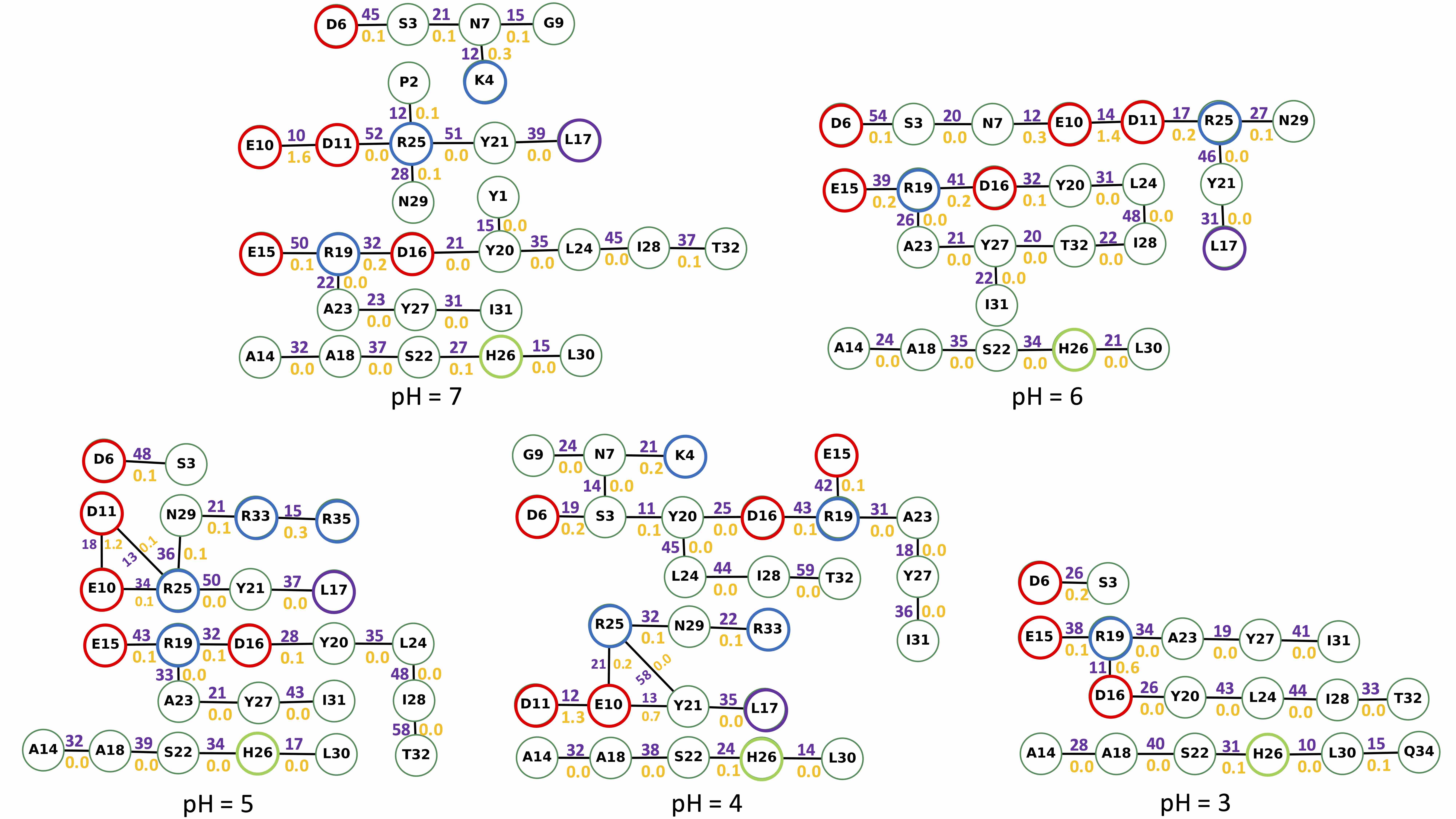

### S49.Fig

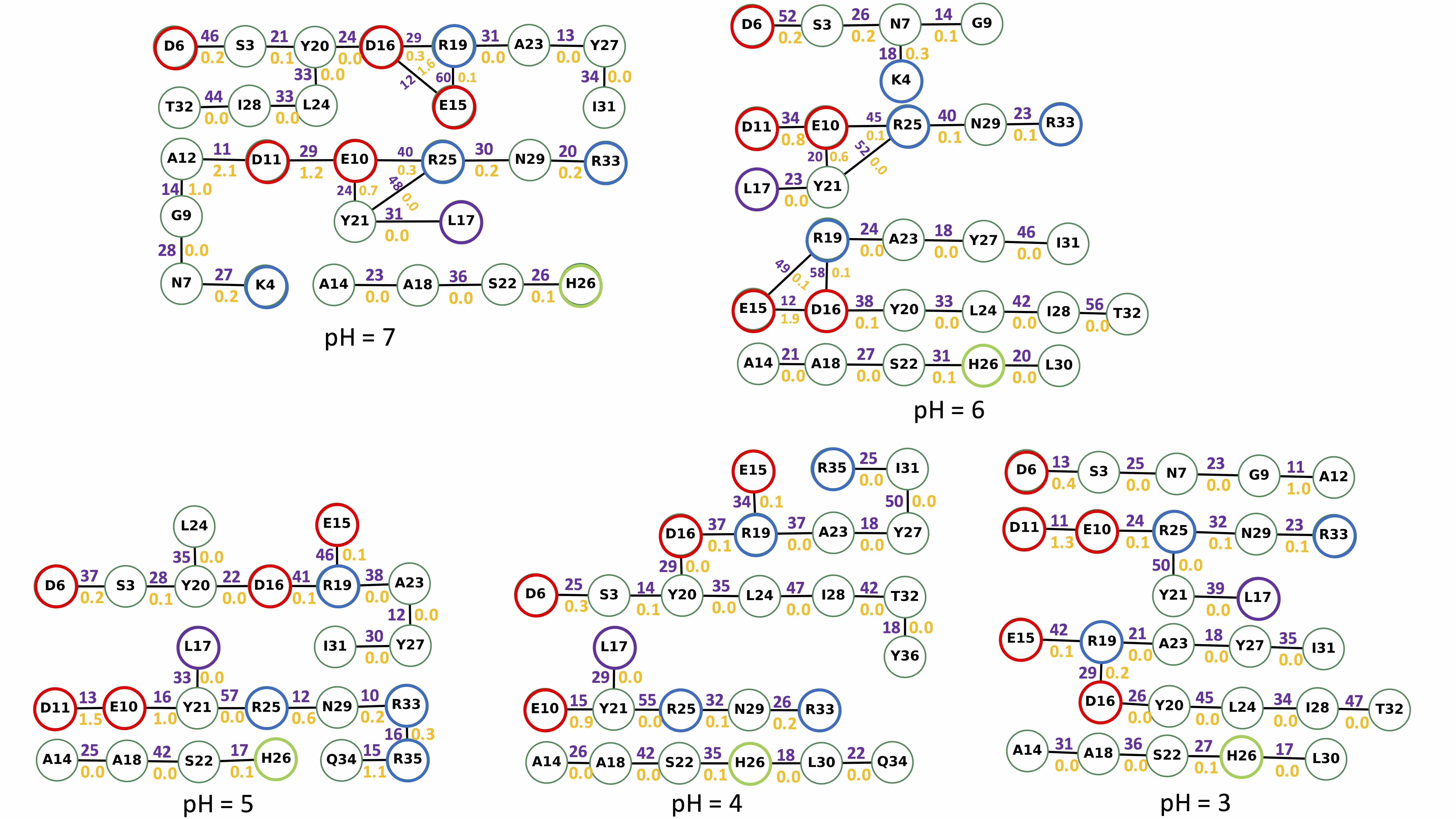

### S50.Fig

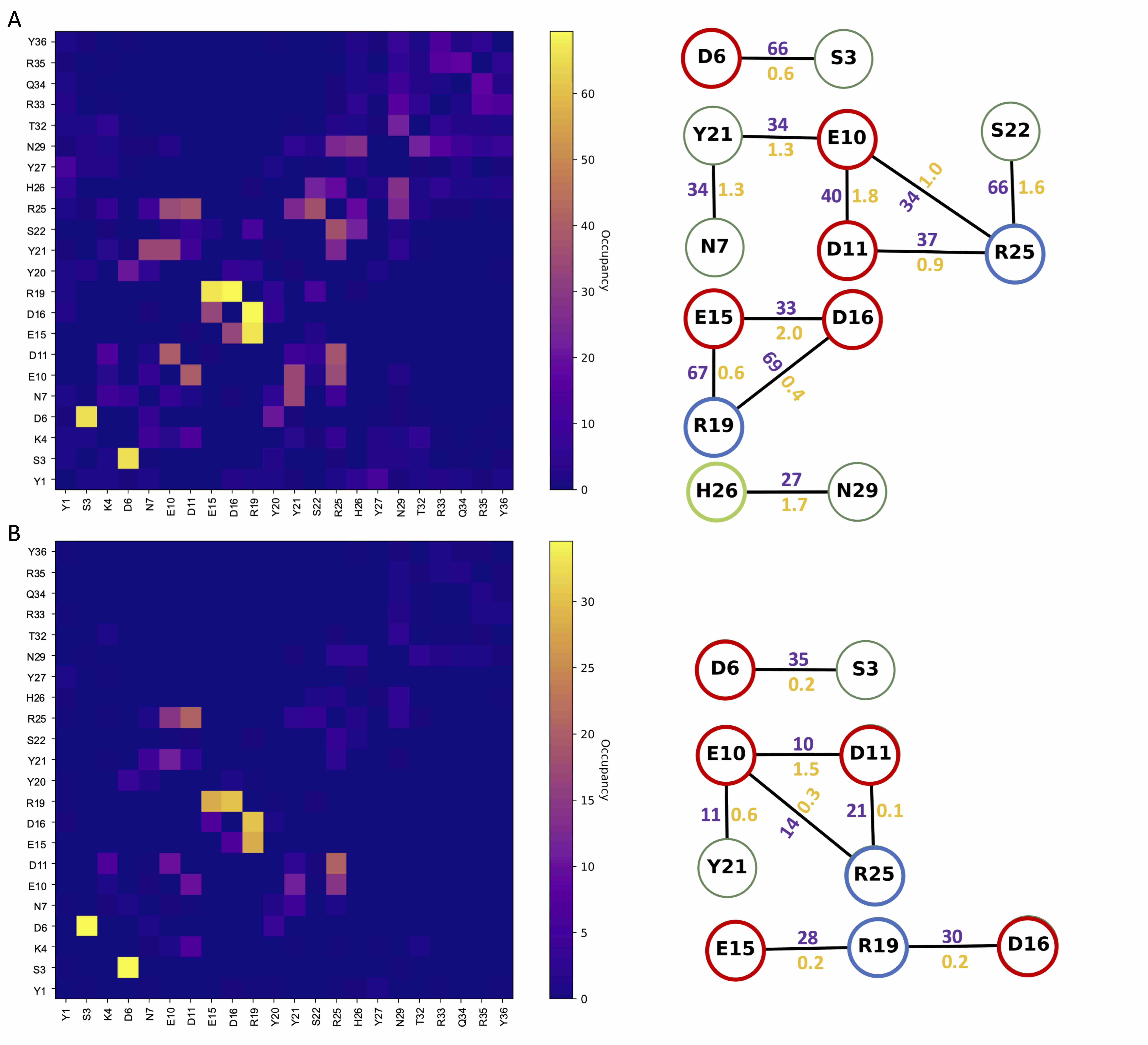

### S51.Fig

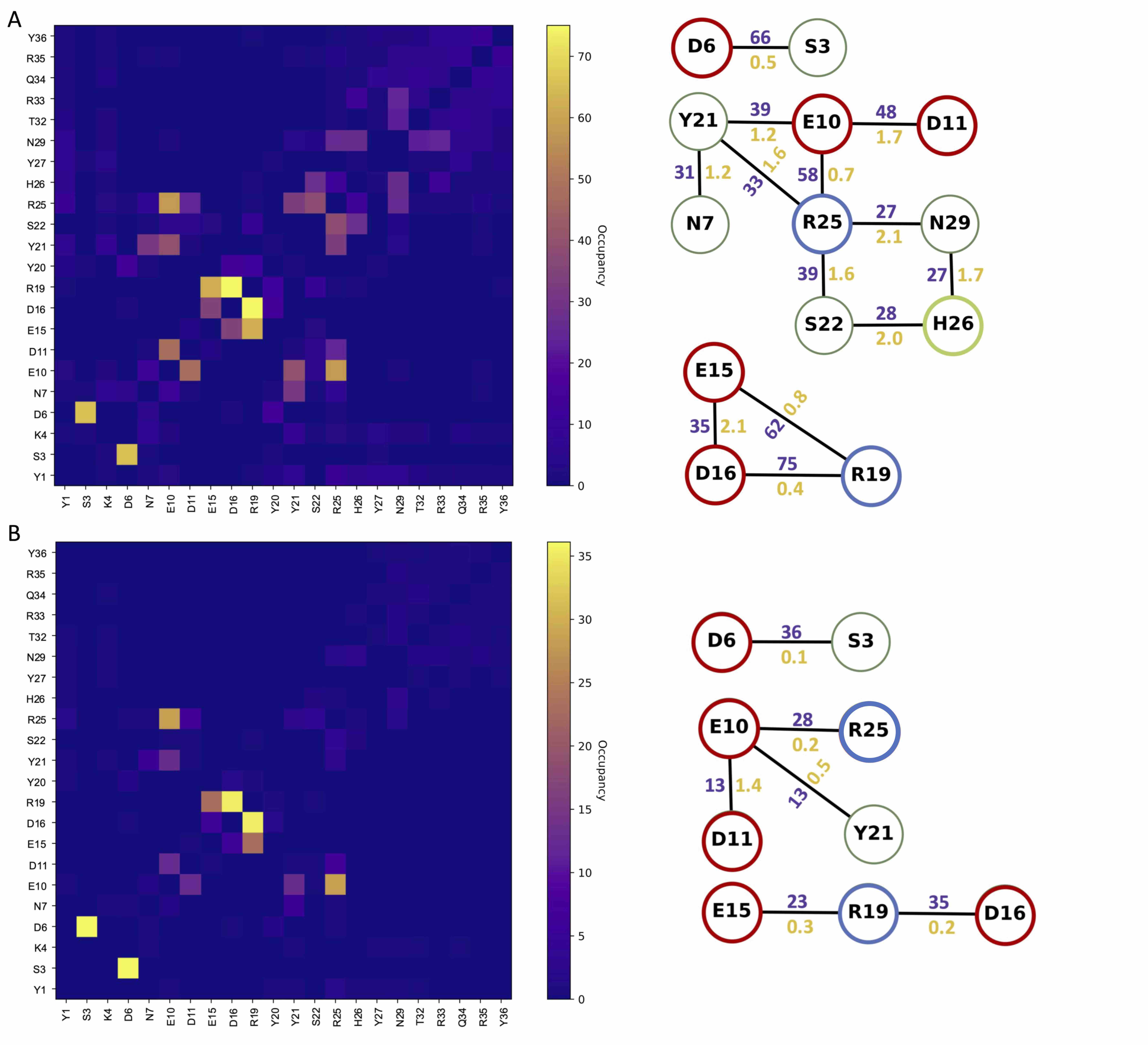

### S52.Fig

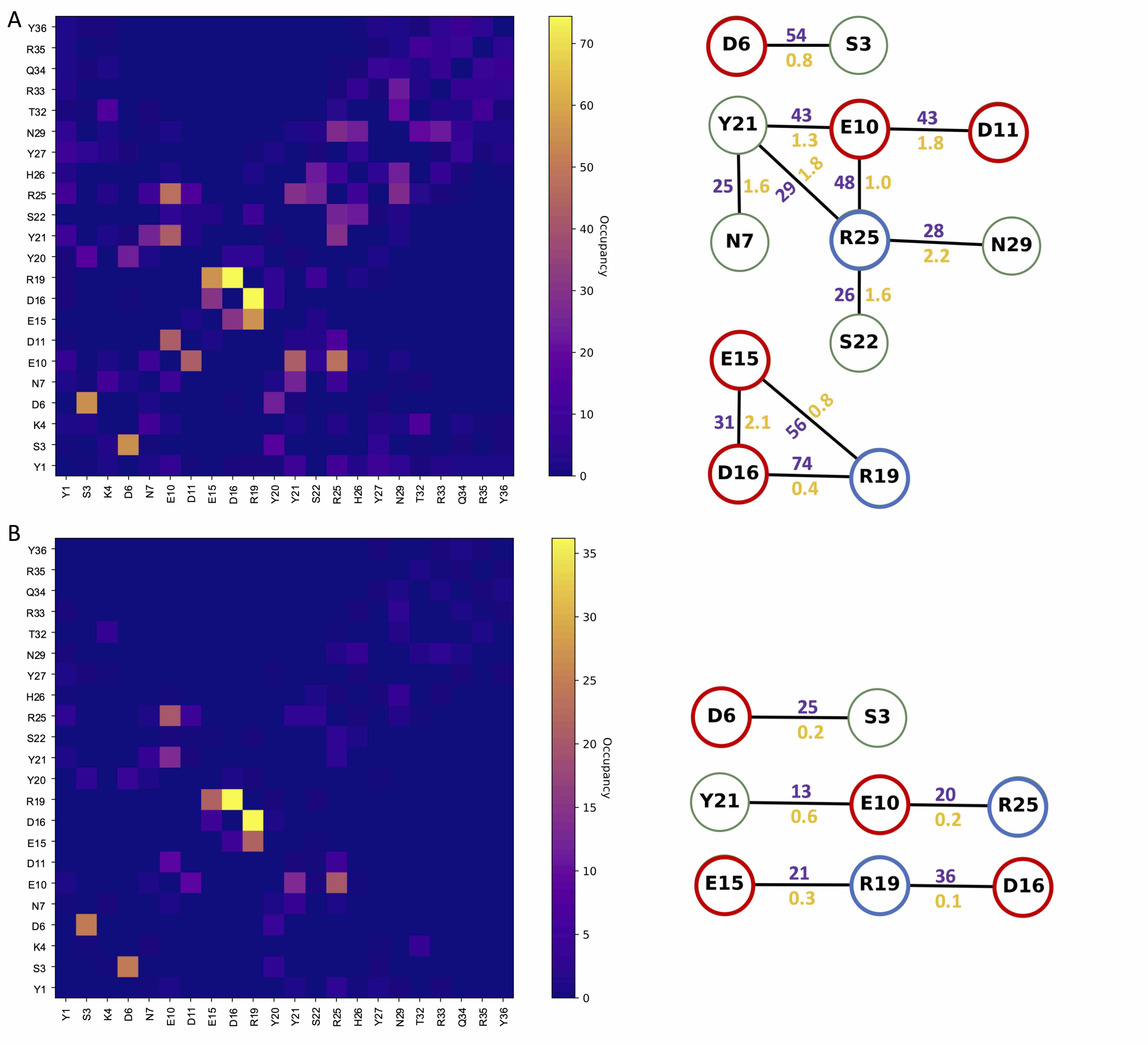

### S53.Fig

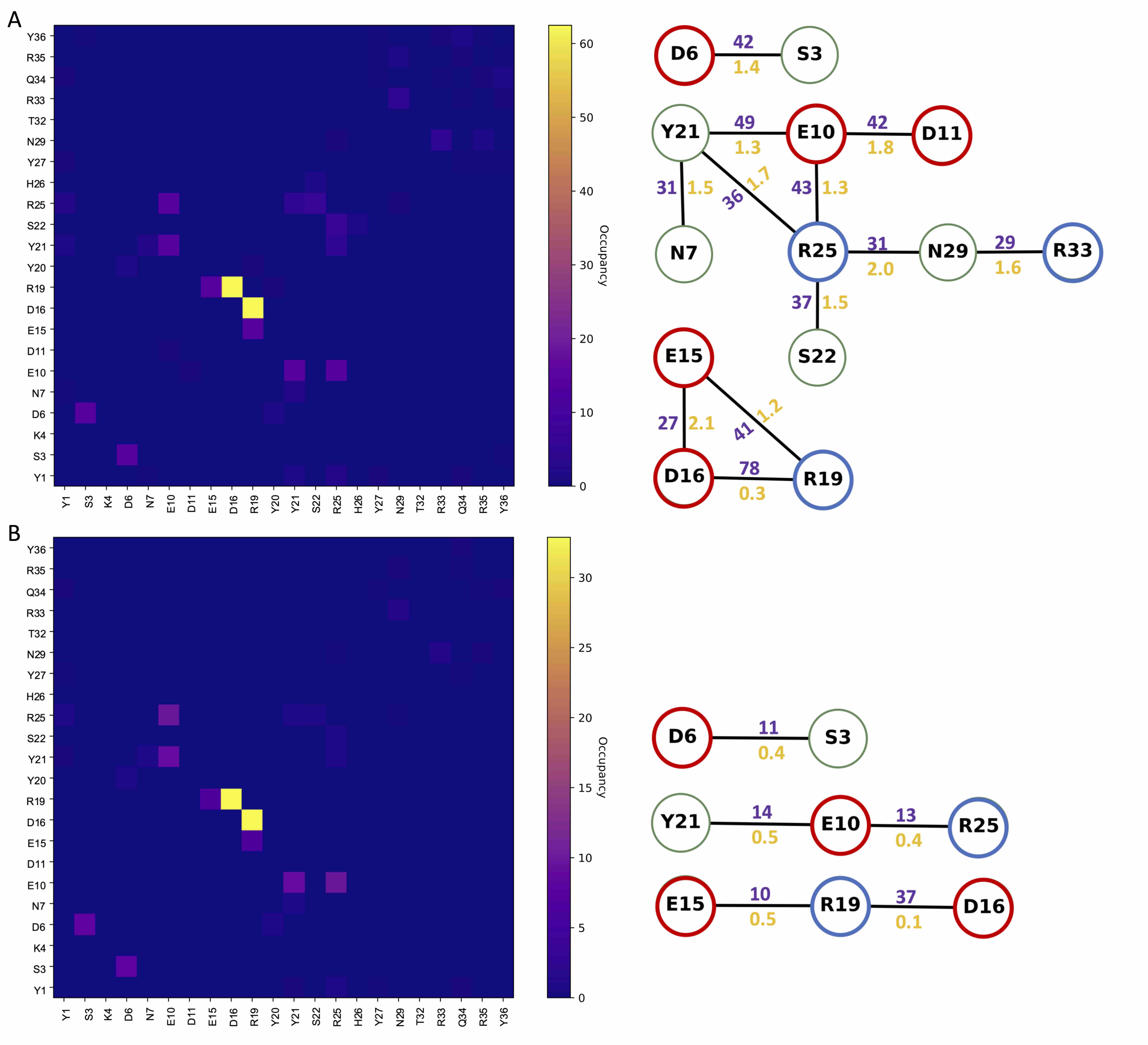

### S54.Fig

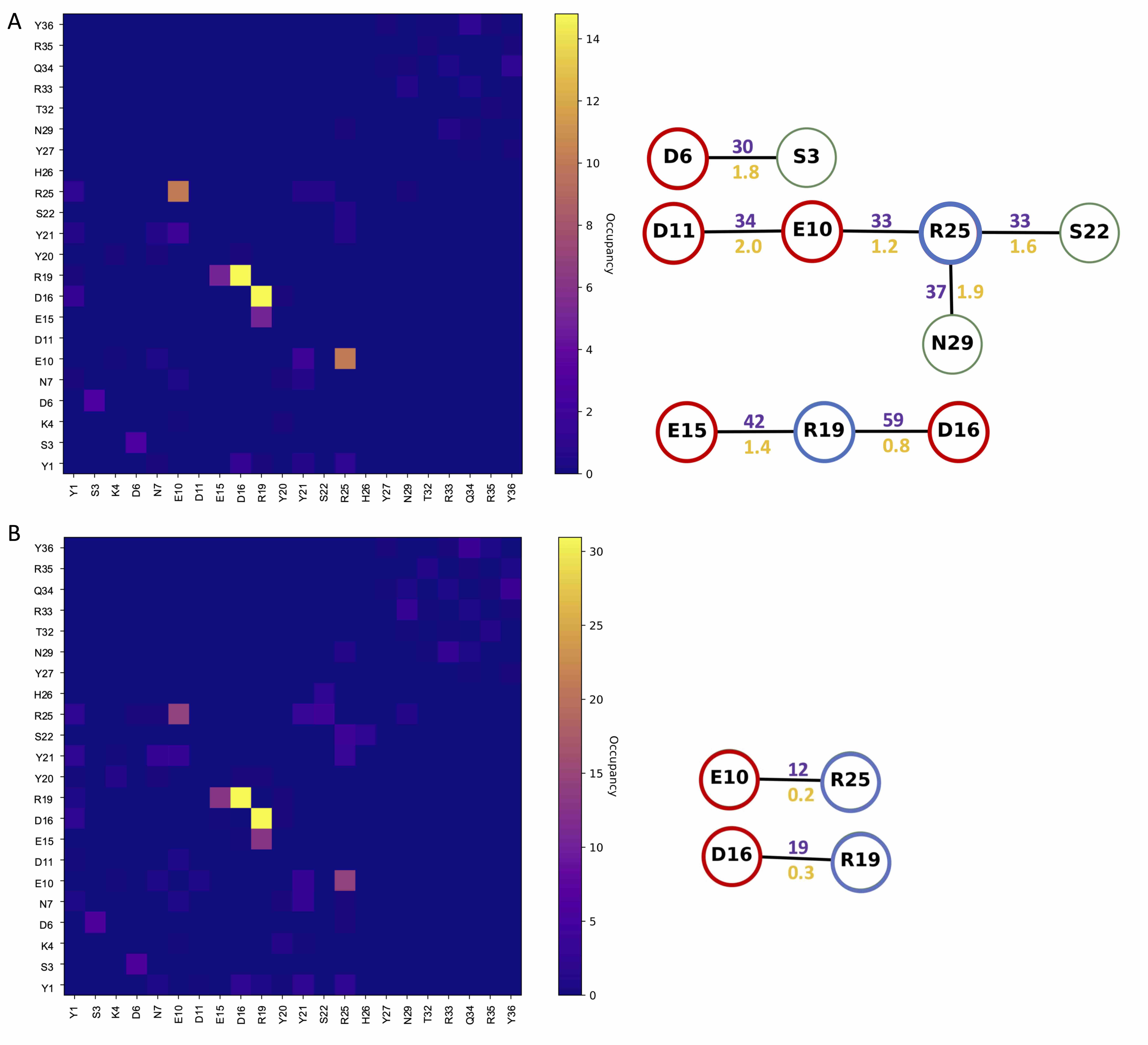

### S54.Fig

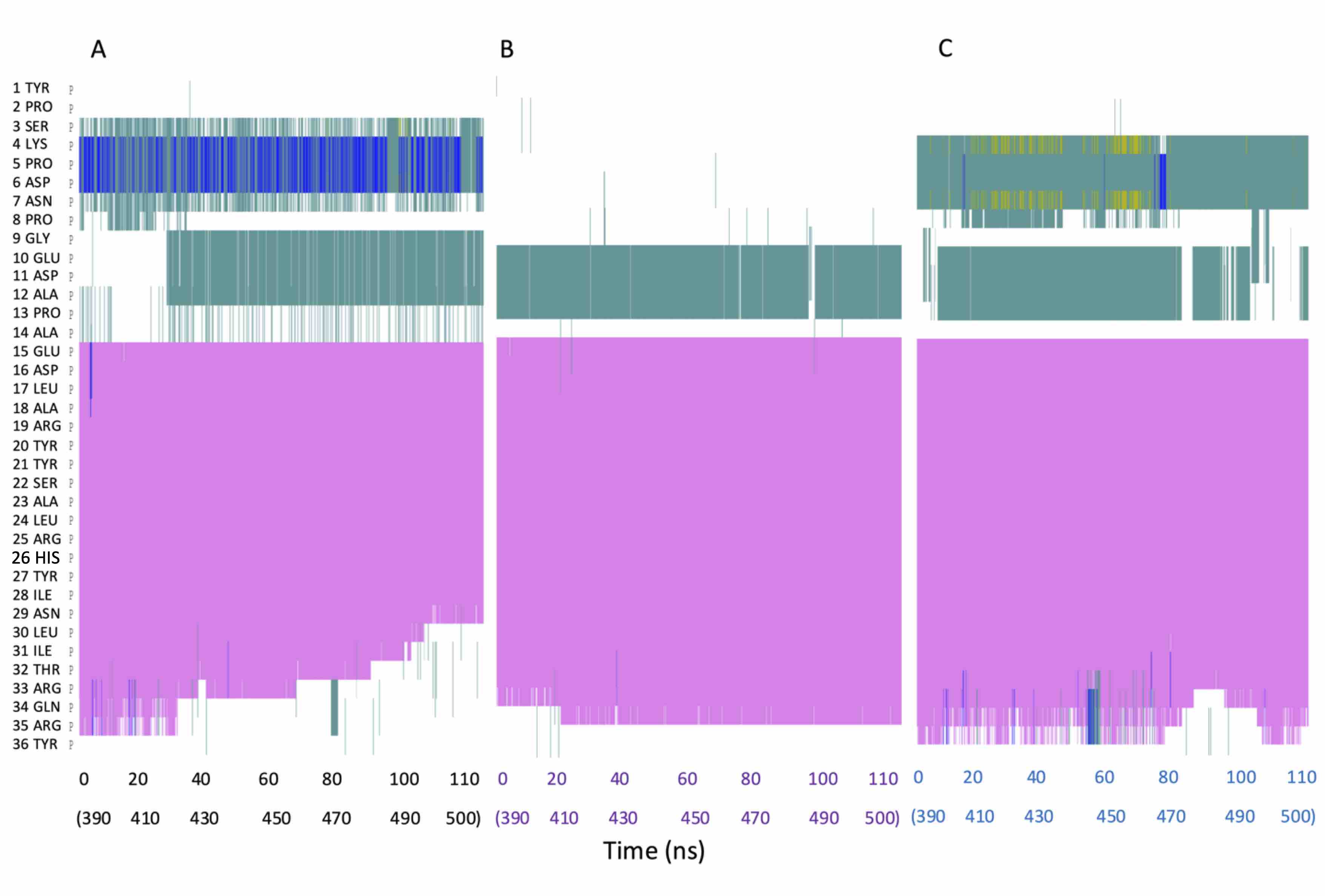

### S55.Fig

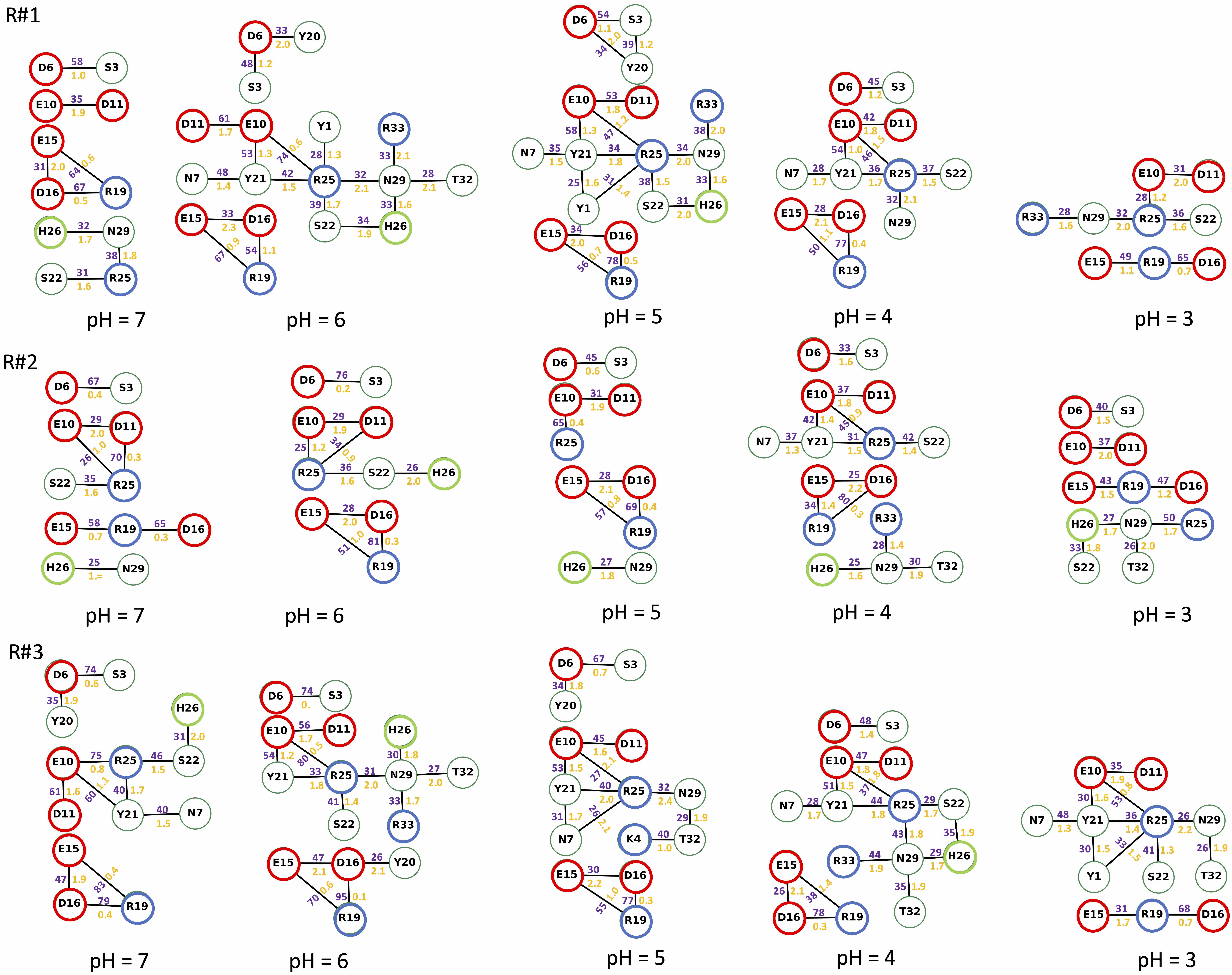

### S56.Fig

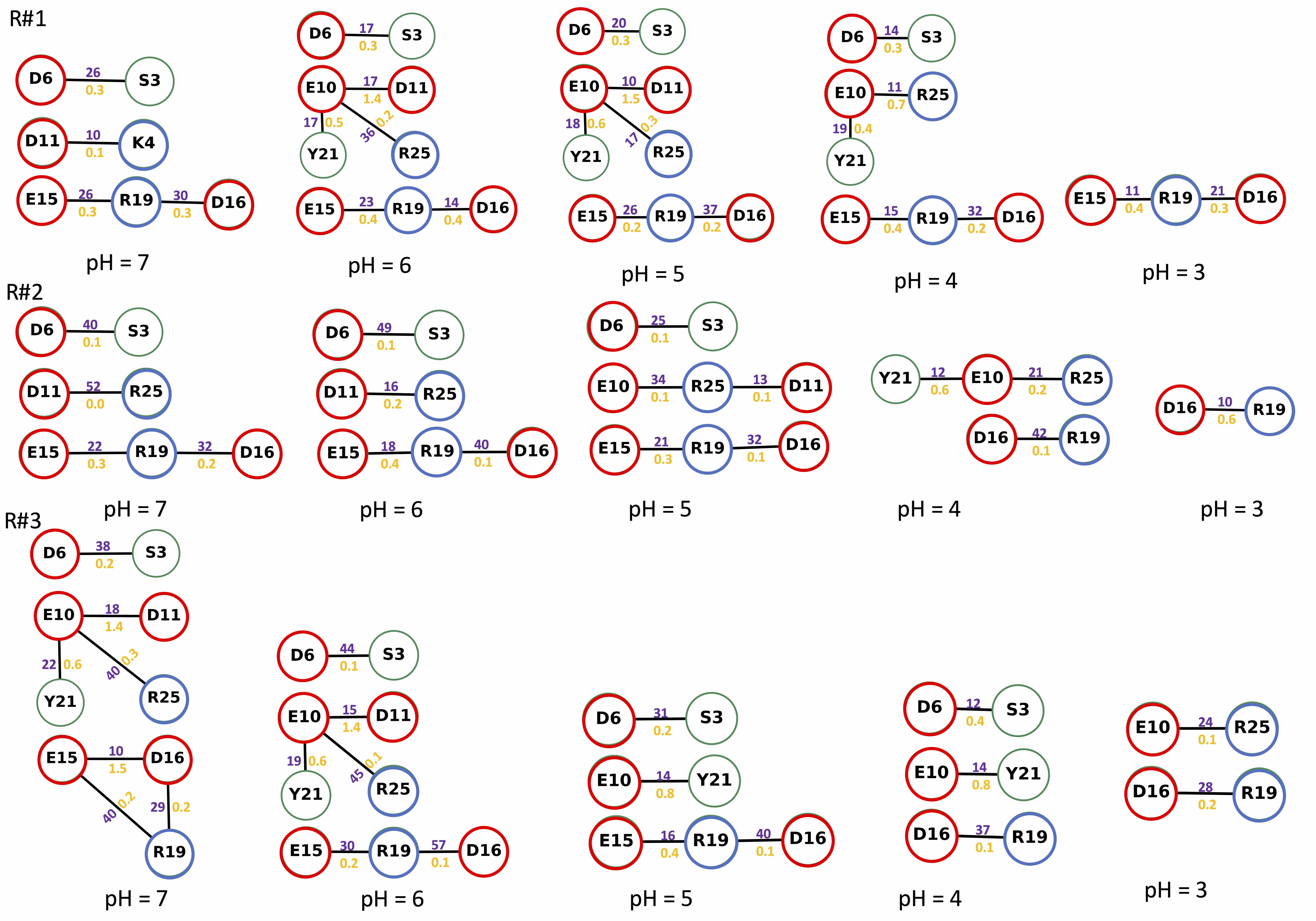

### S57.Fig

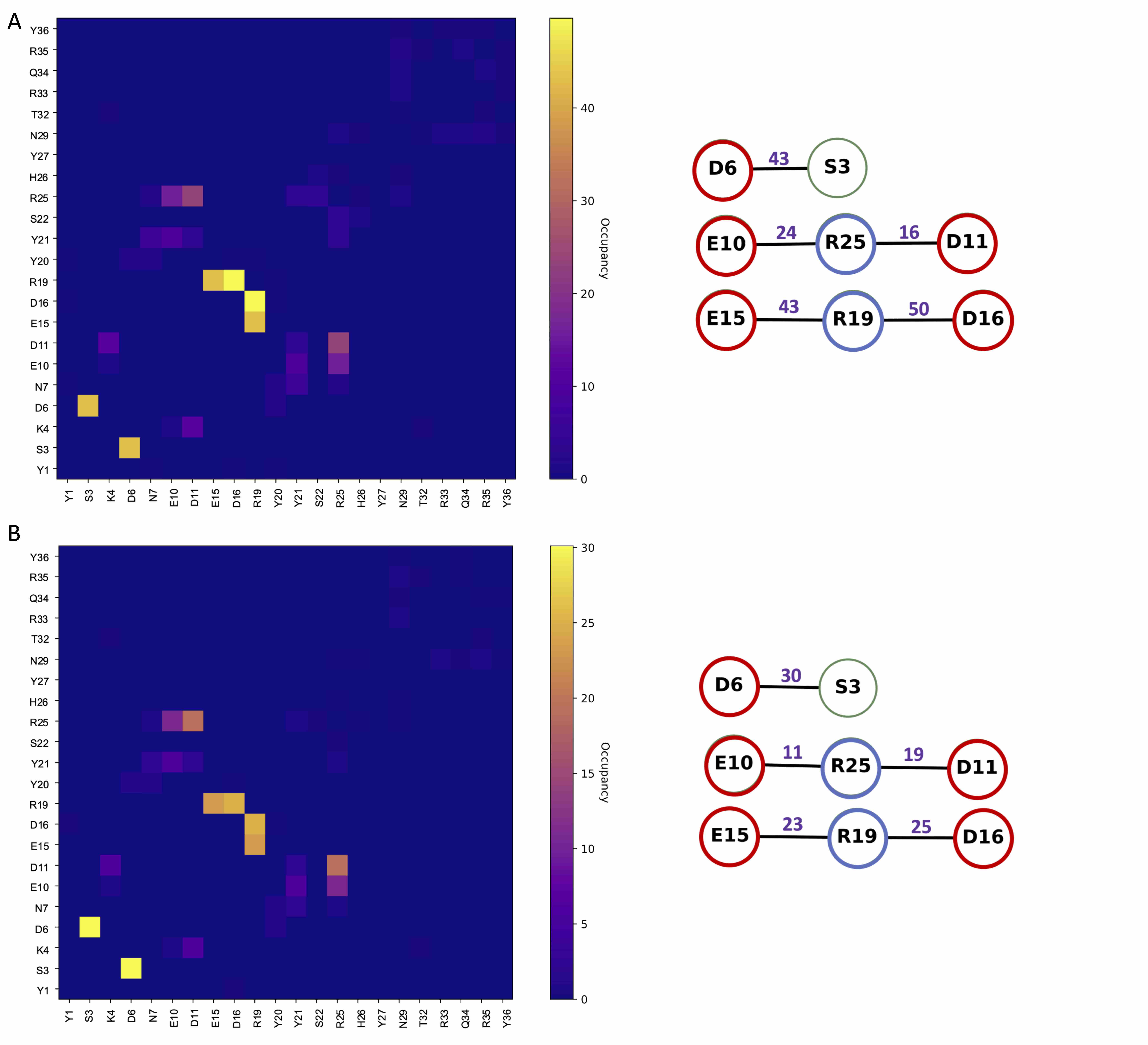

### S58.Fig

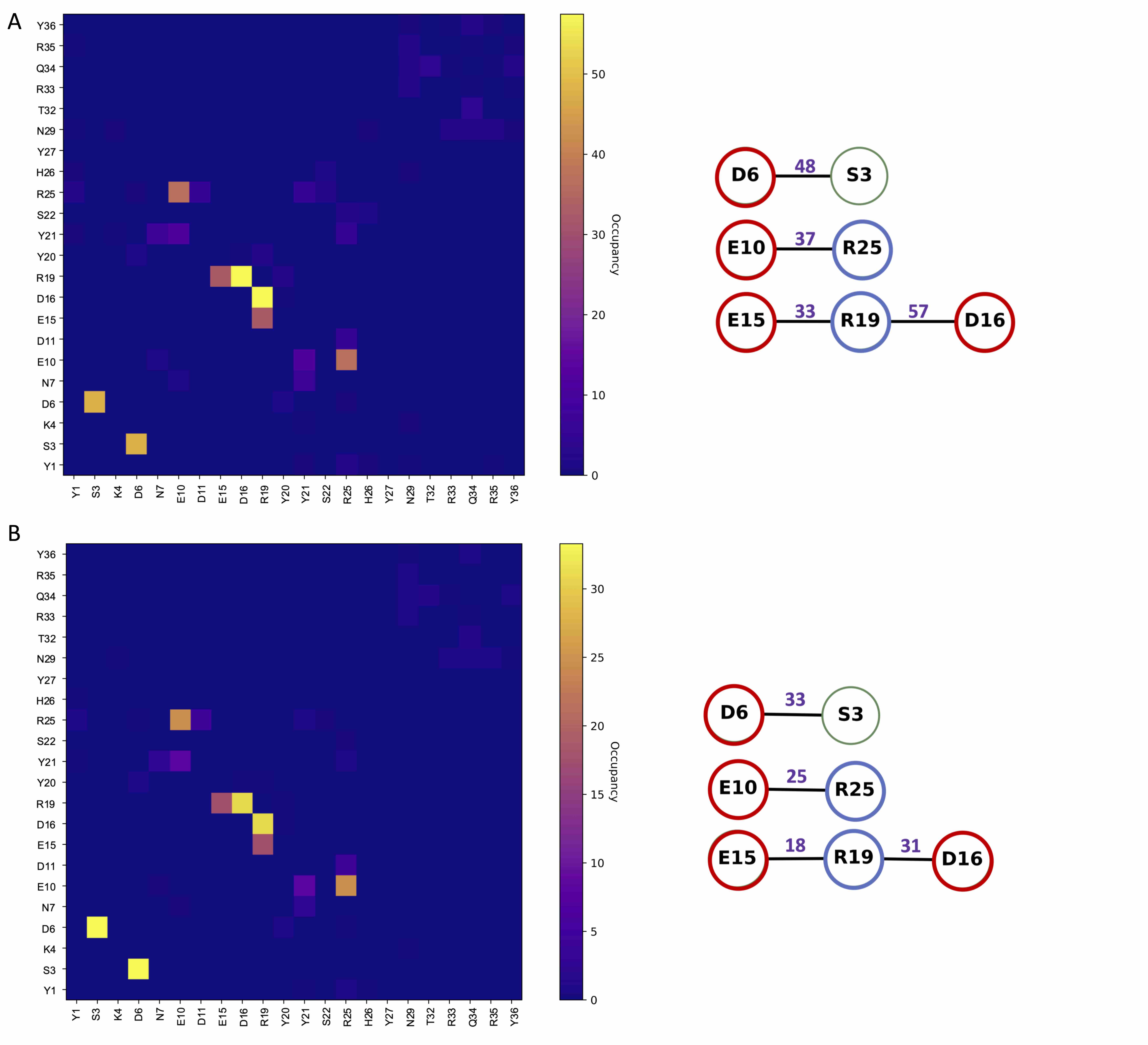

### S59.Fig

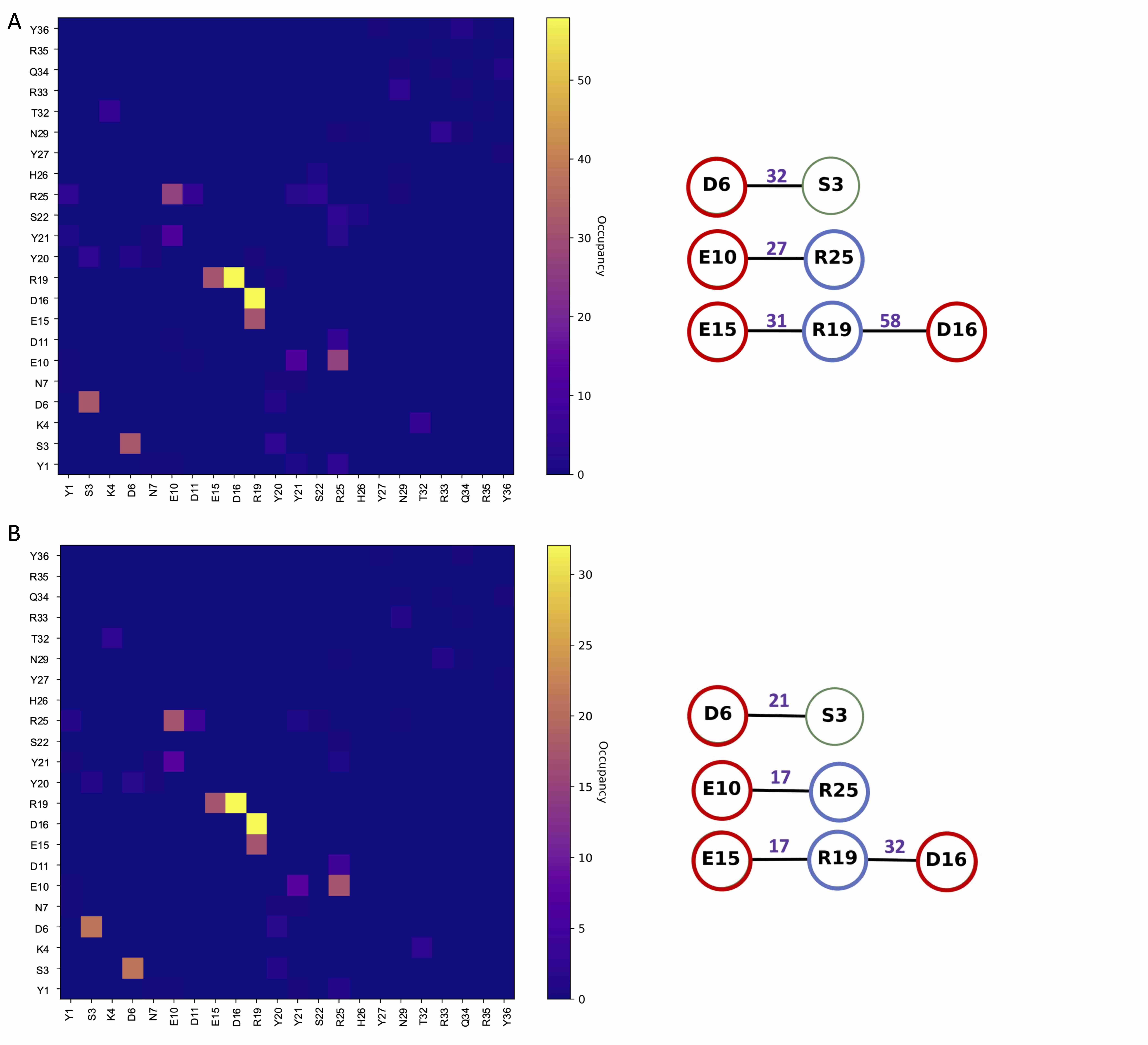

### S60.Fig

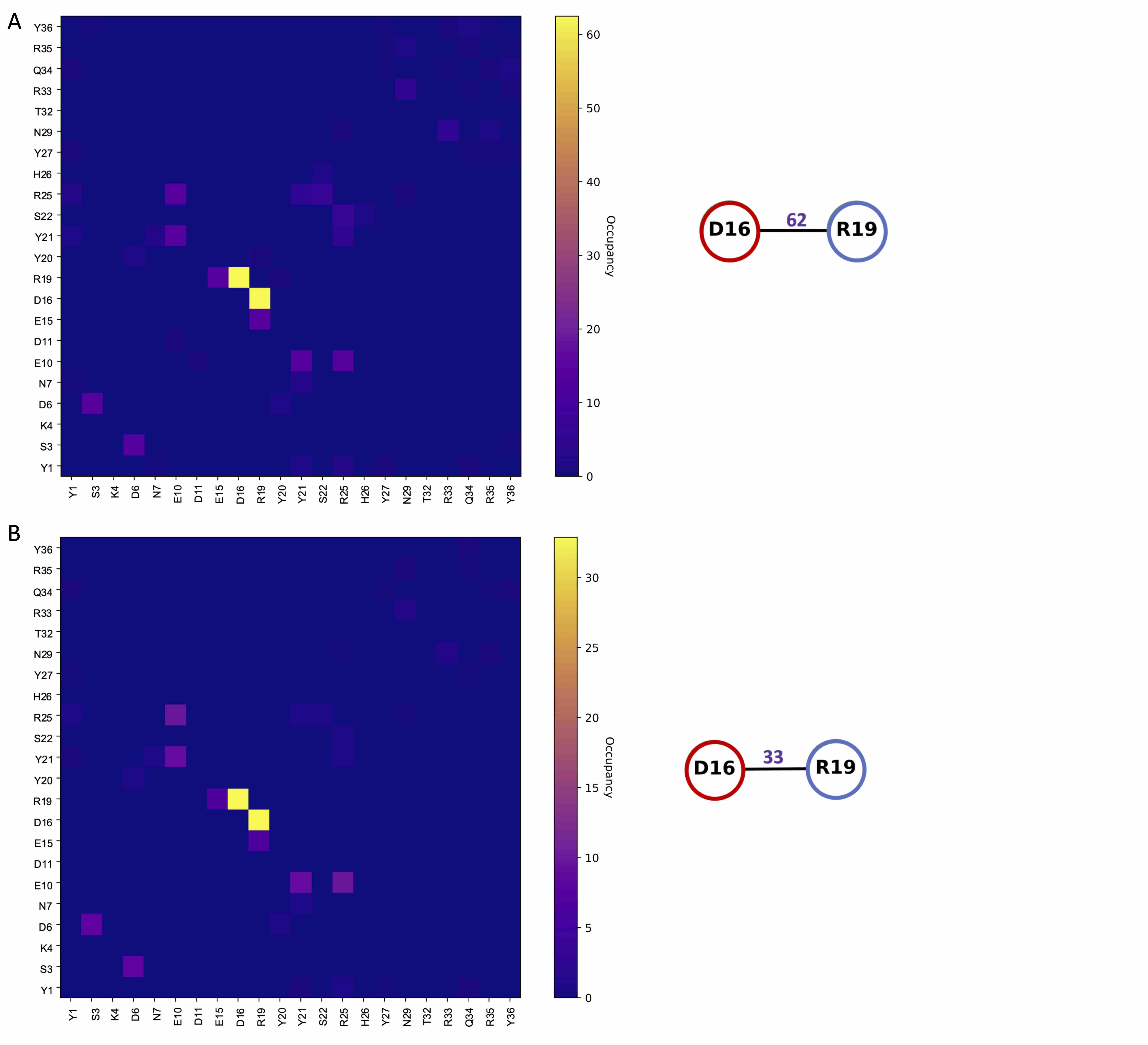

### S61.Fig

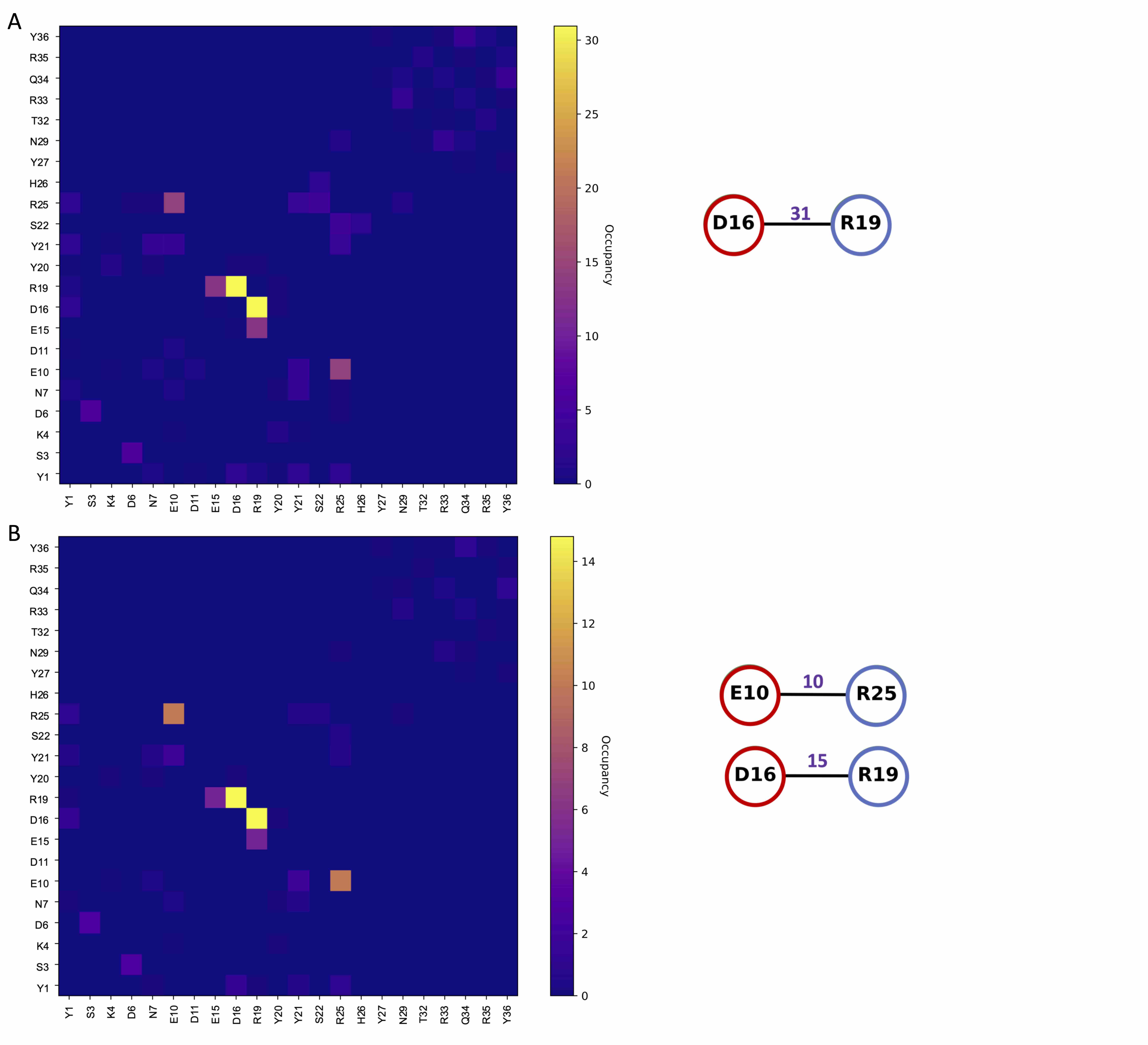

### S62.Fig

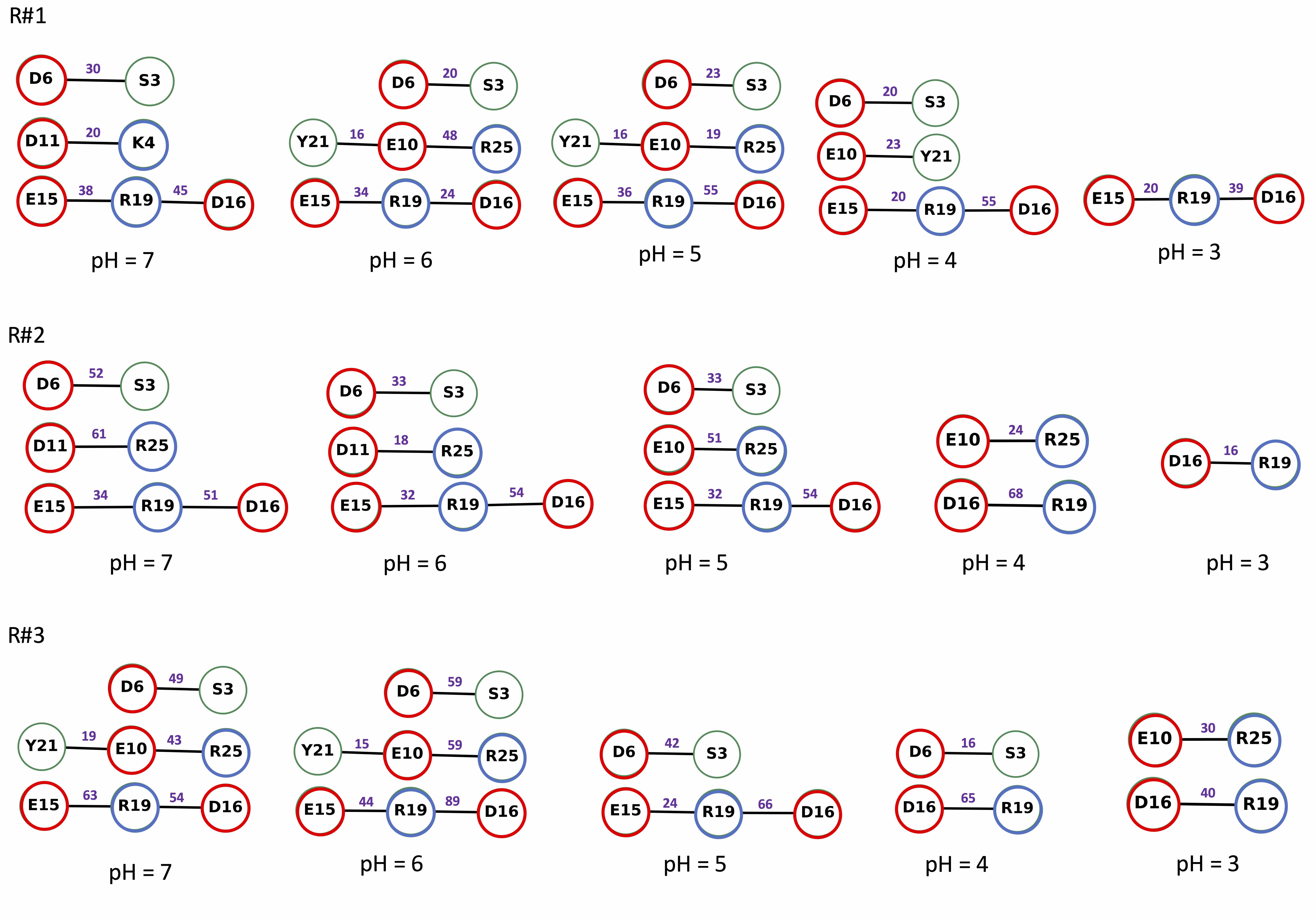

### S63.Fig

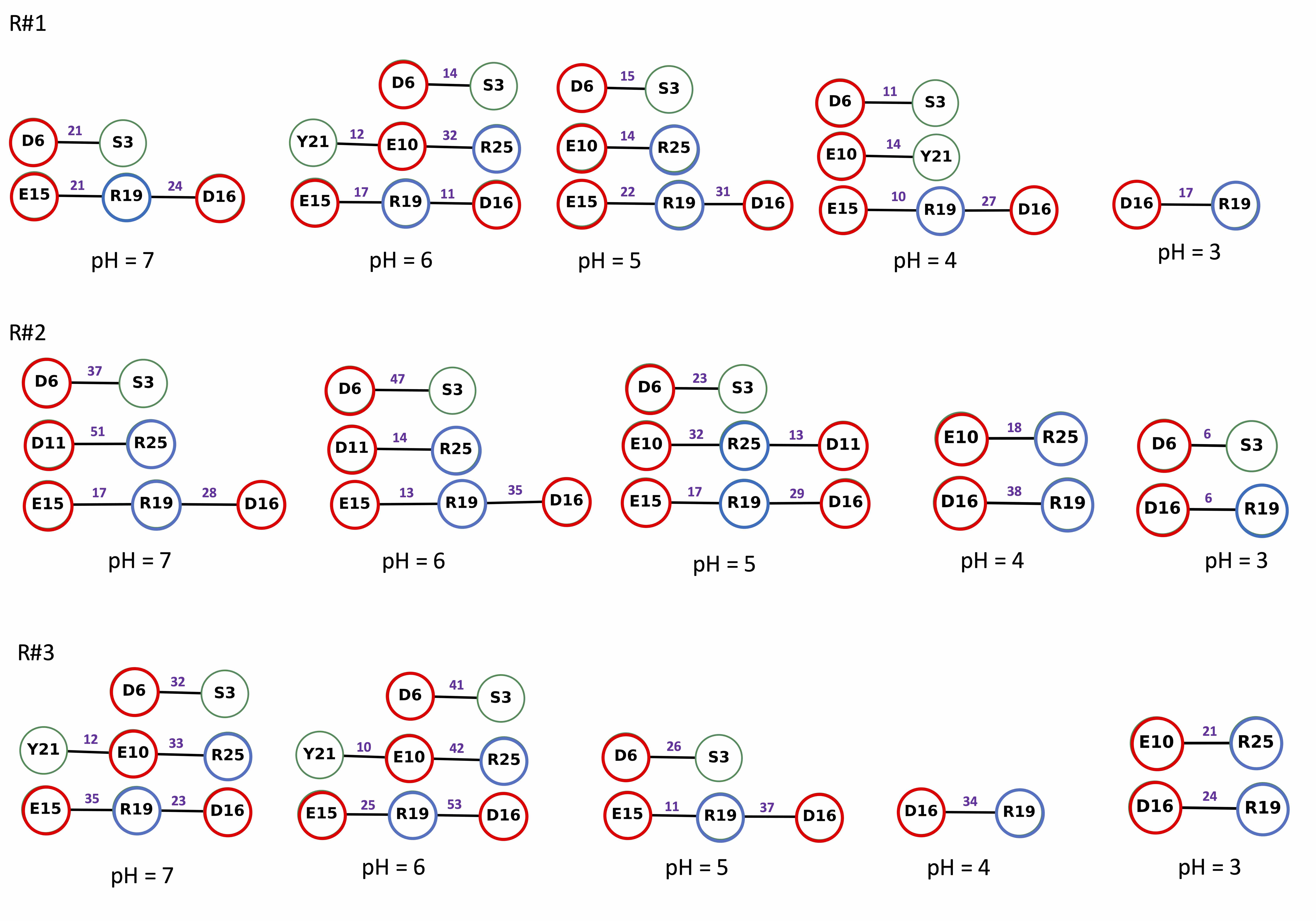

### S65.Fig

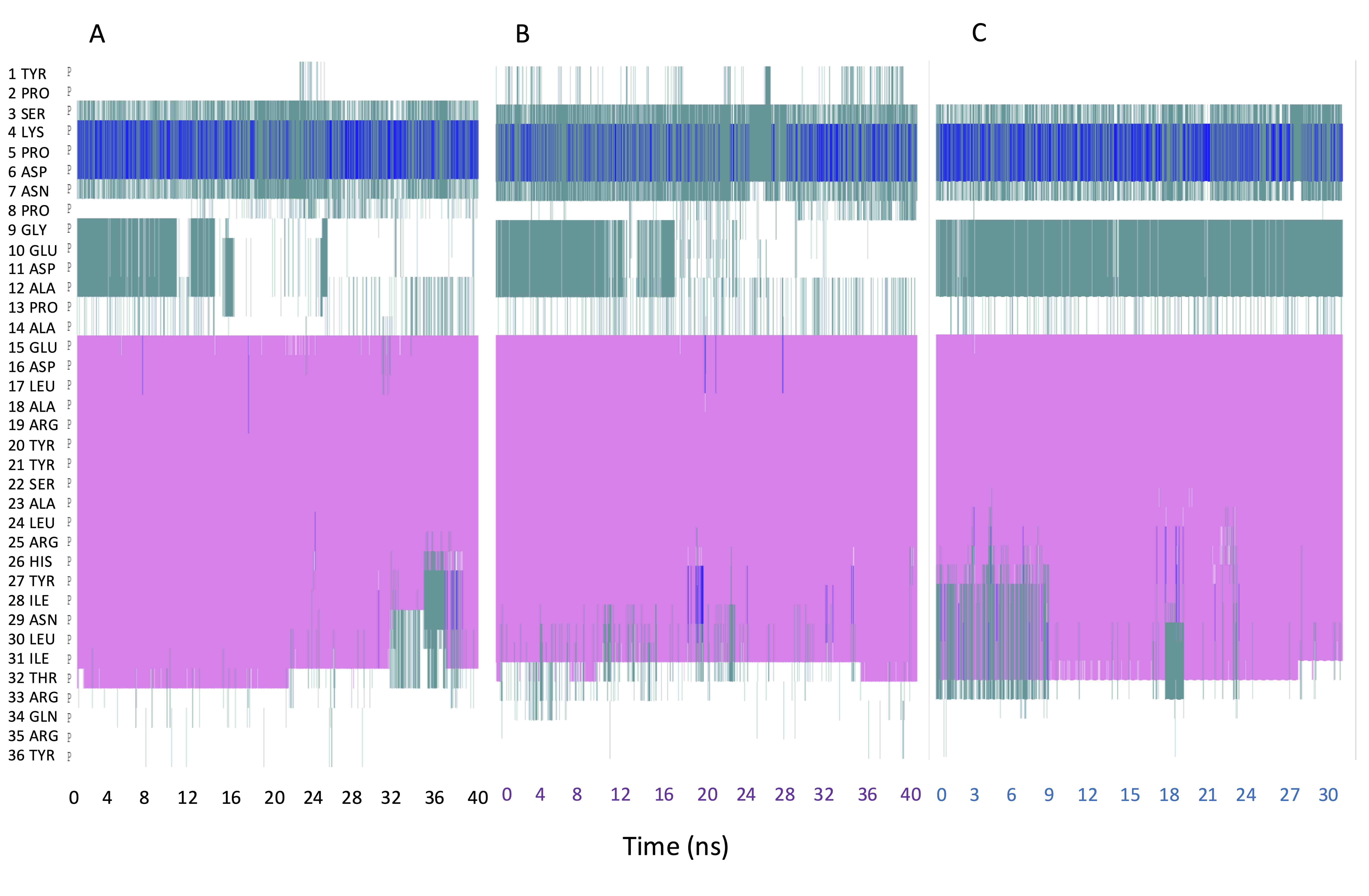

### S66.Fig

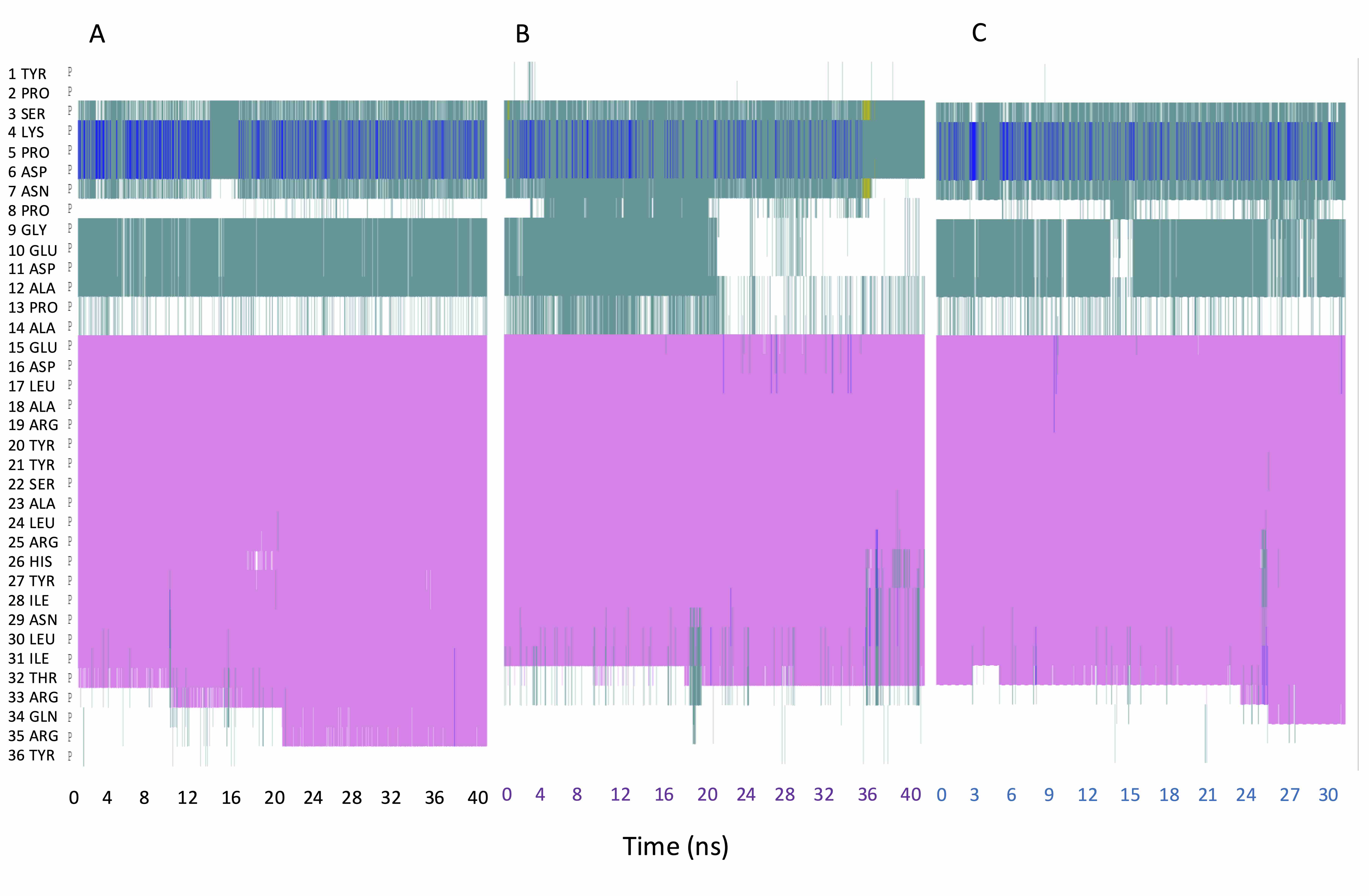

### S67.Fig

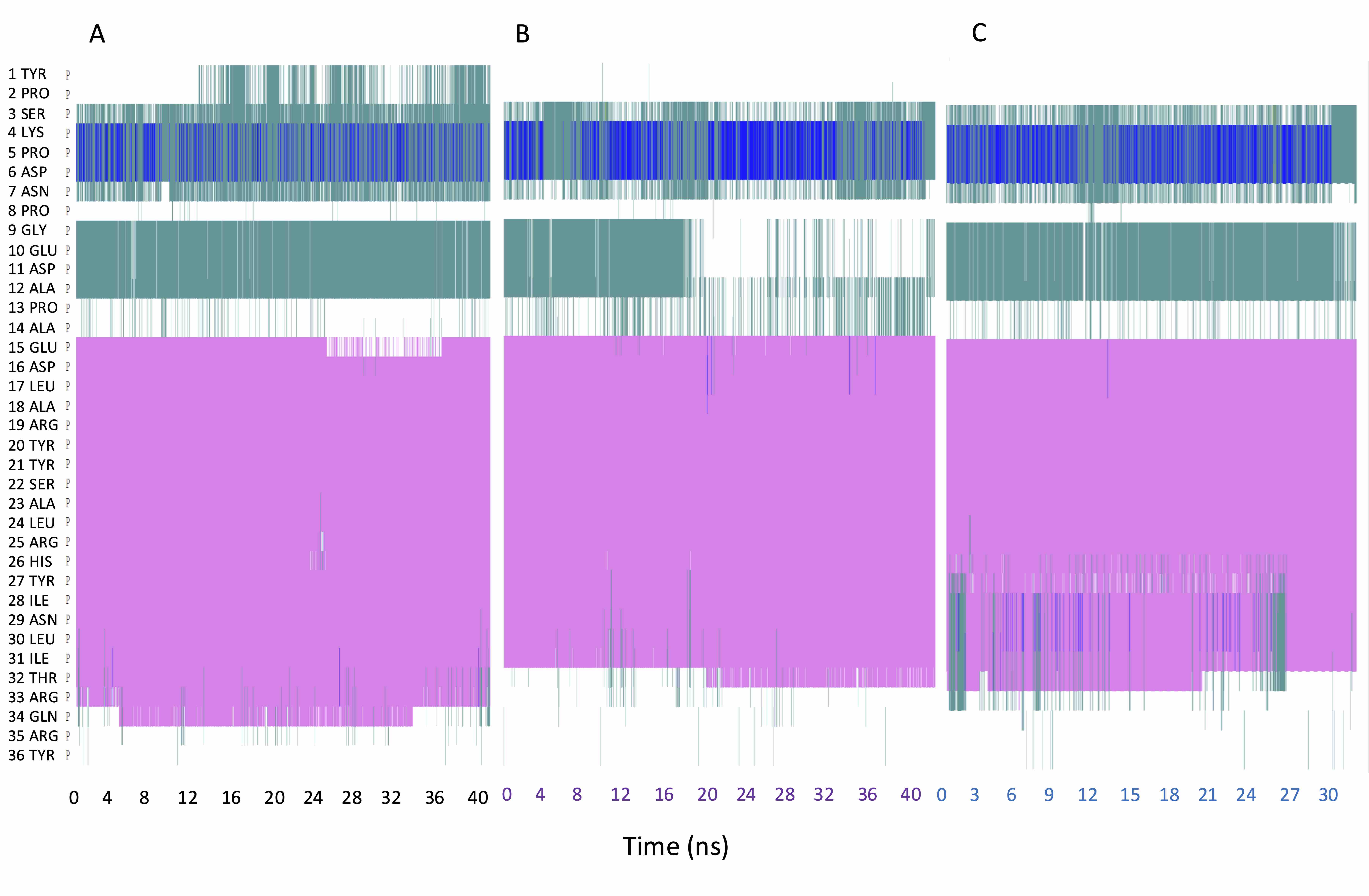

### S68.Fig

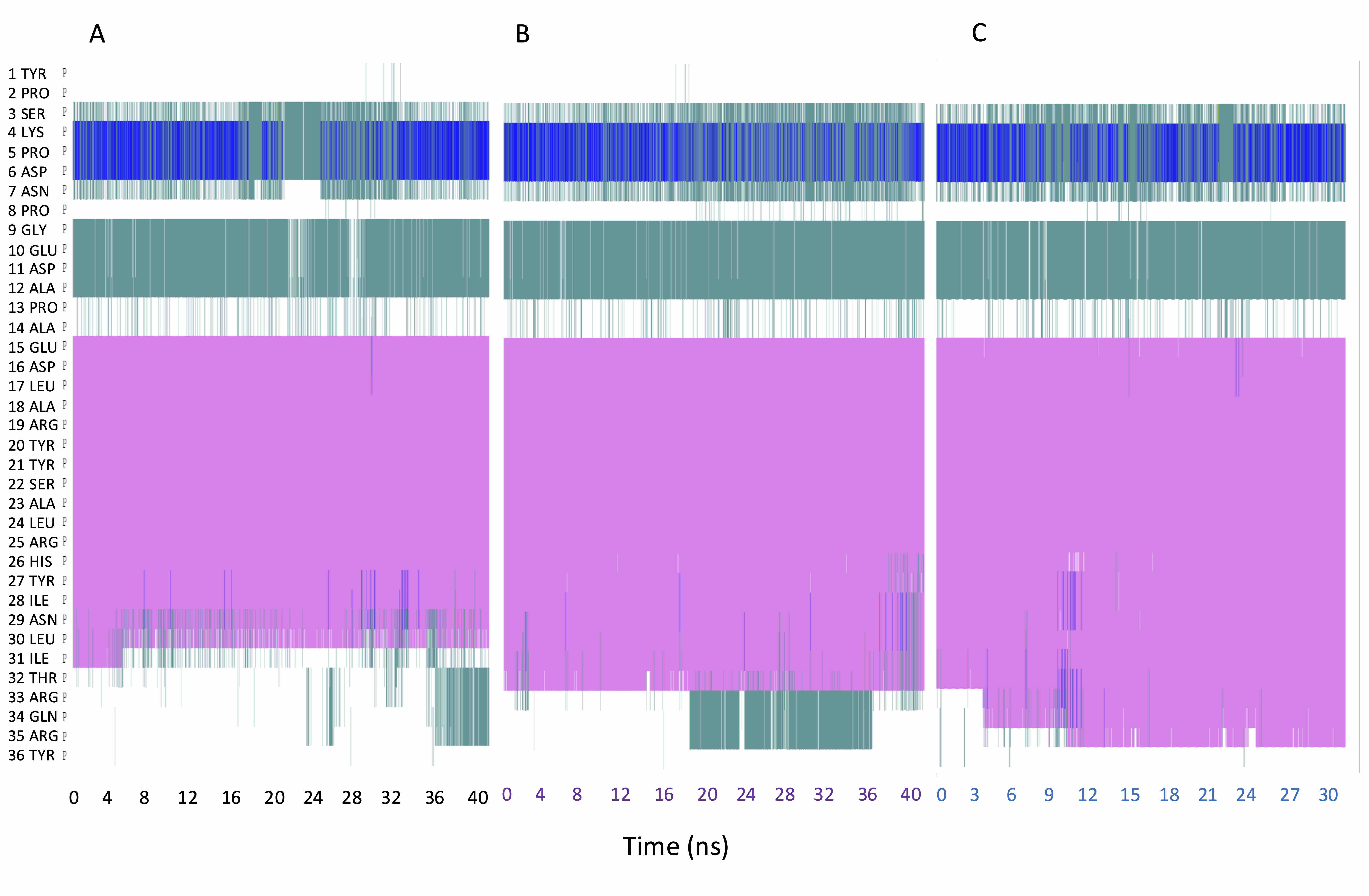

### S69.Fig

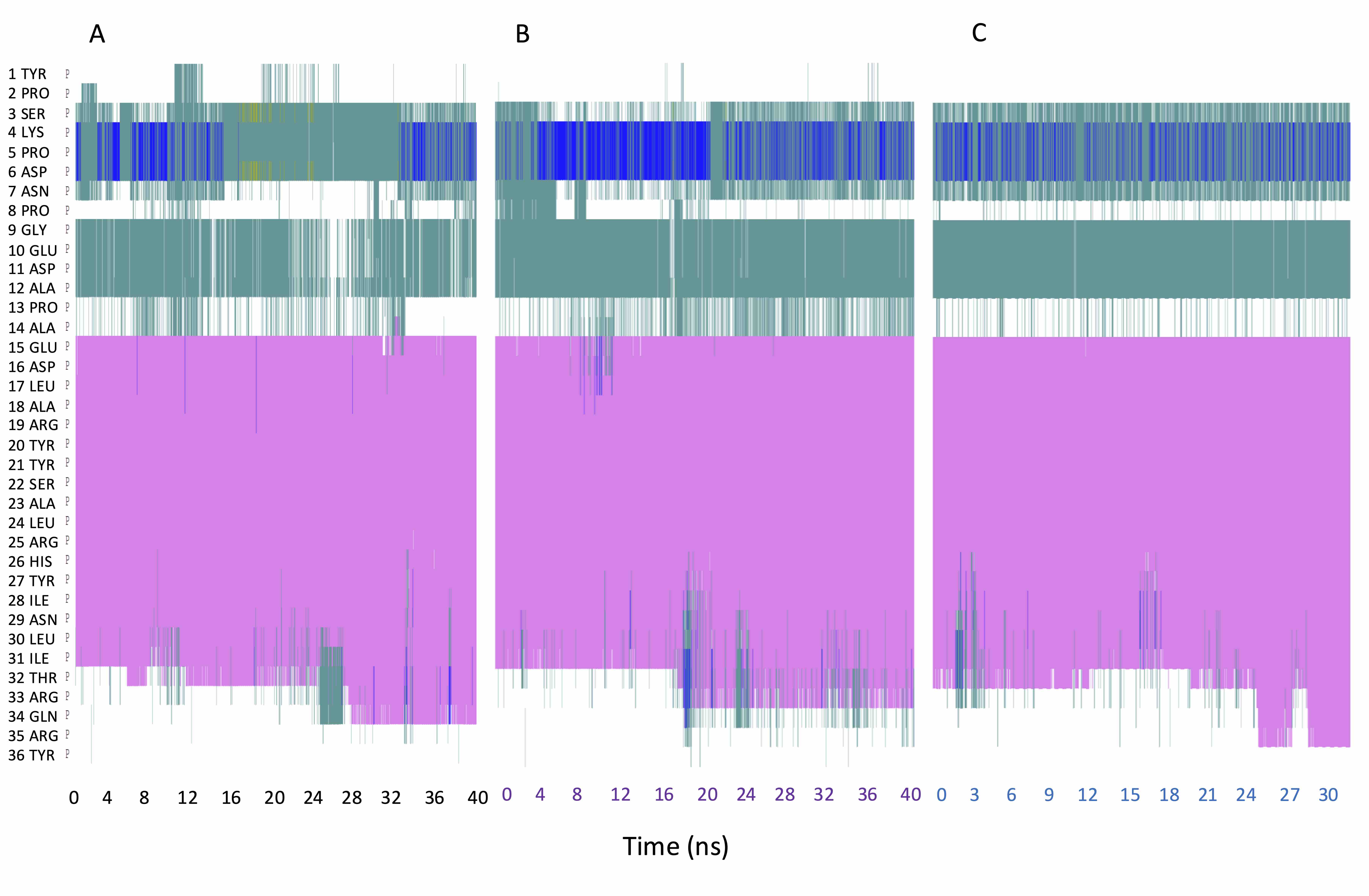

### S70.Fig

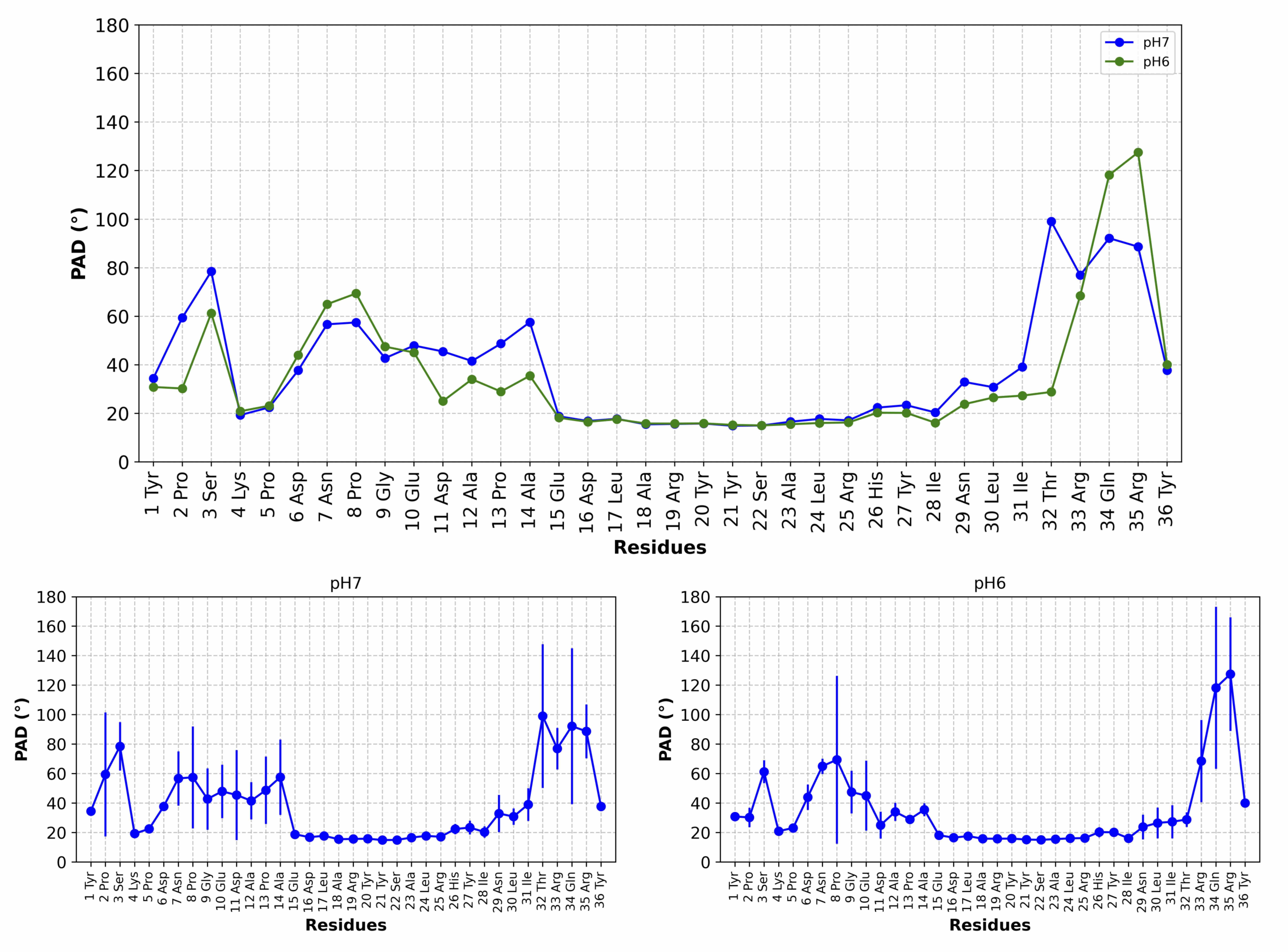

### S71.Fig

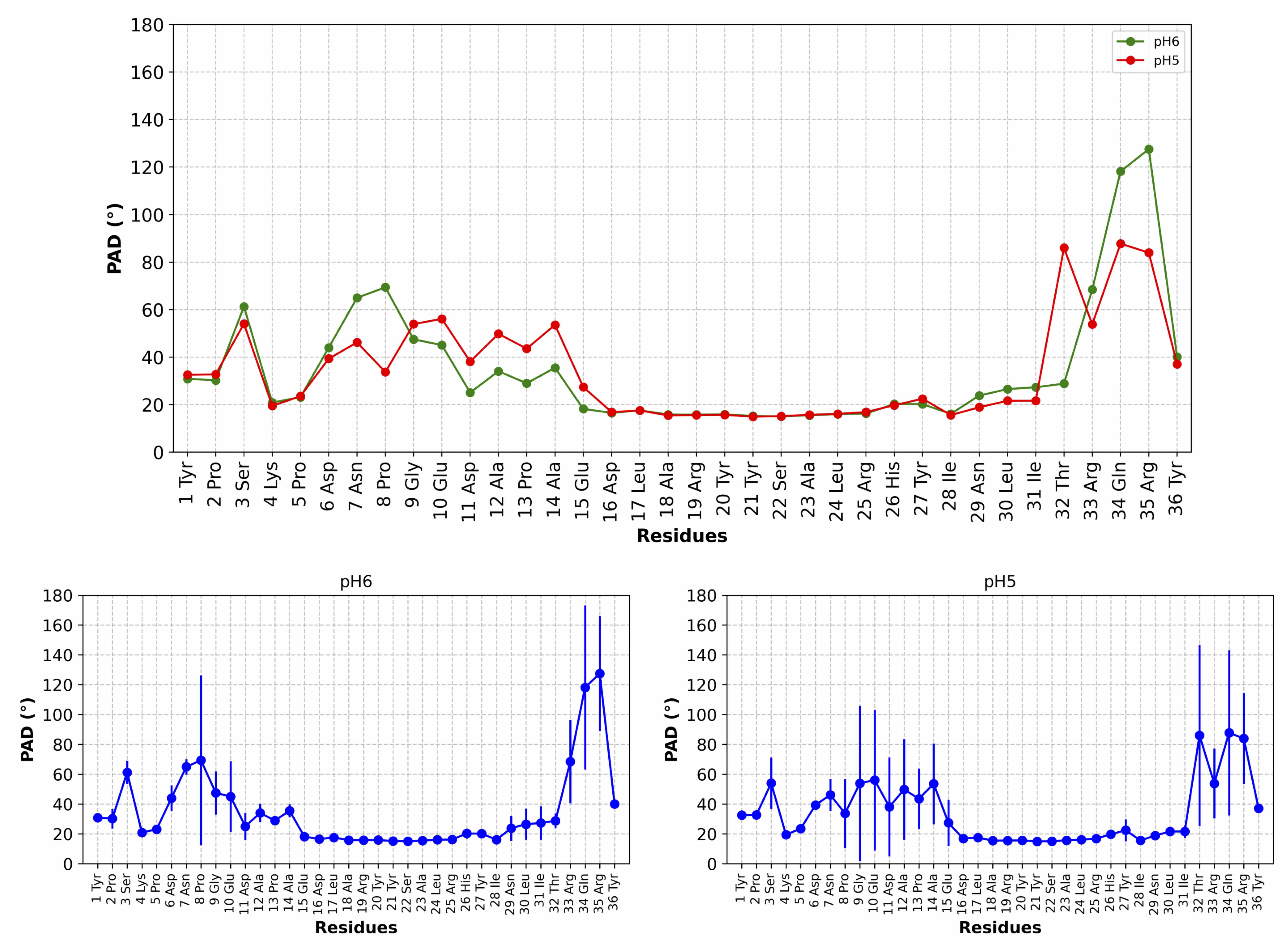

### S72.Fig

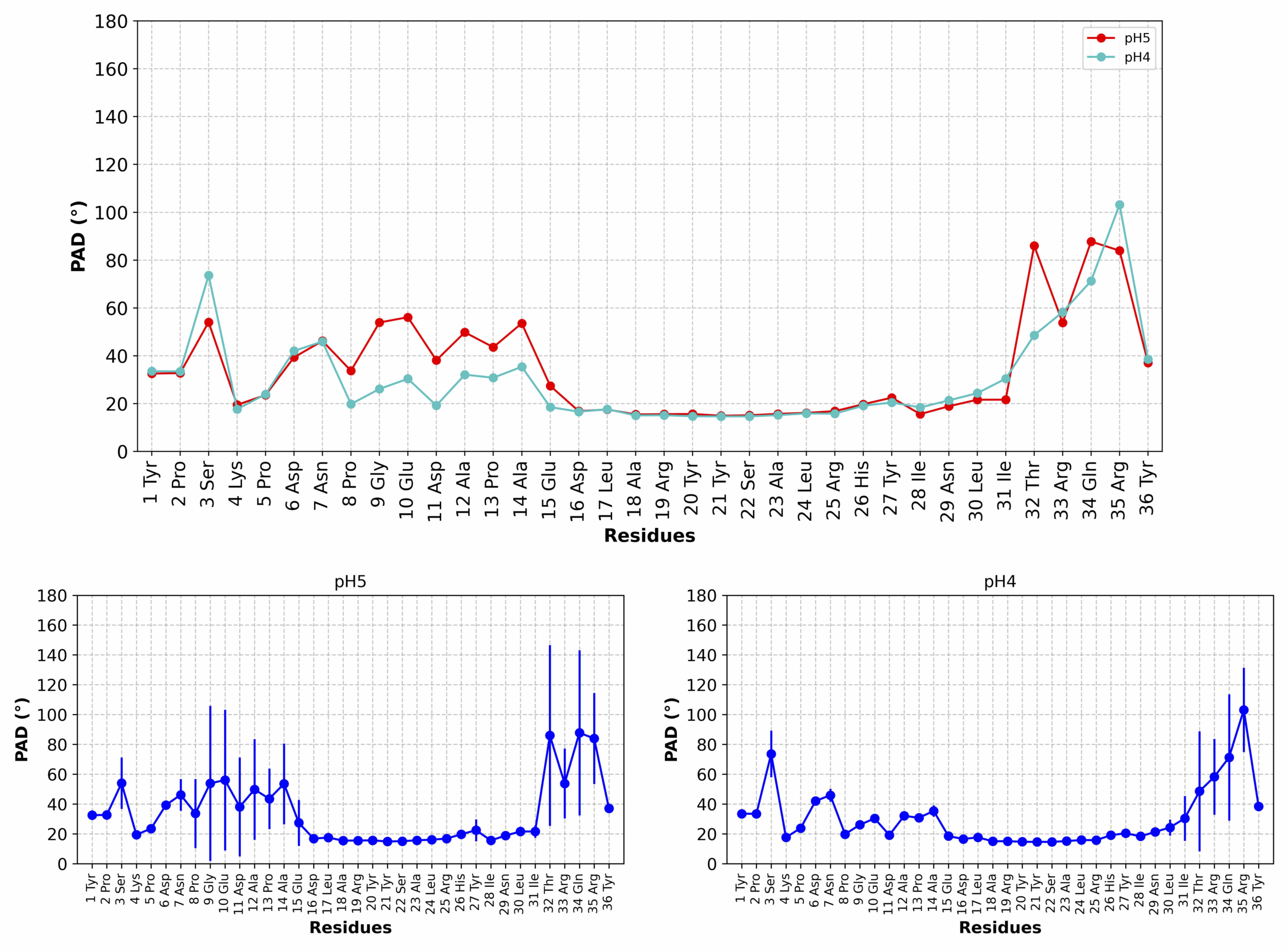

### S73.Fig

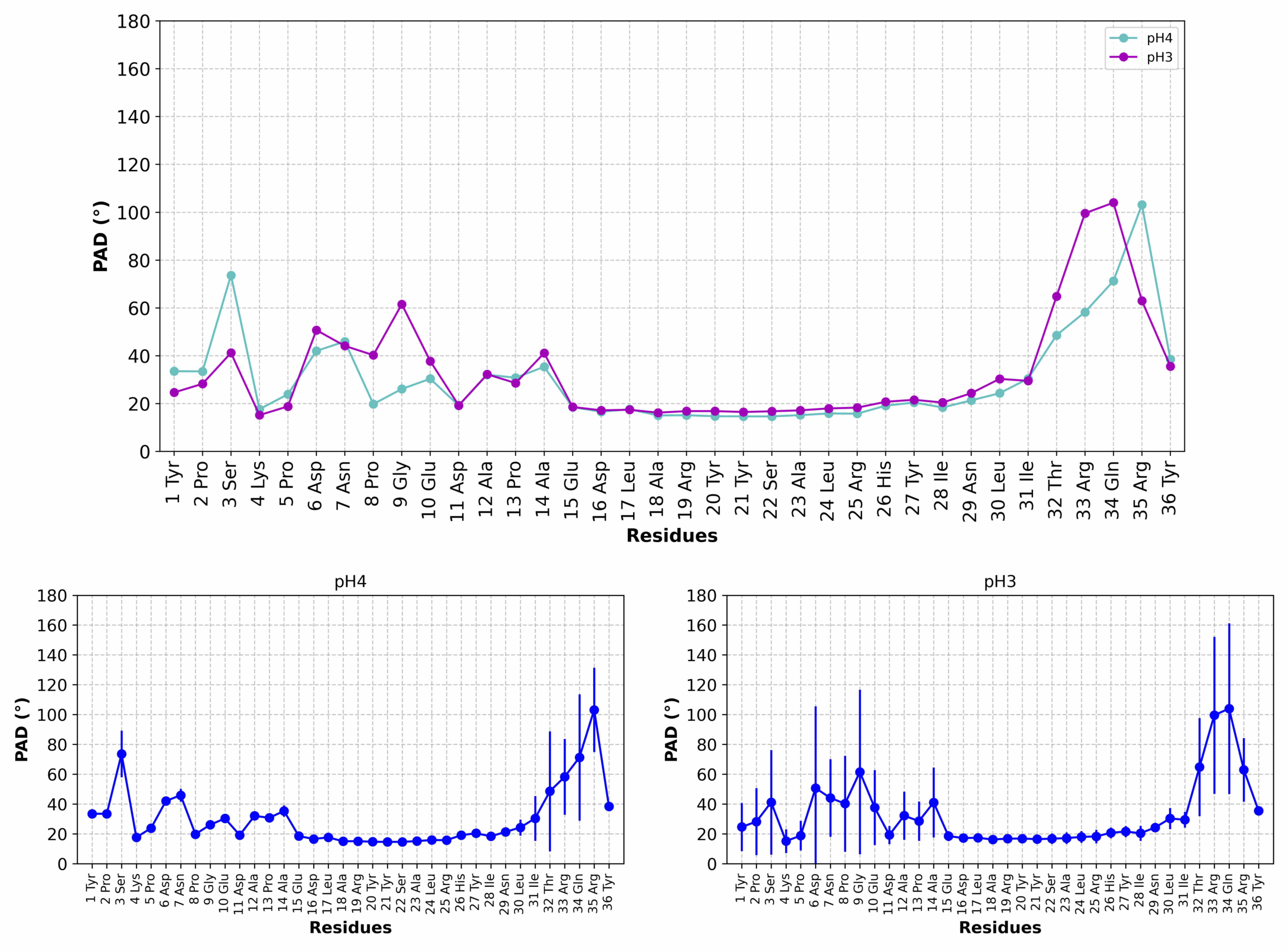
