## Supplementary material for "pH-dependent structural dynamics of the neuropeptide Y in aqueous solution": SI text

*Correspondent authors

### Supporting Information

#### Supporting Information Text

**2. H-bond networks**

We describe here the intramolecular H-bond network using as criterion for the H-bond angle $20^0$ or less (Figures 4-6B (main text); Tables S2-S4). The residues can form direct H-bonds or mediated by 1, 2, 3 water molecules.

**Asp6** side chain H-bonds to **Ser3** side chain across all pH values but pH 3. These H-bonds are direct at pH 7 to 5. **Glu10** side chain H-bonds to (i) **Arg25**. The H-bonds are direct at some pH values; (ii) **Asp11** side chain at pH 7 and 6. These are water-mediated H-bonds. **Asp11** side chain H-bonds also to **Arg25** side chain at pH 7. **Glu15** side chain H-bonds to **Arg19** side chain at all pH values except 3. **Asp16** side chain forms direct H-bonds with **Arg19** side chain at all pH values.

The plot indicates a propensity to form intramolecular hydrogen bonds during deprotonation, a trend already observed using the other criterion for the H-bonds. It also reveals relatively few direct hydrogen bonds between the side chains of the peptide. Thus, as it might be expected, polar and charged side chains tend to interact with water.
