## Supplementary material for "pH-dependent structural dynamics of the neuropeptide Y in aqueous solution": SI table

*Correspondent authors

### Supporting Information

#### Supporting Information Tables

**S1 Table.** Mean $pK_a$ values obtained from our simulations.

| | Model $pK_a$ | R#1 | | R#2 | | R#3 | |
|---|---|---|---|---|---|---|---|
| | | $pK_a$ | $\Delta pK_a$ | $pK_a$ | $\Delta pK_a$ | $pK_a$ | $\Delta pK_a$ |
| Asp6 | 4.0 | 4.5 | 0.5 | 4.5 | 0.5 | 4.5 | 0.5 |
| Glu10 | 4.4 | 3.6 | -0.8 | 3.7 | -0.7 | 3.5 | -0.9 |
| Asp11 | 4.0 | 5.3 | 1.3 | 5.1 | 1.1 | 5.3 | 1.3 |
| Glu15 | 4.4 | 3.5 | -0.9 | 3.7 | -0.7 | 3.7 | -0.7 |
| Asp16 | 4.0 | 2.9 | -1.1 | 3.2 | -0.8 | 2.4 | -1.6 |
| His26 | 7.0 | 6.3 | -0.7 | 6.4 | -0.6 | 6.4 | -0.6 |

**S2 Table.** Occupancies of either direct H-bonds or water-mediated bridges among entire residues (that is, backbone and side chains), averaged over the constant pH simulations at different pH values. This and the next two tables report the occupancies as set in **S39-S49 Figs** (50% and 10% for criteria of $60^0$ and $20^0$, respectively), S50-S56 (25 % and 10%, **Table S3**), and S57-S63 (15% and 10%, **Table S4**). The atoms involved in direct H-bonding are shown. They have the same color code as the amino acids they belong to. As expected, they belong to residues close by.

| pH | H-bond | R#1 | | R#2 | | R#3 | |
|---|---|---|---|---|---|---|---|
| | | $60^0$ | $20^0$ | $60^0$ | $20^0$ | $60^0$ | $20^0$ |
| pH7 | D6-S3 | 96-0.1<br>N-O/OG, OD1/OD2-OG/N | 42-0.3<br>N-O, OD1/OD2-OG/N | 95-0.1<br>N-O/OG, O/OD1/OD2-OG | 45-0.1<br>N-O, OD1/OD2-OG | 96-0.1<br>N-O/OG, OD1/OD2-OG/N | 46-0.2<br>N-O, OD1/OD2-OG |
| | E10-P8 | No | No | No | No | 51-1.4 (only water-mediated) | No |
| | E10-D11 | 63-1.4<br>OE1/OE2-N | 17-1.4<br>OE2-N | 50-1.7<br>OE1/OE2-N | 10-1.6 (only water-mediated) | 83-1.2<br>OE1/OE2-N | 29-1.2<br>OE1/OE2-N |
| | E10-Y21 | No | No | No | No | 70-1.2<br>OE1/OE2-OH | 24-0.7<br>OE1/OE2-OH |
| | E10-R25 | No | No | No | No | 75-0.8<br>OE1/OE2-NH1/NH2 | 40-0.3<br>OE1/OE2-NH1/NH2 |

| | | | | | | | |
|---|---|---|---|---|---|---|---|
| | D11-K4 | No | 10-0.1 OD1/OD2-NZ | No | No | No | No |
| | D11-A12 | 52-1.4 OD1/OD2-N | No | No | No | 66-1.9 (only water-mediated) | 11-2.1 (only water-mediated) |
| | D11-R25 | No | No | 73-0.3 OD1/OD2-NH1/NH2 | 52-0.0 OD1/OD2-NH1/NH2 | No | No |
| | E15-D16 | No | No | No | No | 56-1.9 (only water-mediated) | 12-1.6 (only water-mediated) |
| | E15-R19 | 95-0.1 O-N/NH2, OE1/OE2-NE/NH1/NH2 | 49-0.1 O-N, OE1/OE2-NE/NH1/NH2 | 97-0.0 O-N, OE1/OE2-NE/NH1/NH2 | 50-0.1 O-N, OE1/OE2-NE/NH1/NH2 | 99-0.0 O-N, OE1/OE2-NE/NH1/NH2 | 60-0.1 O-N, OE1/OE2-NH1/NH2 |
| | D16-R19 | 74-0.4 O-N, OD1/OD2-NE/NH1/NH2 | 31-0.3 OD1-NE/NH1/NH2, OD2-NH1/NH2 | 70-0.3 O-N, OD1/OD2-NE/NH1/NH2 | 32-0.2 OD1/OD2-NH1/NH2 | 84-0.4 O-N, OD1-NE/NH1/NH2, OD2-NH1/NH2 | 29-0.3 OD1/OD2-NH1/NH2 |
| | D16-Y20 | 87-0.0 O-N OD2-OH | 21-0.1 O-N | 83-0.0 O-N | 21-0.0 O-N | 91-0.0 O-N | 24-0.0 O-N |
| | H26-S22 | 94-0.1 ND1/N-O | 32-0.1 ND1/N-O | 92-0.1 ND1/N-O, NE2-OG | 27-0.1 ND1/N-O | 84-0.3 ND1/N-O, NE2-OG | 26-0.1 ND1/N-O |
| | H26-N29 | 55-1.2 O-N, ND1-O/ND2 | No | 54-0.7 O-N/ND2 | No | 52-1.3 O-N | No |
| | H26-L30 | 77-0.1 O-N | 17-0.0 O-N | 71-0.0 O-N | 15-0.0 O-N | 51-0.1 O-N | No |
| pH6 | D6 -S3 | 98-0.1 N-O/OG, OD1/OD2-N/OG | 32-0.4 N-O, OD1/OD2-N/OG | 94-0.1 N-O/OG, OD1-OG, OD2-N/OG | 54-0.1 N-O/OG, OD1/OD2-OG | 96-0.1 N-O, OD1/OD2-N/OG | 52-0.2 N-O/OG, OD1/OD2-OG |
| | E10-N7 | No | No | No | 12-0.3 OE1-ND2, N-O/OD1 | No | No |
| | E10-D11 | 79-0.1 OE1/OE2-N | 26-1.3 (only water-mediated) | 53-1.3 OE1/OE2-N | 14-1.4 (only water-mediated) | 84-0.9 OE1/OE2-N | 34-0.8 OE1/OE2-N |
| | E10-Y21 | 63-1.4 OE1/OE2-OH | 19-0.6 OE1/OE2-OH | No | No | 64-1.2 OE1/OE2-OH | 20-0.6 OE1/OE2-OH |
| | E10-R25 | 74-0.7 OE1/OE2-NH1/NH2 | 36-0.2 OE1/OE2-NH1/NH2 | No | No | 80-0.5 OE1/OE2-NE/NH1/NH2 | 45-0.1 OE1/OE2-NH1/NH2, OE2-NE |

| | | | | | | | |
|---|---|---|---|---|---|---|---|
| | D11-A12 | 61-1.9 (only water-mediated) | No | No | No | 54-1.9 (only water-mediated) | No |
| | D11-R25 | No | No | No | 17-0.2 OD1/OD2-NH1/NH2 | No | No |
| | E15-D16 | 54-2.0 (only water-mediated) | No | No | No | 58-2.0 (only water-mediated) | 12-1.9 (only water-mediated) |
| | E15-R19 | 99-0.0 O-N, OE1/OE2-NE/HN1/NH2 | 51-0.1 O-N, OE1/OE2-NE/HN1/NH2 | 94-0.1 O-N, OE1/OE2-NE/HN1/NH2 | 39-0.2 O-N, OE1/OE2-NE/HN1/NH2 | 98-0.0 O-N, OE1/OE2-HN1/NH2 | 49-0.1 O-N, OE1/OE2-HN1/NH2 |
| | D16-R19 | 64-0.0 O-N, OD1-NE/NH1/NH2, OD2-NH1/NH2 | 15-0.5 OD1/OD2-NH1/NH2 | 85-0.2 O-N, OD1/OD2-NE/NH1/NH2 | 41-0.2 OD1/OD2-NH1/NH2 | 97-0.1 O-N/NE, OD1/OD2-NE/NH1/NH2 | 58-0.1 OD1/OD2-NE/NH1/NH2 |
| | D16-Y20 | 89-0.0 O-N | 24-0.0 O-N | 84-0.1 O-N | 32-0.1 O-N | 94-0.0 O-N, OD1-OH | 38-0.1 O-N |
| | H26-S22 | 93-0.1 N/ND1-O, NE2-OG | 34-0.1 N/ND1-O | 94-0.0 N/ND1-O, NE2-OG | 34-0.0 N/ND1-O | 92-0.1 N/ND1-O, NE2-OG | 31-0.1 N-O |
| | H26-N29 | 53-1.0 O-N, ND1-OD1 | No | No | No | No | No |
| | H26-L30 | 82-0.0 O-N | 18-0.0 O-N | 69-0.0 O-N | 21-0.0 O-N | 85-0.0 O-N | 20-0.0 O-N |
| pH5 | D6-S3 | 95-0.1 N-O/OG, OD1/OD2-OG | 29-0.2 N-O, OD1/OD2-OG | 94-0.1 N-O/OG, OD1/OD2-N/OG | 48-0.1 N-O/OG, OD1/OD2-N/OG | 93-0.1 N-O/OG, OD1/OD2-OG | 37-0.2 N-O, OD1/OD2-OG |
| | E10-P8 | 52-1.4 (only water-mediated) | No | No | No | No | No |
| | E10-D11 | 71-1.5 OE1/OE2-N | 17-1.3 OE1/OE2-N | 54-1.2 OE1/OE2-N | 18-1.2 OE2-N | 59-1.6 OE2-N | 13-1.5 (only water-mediated) |
| | E10-Y21 | 67-1.3 OE1/OE2-OH | 20-0.8 OE1/OE2-OH | No | No | 58-1.5 OE1/OE2-OH | 16-1.0 OE1/OE2-OH |
| | E10-R25 | No | 17-0.3 OE1/OE2-NH1/NH2 | 67-0.4 O-NH1, OE1/OE2-NE/NH1/NH2 | 34-0.1 OE1/OE2-NE/NH1/NH2 | No | No |

| | | | | | | | |
|---|---|---|---|---|---|---|---|
| | D11-A12 | 55-1.9 (only water-mediated) | No | No | No | No | No |
| | E15-R19 | 99-0.0 O-N, OE1/OE2-NE/NH1/NH2 | 53-0.1 O-N, OE1/OE2-NE/NH1/NH2 | 95-0.1 O-N, OE1/OE2-NE/NH1/NH2 | 43-0.1 O-N, OE1/OE2-NE/NH1/NH2 | 98-0.0 O-N, OE1/OE2-NE/NH1/NH2 | 46-0.1 O-N, OE1/OE2-NH1/NH2 |
| | D16-R19 | 82-0.4 O-N, OD1/OD2-NE/NH1/NH2 | 38-0.2 OD1-NH1/NH2, OD2-NE/NH1/NH2 | 76-0.3 O-N, OD1/OD2-NE/NH1/NH2 | 32-0.1 OD1/OD2-NE/NH1/NH2 | 83-0.3 O-N, OD1/OD2-NE/NH1/NH2 | 41-0.1 OD1-NH1/NH2, OD2-NE/NH1/NH2 |
| | D16-Y20 | 95-0.0 O-N | 24-0.0 O-N | 84-0.1 O-N | 28-0.1 O-N | 90-0.0 O-N | 22-0.0 O-N |
| | H26-S22 | 89-0.1 N/ND1-O, NE2-OG | 30-0 N/ND1-O | 94-0.0 N/ND1-O, NE2-OG | 34-0.0 N-O | 91-0.0 N/ND1-O, NE2-OG | 17-0.1 N/ND1-O |
| | H26-N29 | 53-0.8 O-N/ND2 | No | No | No | No | No |
| | H26-L30 | 85-0.0 O-N | 22-0.0 O-N | 83-0.0 O-N | 17-0.0 N-O | No | No |
| pH4 | D6-S3 | 92-0.1 N-O/OG, OD1/OD2-N/OG | 27-0.2 N-O, OD1/OD2-N/OG | 96-0.1 N-O, OD1/OD2-OG | 19-0.2 N-O, OD1/OD2-OG | 95-0.1 N-O, OD1/OD2-N/OG | 25-0.3 N-O, OD1-OG, OD2-N/OG |
| | E10-D11 | 59-1.5 OE1/OE2-N | 12-1.4 OE1-N | 55-1.5 OE1/OE2-N | 12-1.3 OE1-N | 58-1.7 (only water-mediated) | No |
| | E10-Y21 | 60-1.1 OE1/OE2-OH | 20-0.5 OE1/OE2-OH | 51-1.5 OE1/OE2-OH | 13-0.7 OE1/OE2-OH | 59-1.5 OE1/OE2-OH | 15-0.9 OE1/OE2-OH |
| | E10-R25 | No | 11-0.7 OE1/OE2-NH1/NH2 | No | 21-0.2 OE1/OE2-NH1/NH2 | No | No |
| | E15-R19 | 97-0.0 O-N, OE1/OE2-NE/NH1/NH2 | 49-0.1 O-N, OE1/OE2-NE/NH1/NH2 | 98-0.0 O-N, OE1/OE2-NH1/NH2 | 42-0.1 OE1/OE2-NH1/NH2 | 95-0.0 O-N/NE, OE1/OE2-NH1/NH2 | 34-0.1 O-N, OE1/OE2-NH1/NH2 |
| | D16-R19 | 81-0.4 O-N, OD1/OD2-NE/NH1/NH2 | 32-0.2 OD1/OD2-NE/NH1/NH2 | 85-0.2 O-N, OD1/OD2-NE/NH1/NH2 | 43-0.1 OD1/OD2-NE/NH1/NH2 | 83-0.3 O-N, OD1/OD2-NE/NH1/NH2 | 37-0.1 O-N, OD1/OD2-NE/NH1/NH2 |
| | D16-Y20 | 89-0.0 O-N | 23-0.0 O-N | 90-0.0 O-N | 25-0.0 O-N | 89-0.0 O-N | 29-0.0 O-N |
| | H26-S22 | 92-0.0 N/ND1-O, NE2-OG | 30-0.1 N/ND1-O | 87-0.1 N/ND1-O, NE2-OG | 24-0.1 N/ND1-O | 97-0.0 N/ND1-O, NE2-OG | 35-0.1 N/ND1-O |
| | H26-N29 | 63-0.4 O-N/ND2 | No | No | No | 53-0.7 O-N/ND2 | No |

| | | | | | | | |
|---|---|---|---|---|---|---|---|
| | H26-L30 | 58-0.0 O-N | No | 71-0.0 O-N | 14-0.0 O-N | 79-0.0 O-N | 18-0.1 O-N |
| pH3 | D6-S3 | 90-0.2 N-O, OD1/OD2-N/OG | 20-0.3 N-O, OD1/OD2-OG | 97-0.1 N-O, OD1/OD2-N/OG | 26-0.2 N-O, OD1/OD2-N/OG | 99-0.0 N-O | 13-0.4 N-O |
| | E10-G9 | No | No | 53-1.4 OE1-N | No | No | No |
| | E10-D11 | 52-1.7 OE1/OE2-N | No | 57-1.9 OE1/OE2-N | No | 57-1.5 OE1/OE2-N | 11-1.3 OE1-N |
| | E10-R25 | No | No | No | No | 53-0.8 OE1/OE2-NH1/NH2 | 24-0.0 OE1/OE2-NH1/NH2 |
| | E15-D16 | No | No | 52-1.9 (only water-mediated) | No | No | No |
| | E15-R19 | 97-0.1 O-N, OE1/OE2-NE/NH1/NH2 | 47-0.1 O-N, OE1/OE2-NH1/NH2 | 90-0.1 (only water-mediated) | 38-0.1 O-N, OE1-NE/NH1 OE2-NH1/NH2 | 98-0.0 O-N, OE1/OE2-NH1/NH2 | 42-0.0 O-N, OE2-NH1 |
| | D16-R19 | 73-0.6 O-N, OD1/OD2-NE/NH1/NH2 | 22-0.3 O-N, OD1/OD2-NH1/NH2 | 63-0.9 O-N, OD1/OD2-NE/NH1/NH2 | 11-0.6 OD1/OD2-NH1/NH2 | 76-0.7 O-N, OD1/OD2-NE/NH1/NH2 | 29-0.0 OD1/OD2-NE/NH1/NH2 |
| | D16-Y20 | 88-0.0 O-N | 20-0.0 O-N | 85-0.1 O-N, OD1/OD2-OH | 26-0.0 O-N | 93-0.0 O-N | 26-0.0 O-N |
| | H26-S22 | 91-0.0 N-O | 27-0.1 N-O | 95-0.1 N/ND1-O, NE2-OG | 31-0.1 N/ND1-O | 95-0.1 N/ND1-O, NE2-OG | 27-0.0 N/ND1-O |
| | H26-N29 | No | No | 51-0.9 O-N | No | No | No |
| | H26-L30 | 80-0.0 O-N | 19-0.0 O-N | 69-0.0 O-N | 10-0.0 O-N | 72-0.0 O-N | 17-0.0 O-N |

**S3 Table.** Same as **S2** but only for side chains and the occupancy of 25% for the criterion of $60^0$ and 10% for the criterion of $20^0$.

| pH | H-bond criterion | R#1 | | R#2 | | R#3 | |
|---|---|---|---|---|---|---|---|
| | | $60^0$ | $20^0$ | $60^0$ | $20^0$ | $60^0$ | $20^0$ |
| pH7 | D6-S3 | 58-1.0 OD1/OD2-OG | 26-0.3 OD1/OD2-OG | 67-0.4 OD1/OD2-OG | 40-0.1 OD1/OD2-OG | 74-0.6 OD1/OD2-OG | 38-0.2 OD1/OD2-OG |
| | D6-Y20 | No | No | No | No | 35-1.9 | No |

| | | | | | | | |
|---|---|---|---|---|---|---|---|
| | | | | | | (only water-mediated) | |
| | E10-D11 | 35-1.9 (only water-mediated) | No | 29-2.0 (only water-mediated) | No | 61-1.6 (only water-mediated) | 18-1.4 (only water-mediated) |
| | E10-Y21 | No | No | No | No | 60-1.1 OE1/OE2-OH | 22-0.6 OE1/OE2-OH |
| | E10-R25 | No | No | 26-1.0 (only water-mediated) | No | 75-0.8 OE1/OE2-NH1/NH2 | 40-0.3 OE1/OE2-NH1/NH2 |
| | D11-K4 | No | 10-0.1 (only water-mediated) | No | No | No | No |
| | D11-R25 | No | No | 70-0.3 OD1/OD2-NH1/NH2 | 52-0.0 OD1/OD2-NH1/NH2 | No | No |
| | E15-D16 | 31-2.0 (only water-mediated) | No | No | No | 47-1.9 (only water-mediated) | 10-1.5 (only water-mediated) |
| | E15-R19 | 64-0.6 OE1/OE2-NE/NH1/NH2 | 26-0.3 OE1/OE2-NE/NH1/NH2 | 58-0.7 OE1/OE2-NE/NH1/NH2 | 22-0.3 OE1/OE2-NE/NH1/NH2 | 83-0.4 OE1/OE2-NE/NH1/NH2 | 40-0.2 OE1/OE2-NH1/NH2 |
| | D16-R19 | 67-0.5 OD1/OD2-NE/NH1/NH2 | 30-0.3 OD1-NE/NH1/NH2, OD2-NH1/NH2 | 65-0.9 OD1/OD2-NE/NH1/NH2 | 32-0.2 OD1/OD2-NH1/NH2 | 79-0.4 OD1/OD2-NE/NH1/NH2 | 29-0.2 OD1/OD2-NH1/NH2 |
| | H26-S22 | No | No | No | No | 31-2.0 (only water-mediated) | No |
| | H26-N29 | 32-1.7 (only water-mediated) | No | 25-1.7 NE2-ND2 | No | No | No |
| pH6 | D6-S3 | 48-1.2 OD1/OD2-OG | 17-0.3 OD1/OD2-OG | 76-0.2 OD1/OD2-OG | 49-0.1 OD1/OD2-OG | 74-0.4 OD1/OD2-OG | 44-0.1 OD1/OD2-OG |
| | D6-Y20 | 33-2.0 (only water-mediated | No | No | No | No | No |

| | | | | | | | |
|---|---|---|---|---|---|---|---|
| | E10-D11 | 61-1.7 (only water-mediated) | 17-1.4 (only water-mediated) | 56-1.7 (only water-mediated) | 15-1.4 (only water-mediated) | 29-1.9 (only water-mediated) | No |
| | E10-Y21 | 53-1.3 (only water-mediated) | 17-0.5 (only water-mediated) | 54-1.2 (only water-mediated) | 19-0.6 (only water-mediated) | No | No |
| | E10-R25 | 74-0.6 OE1/OE2-NH1/NH2 | 36-0.2 OE1/OE2-NH1/NH2 | 80-0.5 OE1/OE2-NE/NH1/NH2 | 45-0.1 OE1-NH1/NH2, OE2-NE/NH1/NH2 | 25-1.2 (only water-mediated) | No |
| | D11-R25 | No | No | No | No | No | 16-0.2 OD1/OD2-NH1/NH2 |
| | E15-D16 | 33-2.3 (only water-mediated) | No | 28-2.0 (only water-mediated) | No | 47-2.1 (only water-mediated) | No |
| | E15-R19 | 67-0.9 OE1/OE2-NE/NH1/NH2 | 23-0.4 (only water-mediated) | 51-1.0 OE1/OE2-NE/NH1/NH2 | 18-0.4 OE1/OE2-NE/NH1/NH2 | 70-0.6 OE1/OE2-NH1/NH2 | 30-0.2 OE1/OE2-NH1/NH2 |
| | D16-R19 | 54-1.1 OD1-NE/NH1/NH2, OD2-NH1/NH2 | 14-0.4 (only water-mediated) | 81-0.3 OD1/OD2-NE/NH1/NH2 | 40-0.1 OD1/OD2-NH1/NH2 | 95-0.1 OD1/OD2-NE/NH1/NH2 | 57-0.1 OD1/OD2-NE/NH1/NH2 |
| | H26-S22 | 34-1.9 (only water-mediated) | No | 26-2.0 (only water-mediated) | No | No | No |
| | H26-N29 | 33-1.6 (only water-mediated) | No | No | No | 30-1.8 (only water-mediated) | No |
| pH5 | D6-S3 | 54-1.1 OD1/OD2-OG | 20-0.3 OD1/OD2-OG | 67-0.7 OD1/OD2-OG | 31-0.2 OD1/OD2-OG | 45-0.6 OD1/OD2-OG | 25-0.1 OD1/OD2-OG |
| | D6-Y20 | 34-2.0 (only water-mediated) | No | 34-1.8 (only water-mediated) | No | No | No |
| | E10-D11 | 53-1.8 (only water-mediated) | 10-1.5 (only water-mediated) | 45-1.6 (only water-mediated) | No | 31-1.9 (only water-mediated) | No |

| | | | | | | | |
|---|---|---|---|---|---|---|---|
| | E10-Y21 | 58-1.3 OE1/OE2-OH | 18-0.6 (only water-mediated) | 53-1.5 (only water-mediated) | 14-0.8 (only water-mediated) | No | No |
| | E10-R25 | 47-1.2 OE1/OE2-NH1/NH2 | 17-0.3 OE1/OE2-NH1/NH2 | 27-2.1 (only water-mediated) | No | 65-0.4 OE1/OE2-NE/NH1/NH2 | 34-0.1 OE1/OE2-NH1/NH2 |
| | D11-R25 | No | No | No | No | No | 13-0.1 OD1/OD2-NH1/NH2 |
| | E15-D16 | 34-2.0 (only water-mediated) | No | 30-2.2 (only water-mediated) | No | 28-2.1 (only water-mediated) | No |
| | E15-R19 | 56-0.7 OE1/OE2-NE/NH1/NH2 | 26-0.2 OE1/OE2-NE/NH1/NH2 | 55-1.0 OE1/OE2-NE/NH1/NH2 | 16-0.4 OE1/OE2-NE/NH1/NH2 | 57-0.8 OE1/OE2-NE/NH1/NH2 | 21-0.3 OE1/OE2-NE/NH1/NH2 |
| | D16-R19 | 78-0.5 OD1/OD2-NE/NH1/NH2 | 37-0.2 OD1-NH1/NH2, OD1/OD2-NE/NH1/NH2 | 77-0.3 OD1/OD2-NE/NH1/NH2 | 40-0.1 OD1-NH1/NH2, OD1/OD2-NE/NH1/NH2 | 69-0.4 OD1/OD2-NE/NH1/NH2 | 32-0.1 OD1/OD2-NE/NH1/NH2 |
| | H26-S22 | 31-2.0 (only water-mediated) | No | No | No | No | No |
| | H26-N29 | 33-1.6 (only water-mediated) | No | No | No | 27-1.8 (only water-mediated) | No |
| pH4 | D6-S3 | 45-1.2 OD1/OD2-OG | 14-0.3 OD1/OD2-OG | 48-1.4 OD1/OD2-OG | 12-0.4 (only water-mediated) | 33-1.6 (only water-mediated) | No |
| | E10-D11 | 42-1.8 (only water-mediated) | No | 47-1.8(only water-mediated) | No | 37-1.8 (only water-mediated) | No |
| | E10-Y21 | 54-1.0 OE1/OE2-OH | 19-0.4 OE1/OE2-OH | 51-1.5 (only water-mediated) | 14-0.8 (only water-mediated) | 42-1.4 (only water-mediated) | 12-0.6 (only water-mediated) |
| | E10-R25 | 46-1.5 (only water-mediated) | 11-0.7 (only water-mediated) | 37-1.8 (only water-mediated) | No | 45-0.9 OE1/OE2-NH1/NH2 | 21-0.2 OE1/OE2-NH1/NH2 |

| | | | | | | | |
|---|---|---|---|---|---|---|---|
| | E15-D16 | 28-2.1 (only water-mediated) | No | 26-2.1 (only water-mediated) | No | 25-2.2 (only water-mediated) | No |
| | E15-R19 | 50-1.1 OE1/OE2-NE/NH1/NH2 | 15-0.4 OE1/OE2-NE/NH1/NH2 | 38-1.4 OE1/OE2-NE/NH1/NH2 | No | 34-1.4 (only water-mediated) | No |
| | D16-R19 | 77-0.4 OD1/OD2-NE/NH1/NH2 | 32-0.2 OD1/OD2-NE/NH1/NH2 | 78-0.3 OD1/OD2-NE/NH1/NH2 | 37-0.1 OD1/OD2-NE/NH1/NH2 | 80-0.3 OD1/OD2-NE/NH1/NH2 | 42-0.1 OD1/OD2-NE/NH1/NH2 |
| | H26-S22 | No | No | 35-1.9 (only water-mediated) | No | No | No |
| | H26-N29 | No | No | 29-1.7 (only water-mediated) | No | 25-1.6 (only water-mediated) | No |
| pH3 | D6-S3 | No | No | No | No | 40-1.5 | No |
| | E10-D11 | 31-2.0 (only water-mediated) | No | 35-1.9 (only water-mediated) | No | 37-2.0 (only water-mediated) | No |
| | E10-Y21 | No | No | 30-1.6 (only water-mediated) | No | No | No |
| | E10-R25 | 28-1.2 (only water-mediated) | No | 53-0.8 OE1/OE2-NH1/NH2 | 24-0.1 OE1/OE2-NH1/NH2 | No | No |
| | E15-R19 | 49-1.1 OE1/OE2-NE/NH1/NH2 | 11-0.4 (only water-mediated) | 31-1.7 (only water-mediated) | No | 43-1.5 (only water-mediated) | No |
| | D16-R19 | 65-0.7 OD1/OD2-NE/NH1/NH2 | 21-0.3 OD1/OD2-NH1/NH2 | 47-1.2 OD1/OD2-NE/NH1/NH2 | 10-0.6 OD1/OD2-NH1/NH2 | 68-0.7 OD1/OD2-NE/NH1/NH2 | 28-0.2 OD1/OD2-NE/NH1/NH2 |
| | H26-S22 | No | No | 33-1.8 (only water-mediated) | No | No | No |
| | H26-N29 | No | No | 27-1.7 | No | No | No |

| | | | | |
|---|---|---|---|---|
| | | | | (only water-mediated) |

**S4 Table.** Same as **S3** but only for direct H-bonds and the occupancy of 15% for the criterion of $60^0$ 00 and 10% for the criterion of $20^0$.

| pH | H-bond criterion | R#1 | | R#2 | | R#3 | |
|---|---|---|---|---|---|---|---|
| | | $60^0$ | $20^0$ | $60^0$ | $20^0$ | $60^0$ | $20^0$ |
| pH7 | D6-S3 | 30<br>OD1/OD2-OG | 21<br>OD1/OD2-OG | 52<br>OD1/OD2-OG | 37<br>OD1/OD2-OG | 49<br>OD1/OD2-OG | 32<br>OD1/OD2-OG |
| | E10-Y21 | No | No | No | No | 19<br>OE1/OE2-OH | 12<br>OE1/OE2-OH |
| | E10-R25 | No | No | No | No | 43<br>OE1/OE2-NH1/NH2 | 33<br>OE1/OE2-NH1/NH2 |
| | D11-K4 | 20<br>OD2-NZ | No | No | No | No | No |
| | D11-R25 | No | No | 61<br>OD1/OD2-NH1/NH2 | 51<br>OD1/OD2-NH1/NH2 | No | No |
| | E15-R19 | 38<br>OE1/OE2-NE/NH1/NH2 | 21<br>OE1/OE2-NE/NH1/NH2 | 34<br>OE1/OE2-NE/NH1/NH2 | 17<br>OE1/OE2-NE/NH1/NH2 | 63<br>OE1/OE2-NE/NH1/NH2 | 35<br>OE1/OE2-NH1/NH2 |
| | D16-R19 | 45<br>OD1/OD2-NE/NH1/NH2 | 24<br>OD1-NE/NH1/NH2, OD2-NH1/NH2 | 51<br>OD1/OD2-NE/NH1/NH2 | 28<br>OD1/OD2-NH1/NH2 | 54<br>OD1/OD2-NE/NH1/NH2 | 23<br>OD1/OD2-NH1/NH2 |
| pH6 | D6-S3 | 20<br>OD1/OD2-OG | 14<br>OD1/OD2-OG | 67<br>OD1/OD2-OG | 47<br>OD1/OD2-OG | 69<br>OD1/OD2-OG | 41<br>OD1/OD2-OG |
| | E10-Y21 | 16<br>OE1/OE2-OH | 12<br>OE1/OE2-OH | No | No | 15<br>OE1/OE2-OH | 10<br>OE1/OE2-OH |
| | E10-R25 | 48<br>OE1/OE2-NH1/NH2 | 32<br>OE1/OE2-NH1/NH2 | No | No | 59<br>OE1/OE2-NE/NH1/NH2 | 42<br>OE1-NH1/NH2, OE2-NE/NH1/NH2 |
| | D11-R25 | No | No | 18<br>OD1/OD2-NH1/NH2 | 14<br>OD1/OD2-NH1/NH2 | No | No |
| | E15-R19 | 34<br>OE1/OE2-NE/NH1/NH2 | No | 23<br>OE1/OE2-NE/NH1/NH2 | 13<br>OE1/OE2-NE/NH1/NH2 | 44<br>OE1/OE2-NH1/NH2 | 25<br>OE1/OE2-NH1/NH2occu |

| | | | | | | | |
|---|---|---|---|---|---|---|---|
| | D16-R19 | 24 OD1-NE/NH1/NH2, OD2-NH1/NH2 | No | 67 OD1/OD2-NE/NH1/NH2 | 35 OD1/OD2-NE/NH1/NH2 | 89 OD1/OD2-NE/NH1/NH2 | 53 OD1/OD2-NH1/NH2 |
| pH5 | D6-S3 | 23 OD1/OD2-OG | 15 OD1/OD2-OG | 33 OD1/OD2-OG | 23 OD1/OD2-OG | 42 OD1/OD2-OG | 26 OD1/OD2-OG |
| | E10-Y21 | 16 OE1/OE2-OH | No | No | No | No | No |
| | E10-R25 | 19 OE1/OE2-NH1/NH2 | 14 OE1/OE2-NH1/NH2 | 51 OE1/OE2-NE/NH1/NH2 | 32 OE1/OE2-NH1/NH2 | No | No |
| | D11-R25 | No | No | No | 13 OD1/OD2-NH1/NH2 | No | No |
| | E15-R19 | 36 OE1/OE2-NE/NH1/NH2 | 22 OE1/OE2-NE/NH1/NH2 | 32 OE1/OE2-NE/NH1/NH2 | 17 OE1/OE2-NE/NH1/NH2 | 24 OE1/OE2-NE/NH1/NH2 | 11 OE1/OE2-NE/NH1/NH2 |
| | D16-R19 | 55 OD1/OD2-NE/NH1/NH2 | 31 OD1-NH1/NH2, OD2-NE/NH1/NH2 | 54 OD1/OD2-NE/NH1/NH2 | 29 OD1/OD2-NE/NH1/NH2 | 66 OD1/OD2-NE/NH1/NH2 | 37 OD1-NH1/NH2, OD2-NE/NH1/NH2 |
| pH4 | D6-S3 | 20 OD1/OD2-OG | 11 OD1/OD2-OG | No | No | 16 OD1/OD2-OG | No |
| | E10-Y21 | 23 OE1/OE2-OH | 14 OE1/OE2-OH | No | No | No | No |
| | E10-R25 | No | No | 24 OE1/OE2-NH1/NH2 | 18 OE1/OE2-NH1/NH2 | No | No |
| | E15-R19 | 20 OE1/OE2-NE/NH1/NH2 | 10 OE1/OE2-NE/NH1/NH2 | No | No | No | No |
| | D16-R19 | 55 OD1/OD2-NE/NH1/NH2 | 27 OD1/OD2-NE/NH1/NH2 | 68 OD1/OD2-NE/NH1/NH2 | 38 OD1/OD2-NE/NH1/NH2 | 65 OD1/OD2-NE/NH1/NH2 | 34 OD1/OD2-NE/NH1/NH2 |
| pH3 | E10-R25 | No | No | No | No | 30 OE1/OE2-NH1/NH2 | 21 OE1/OE2-NH1/NH2 |
| | E15-R19 | 20 OE1/OE2-NE/NH1/NH2 | No | No | No | No | No |
| | D16-R19 | 39 OD1/OD2-NE/NH1/NH2 | 17 OD1/OD2-NH1/NH2 | 16 OD1/OD2-NH1/NH2 | No | 40 OD1/OD2-NE/NH1/NH2 | 24 OD1/OD2-NE/NH1/NH2 |

**S5 Table**. Helical secondary structure content in the last 110 of standard MD simulations (R#1-3) and for the last 40 ns (R#1and R#2) and 30 ns (R#3) of the constant pHD simulations.

| Type of simulations | Residues of the α-helical segment |
|---|---|

| | | Always within the α-helix | Transient extensions<br>(that is, at times during the dynamics) |
|---|---|---|---|
| *Standard MD* | | | |
| R#1 | | Glu15-Ile31 | Thr32-Arg35 |
| R#2 | | Ala14-Gln34 | none |
| R#3 | | Ala14-Arg33 | Gln34-Arg35 |
| *Constant pH MD* | | | |
| pH = 7 | R#1 | Glu15-Ile31 | Thr32 |
| | R#2 | Glu15-Ile31 | Thr32 |
| | R#3 | Glu15-Thr32 | none |
| pH = 6 | R#1 | Glu15-Thr32 | Arg33-Arg35 |
| | R#2 | Glu15-Ile31 | Thr32 |
| | R#3 | Glu15-Thr32 | Arg33-Gln34 |
| pH = 5 | R#1 | Glu15-Arg33 | Gln34 |
| | R#2 | Glu15-Ile31 | Thr32 |
| | R#3 | Glu15-Ile31 | Thr32 |
| pH = 4 | R#1 | Glu15-Leu30 | Ile31 |
| | R#2 | Glu15-Ile31 | None |
| | R#3 | Glu15-Gln34 | Arg35 |
| pH = 3 | R#1 | Glu15-Thr32 | Arg33-Gln34 |
| | R#2 | Glu15-Ile31 | Thr32-Arg33 |
| | R#3 | Glu15-Thr32 | Arg33-Arg35 |
